# Conditional Chemoconnectomics (cCCTomics): Conditional Targeting of Chemical Transmission Efficiently

**DOI:** 10.1101/2023.09.26.559642

**Authors:** Renbo Mao, Jianjun Yu, Bowen Deng, Xihuimin Dai, Yuyao Du, Sujie Du, Wenxia Zhang, Yi Rao

## Abstract

Dissection of neural circuitry underlying behaviors is a central theme in neurobiology. We have previously proposed the concept of chemoconnectome (CCT) to cover the entire chemical transmission between neurons and target cells in an organize and created tools for studying it (CCTomics) by targeting all genes related to the CCT in Drosophila. Here we have created lines targeting the CCT in conditional manners after modifying GFP RNA interference, Flp-out and CRISPR/Cas9 technologies. All three strategies are validated to be highly effective with the best using chromatin-peptide fused Cas9 variants and scaffold optimized sgRNAs. As a proof of principle, we conduct a comprehensive intersection analysis of CCT genes expression profiles in the clock neurons, uncovering 43 CCT genes present in clock neurons. Specific elimination of each from clock neurons revealed that loss of the neuropeptide CNMa in two posterior dorsal clock neurons (DN1ps) or its receptor (CNMaR) caused advanced morning activity, indicating a suppressive role of CNMa-CNMaR on morning anticipation, opposite to the promoting role of PDF-PDFR on morning anticipation. These results demonstrate the effectiveness of conditional CCTomics and its tools created by us here and establish an antagonistic relationship between CNMa-CNMaR and PDF-PDFR signaling in regulating morning anticipation.

## Introduction

Much research efforts have been made to uncover the wiring and signaling pathways of neural circuits underlying specific behaviors. Circuit dissection strategies include genetic screening, genetic labelling, circuit tracing, live imaging, genetic sensors and central nervous system (CNS) reconstruction via electron microscopy (EM). Recently, we have developed chemoconnectomics (CCTomics), focusing on building a comprehensive set of knockout and knockin tool lines of chemoconnectome (CCT) genes, to dissect neural circuitry based on chemical transmission (Deng et al., 2019).

Each strategy has advantages and disadvantages. For example, genetic screening is less biased but inefficient; circuit tracing with viruses provides information of connection, but is often prone to leaky expression and inaccurate labelling; and EM reconstruction is anatomically accurate but does not allow for manipulation of corresponding neurons. CCTomics overcomes limitations of previous strategies by allowing for behavioral screening of CCT genes and accurate labelling or manipulation of corresponding neurons. However, it is still limited in that knockout of some CCT genes can be lethal during development and that CCT genes may function differently in different neurons, which require a cell type-specific manipulation. Thus, we decided to invent a conditional CCTomics (cCCTomics) in which gene deletion was conditional.

There are three major strategies for somatic gene mutagenesis at the DNA/RNA level: RNA interference, DNA site-specific recombination enzymes, and CRISPR/Cas system. RNA interference targets RNAs conveniently and efficiently (Martin and Caplen, 2007; Oberdoerffer et al., 2005). Libraries of transgenic RNAi flies covering almost the entire fly genome have been established (Ni et al., 2011; Perkins et al., 2015). DNA site specific recombination enzymes such as Flp, B3 and Cre mediate specific and efficient gene editing (Gaj et al., 2014; Grindley et al., 2006). These strategies require flies with reverse repetitive sequences knocked into the corresponding genes, which is time-consuming with relatively complex recombination for genetic assays. CRISPR/Cas systems, particularly CRISPR/Cas9, which targets DNA with a sgRNA/Cas protein complex, have been broadly applied in gene manipulation over the last decade. The widespread use of CRIPSR/Cas9 in Drosophila somatic gene manipulation began in 2014 (Xue et al., 2014). Later, tRNA-flanking sgRNAs was designed and applied, which enabled multiple sgRNAs to mature in a single transcript (Xie et al., 2015), accelerating the application of this strategy in conditional gene manipulation in flies with impressive efficiency (Delventhal et al., 2019; Port and Bullock, 2016; Port et al., 2020; Schlichting et al., 2019; Schlichting et al., 2022). Additionally, libraries of UAS-sgRNA targeting kinases (Port et al., 2020) and GPCRs (Schlichting et al., 2022) have been established, but no sgRNA libraries covering all the CCT genes exists yet. The efficiency of CRISPR/Cas9 has not been validated systematically in the Drosophila nervous system.

The circadian rhythm can be used for proof of principle testing of cCCTomics. Organisms evolve periodic behaviors and physiological traits in response to cyclical environmental changes. The rhythmic locomotor behavior of Drosophila, for instance, shows enhanced activity before the light is turned on and off in a light-dark (LD) cycle, referred to as morning and evening anticipations, respectively (Collins et al., 2005; Helfrich-Förster, 2001). Under 12 hours (h) dark-12 h dark (DD) conditions, the activities peaks regularly about every 24 h (Konopka and Benzer, 1971). Approximately 150 clock neurons, circadian output neurons and extra-clock electrical oscillators (xCEOs) coordinate Drosophila circadian behaviors (Dubowy and Sehgal, 2017; Tang et al., 2022). The regulation of morning and evening anticipations, the most prominent features in the LD condition, is primarily mediated by four pairs of sLNvs expressing pigment dispersing factor (PDF), six pairs of LNds and the 5^th^ s-LNv (Grima et al., 2004; Rieger et al., 2006; Stoleru et al., 2004). At the molecular level, Pdf and Pdf receptor (PDFR) are well-known, with their mutants showing an advanced evening activity peak and no morning anticipation (Hyun et al., 2005; Lear et al., 2005; Renn et al., 1999). Other neuropeptides and their receptors, including AstC/AstC-R2 and neuropeptide F (NPF) and its receptor (NPFR), have also been reported to modulate evening activities (Díaz et al., 2019; He et al., 2013; Hermann et al., 2012), while CCHa1/CCHa1-R and Dh31 regulate morning activities (Fujiwara et al., 2018; Goda et al., 2019). To date, no advanced morning activity phenotype has been reported in flies.

To develop a more efficient approach for somatic gene manipulation, we have now generated two systems for conditional manipulation of CCT genes. One is GFPi/Flp-out-based conditional knockout system of CCT genes (cCCTomics) and another is CRISPR/Cas9-based (C-cCCTomics). Both systems have achieved high efficiency of gene mutagenesis in the Drosophila nervous system. C-cCCTomics, utilizing chromatin-peptide fused Cas9 and scaffold optimized sgRNA, makes efficient conditional gene knockout as simple as RNAi. Further application of C-cCCTomics in clock neurons revealed novel roles of CCT genes in circadian behavior: CNMa-CNMaR modulates morning anticipation as an antagonistic signal of PDF-PDFR.

## Results

### Near Complete Disruption of Target Genes by GFPi and Flp-out Based cCCTomics

For the purpose of conditional chemoconnectomics (cCCTomics), we initially leveraged the benefits of our previously generated CCTomics attP lines (Deng et al., 2019), which enabled us to fuse enhanced GFP coding sequence at the 3’ end of each gene’s coding region and flank most or entire gene span with FRT sequence through site-specific recombination (Fig. 1A, Table S1). We designed this system so that it could be used to target genes tagged with GFP by RNAi (Neumüller et al., 2012) and at the same time to enable flippase(Flp) mediated DNA fragment excision between two FRT sequences when FRT sequences are in the same orientation (Vetter et al., 1983) (Fig. 1A).

**Fig. 1.**
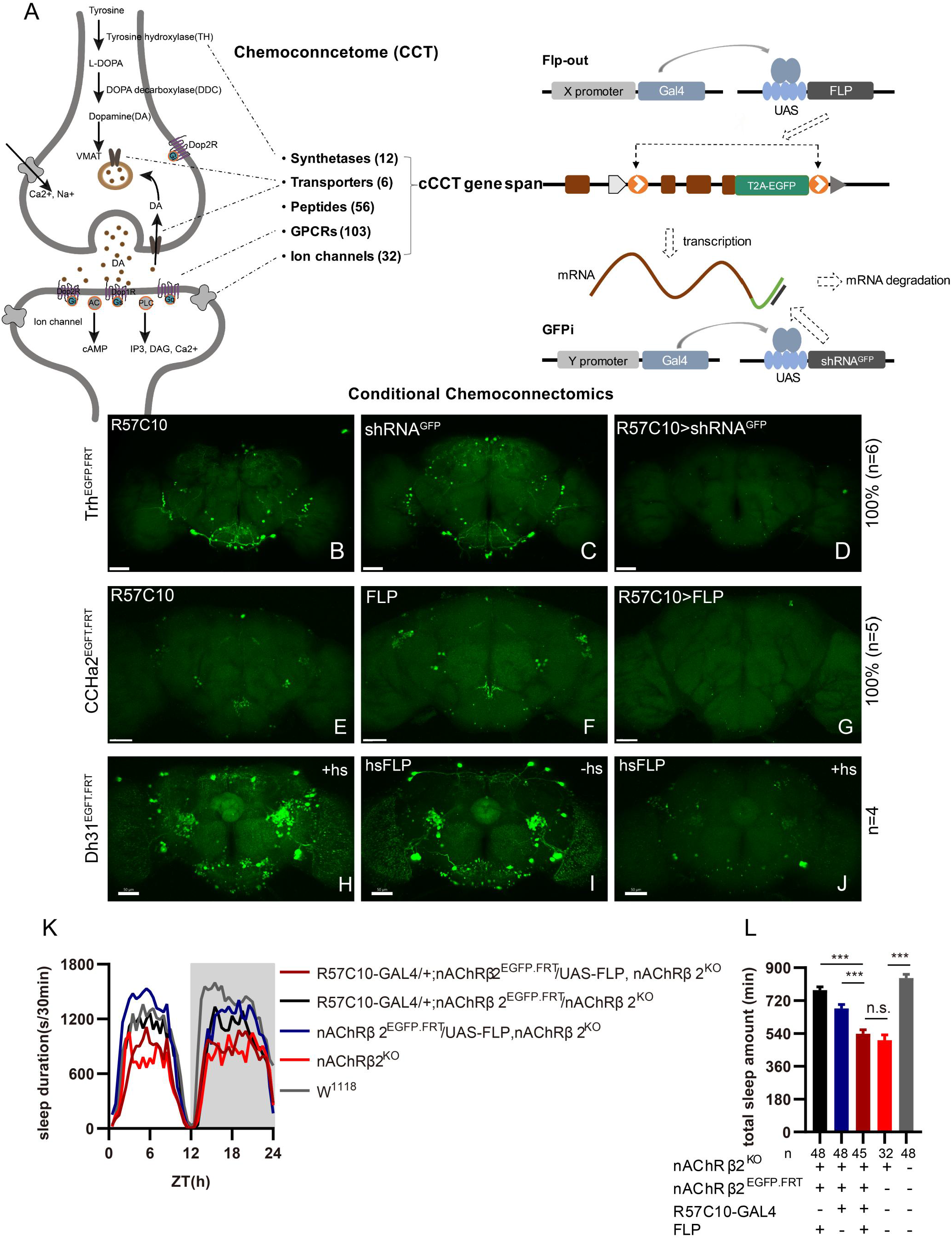
cCCTomics Mediates Efficient Conditional Disruption of CCT Genes. (A) Schematic of cCCT gene span and principle of cCCTomics. A T2A-EGFP sequence was introduced at the 3’ end of CCT genes and their most or all coding regions (depending on attP-KO lines) were flanked by 34 bp FRT sequence. Both Flp-out (top) and GFP RNAi (down) could mediate CCT gene manipulation. (B-J) Expression of Trh (B-D), CCHa2 (E-G), and Dh31 (H-J) are efficiently disrupted by pan-neuronal expression of GFP-RNAi (B-D), pan-neuronal expression of Flp-out (E-G), and heatshock-Flp (H-J) respectively. Representative fluorescence images of R57C10-Gal4/+;Trh^EGFP.FRT^/+ (B), UAS-shRNA^GFP^/Trh^EGFP.FRT^ (C), R57C10-Gal4/+; UAS-shRNA^GFP^/Trh^EGFP.FRT^ (D), R57C10-Gal4/+;CCHa2^EGFP.FRT^/+; (E), UAS-Flp/CCHa2^EGFP.FRT^ (F), R57C10-Gal4/+; UAS-Flp/CCHa2^EGFP.FRT^ (G), Dh31^EGFP.FRT^ with heatshock (H), hs-Flp/Dh31^EGFP.FRT^ without heatshock (I), and hs-Flp/Dh31^EGFP.FRT^ with heatshock are shown. Manipulation efficiency and experiment group fly number is noted on the right. Scale bar, 50um. (K-L) sleep profiles (K) and statistical analysis (L) of Flp-out induced nAChRβ2 neuronal knockout flies (dark red), nAChRβ2 knockout flies, and genotype controls (dark and blue). Sleep profiles are plotted in 30 min bins. In this and other figures, blank background indicates the light phase (ZT 0-12); shaded background indicates the dark phase (ZT 12-24). Daily sleep duration were significantly reduced in nAChRβ2 neuronal knockout files which is comparable to nAChRβ2 knockout. In all statistical panels, unless otherwise noted, 1) Numbers below each bar represent the number of flies tested. 2) Mean ± SEM is shown. 3) The Kruskal-Wallis test followed by Dunn’s post test was used. \*\*\**p<0.001*. ** *p<0.01*. * *p<0.05*. n.s. *p>0.05.* Male flies were used unless otherwise noted.

To validate the efficiency of cCCTomics, we performed pan-neuronal expression of either shRNA^GFP^ or flipase in cCCT flies. Immunofluorescent imaging showed that constitutive expression of shRNA^GFP^ (Fig. 1B-1D, Fig. S1A-S1I) or flipase (Fig. 1E-1G, Fig. S1J-S1U) almost completely eliminated GFP signals of target genes, indicating high efficiency. Knocking out at the adult stage using either hsFLP driven Flp-out (Golic and Lindquist, 1989) (Fig. 1H-1J) or neural (elav-Switch) driven shRNA^GFP^ (Nicholson et al., 2008; Osterwalder et al., 2001) (Fig. S2A-S2I), also resulted in the elimination of most, though not all, GFP signals. Notably, control group of CCT^EGFP.FRT^; elav-Switch/UAS-shRNAGFP flies fed with solvent (ethanol) showed obvious decreased GFP (Fig. S2B, S2E, and S2H) comparing with UAS-shRNA^GFP^/CCT^EGFP.FRT^ flies fed with RU486 (Fig. S2A, S2D, and S2G), indicating leaky expression of elav-Switch.

We then applied cCCTomics in pan-neuronal knockout of nAChRβ2 which is required for Drosophila sleep (Dai et al., 2021). Ablation of nAChRβ2 in the nervous system dramatically decreased sleep of flies, mirroring the nAChRβ2 knockout phenotype (Fig. 1K, 1L). Therefore, cCCTomics is an effective toolkit for manipulation of CCT genes and suitable for functional investigations of genes. Expression of in-frame fused eGFP-labelled CCT genes highly co-localized with signals revealed by immunocytochemistry (Fig. S3A-S3I), allowing direct examination of gene expression without amplification, which is different from the GAL4/UAS binary system.

We then checked the viability of cCCT lines and found that cCCT lines including Capa^EGFP.FRT^, ChAT^EGFP.FRT^ and Eh^EGFP.FRT^ were viable whereas their CCT mutants were lethal. Gad1^EGFP.FRT^, GluRIID^EGFP.FRT^ and CapaR^EGFP.FRT^ were still lethal as their CCT mutants were (Table S2). This indicates that some of the cCCT knockin flies may functionally affect corresponding genes, which are not suitable for conditional gene manipulation. Combination of cCCT transgenic flies with UAS-flp, UAS-shRNA^GFP^ or specific drivers is complicated and unfriendly for screen work, despite the almost 100% efficiency of gene suppression when driven by a pan-neuronal driver. Because of the limitations of this method, we further created a CRISPR/Cas9 based conditional knockout system of chemoconnectomics (C-cCCTomics).

### CRISPR/Cas9-Based Conditional Knockout System for CCTomics

To simplify effective manipulation of CCT genes, we designed a vector based on pACU2 (Han et al., 2011) with tRNA flanking sgRNAs (Port and Bullock, 2016; Xie et al., 2015) targeting CCT genes (Fig. 2A). We also adopted an optimized sgRNA scaffold “E+F” (E, stem extension; F, A-U flip) (Chen et al., 2013), which facilitates Cas9-sgRNA complex formation and gene knockout efficiency (Dang et al., 2015; Poe et al., 2019; Zhao et al., 2016), to all sgRNAs to improve gene knockout efficiency. To balance efficiency and off-target effect, we selected three sgRNAs for each CCT gene with highest predicted efficiency and no predicted off-target effect based on previously reported models (Chu et al., 2016; Doench et al., 2014; Graf et al., 2019; Gratz et al., 2014; Heigwer et al., 2014; Stemmer et al., 2015; Xu et al., 2015) (see Table S3 and details at Methods).

**Fig. 2.**
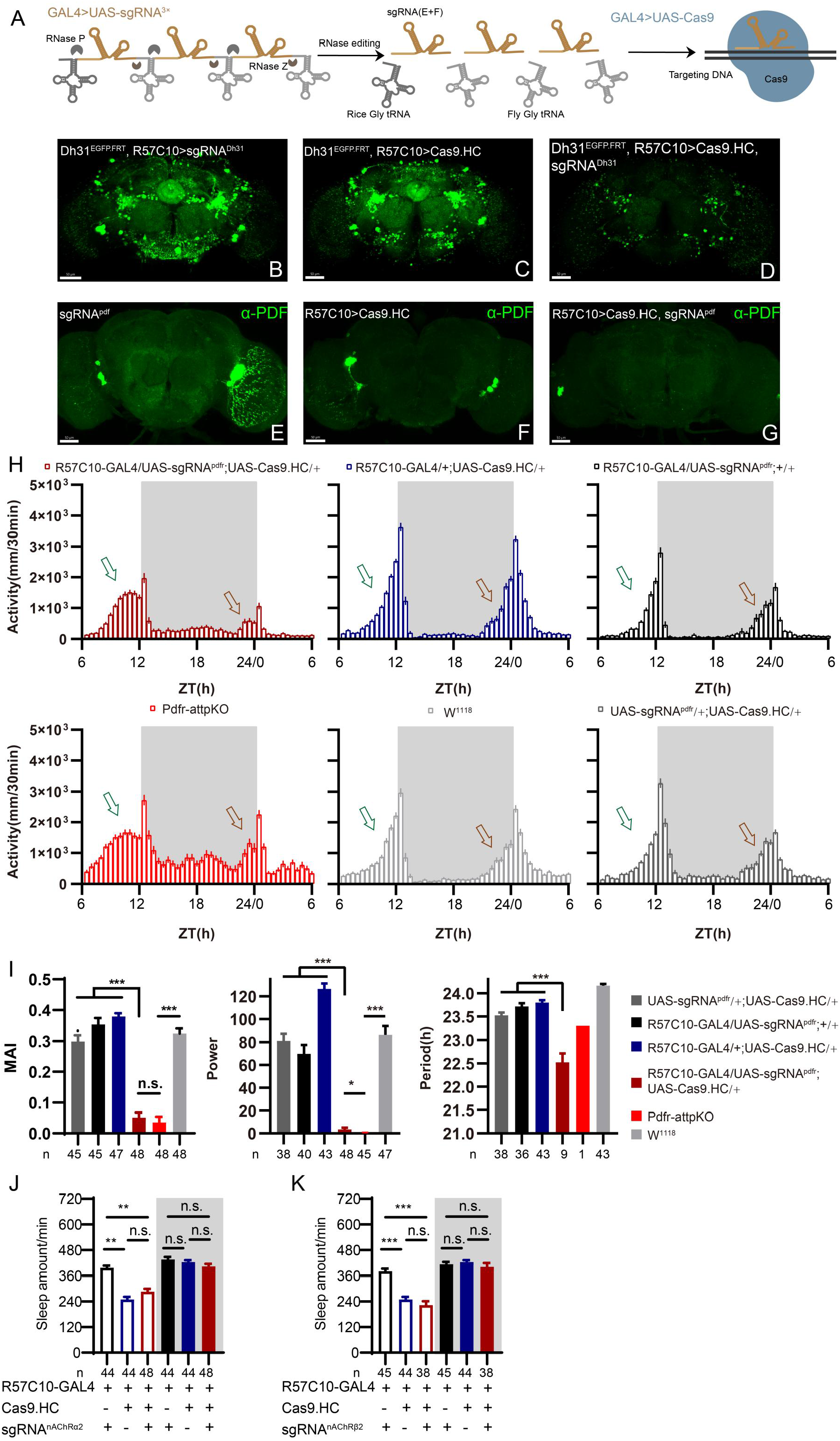
C-cCCTomics Mediates Efficient Conditional Knockout of CCT Genes. (A) Schematic of C-cCCTomics principle. Cas9 and three sgRNAs are driven by GAL4/UAS system. Three tandem sgRNAs are segregated by fly tRNA^Gly^ and matured by RNase Z and RNase P. (B-G) Pan-neuronal knockout of Dh31 (D) and Pdf (G) by C-cCCTomics strategy. Representative fluorescence images presented expression of Dh31 (B-D) or anti-PDF (E-G). Pan-neuronal expression of Cas9 and sgRNA eliminated most (D) or all (G) fluorescent signal compared to control fly brains (B-C, E-F). Scale bar, 50 μm. (H) Activity profiles of pan-neuronal knockout of Pdfr and Pdfr-attpKO. Activity profiles were centered of the 12h darkness in all figures with evening activity on the left and morning activity on the right, which is different from general circadian literatures. Plotted in 30 min bins. (I) Statistical analysis of morning anticipation index (MAI), power, and period for pan-neuronal Pdf knockout and Pdfr-attpKO flies. Knocking out of Pdfr in neurons reduced both morning anticipation index, power and period significantly. (J-K) Statistical analysis of nAChRα2 (J) and nAChRβ2 (K) pan-neuronal knockout flies’ sleep phenotype. Sleep of these flies were not disrupted.

We generated UAS-sgRNA^3x^ transgenic lines for all 209 defined CCT genes (Abruzzi et al., 2017; Dai et al., 2019; Deng et al., 2019) and UAS-Cas9.HC (Cas9.HC) (Mali et al., 2013). We first verified that C-cCCTomics mediated precise target DNA breaking by ubiquitous expression of Cas9.P2 (Port et al., 2014) and sgRNA by targeting Pdf or Dh31. Sanger sequencing showed that indels were present exactly at the Cas9 cleavage sites (Fig. S4A-S4H).

To determine the efficiency of C-cCCTomics, we employed pan-neuronal expression of Cas9.HC with sgRNA^Dh31^ or sgRNA^pdf^. Targeting by Cas9.HC/sgRNA^Dh31^ eliminated most but not all of the GFP signal in Dh31^EGFP.FRT^ (Fig 2B-2D), whereas all anti-PDF signals were eliminated by Cas9.HC/sgRNA^pdf^ (Fig 2E-2G). Furthermore, we used the C-cCCT strategy to conditionally knockout genes for pdfr, nAChRβ2 and nAChRα2, which were previously reported as essential for circadian rhythm or sleep (Dai et al., 2021; Renn et al., 1999). Pan-neuronal knockout of Pdfr resulted in a tendency towards advanced evening activity and weaker morning anticipation compared to control flies (Fig. 2H-2I), which is similar to Pdfr-attpKO flies. These phenotypes were not as pronounced as those reported previously, when han^5304^ mutants exhibited a more obvious advanced evening peak and no morning anticipation (Hyun et al., 2005). Furthermore, there was no significant sleep decrease in these conditional knockout (cKO) flies (Fig. 2J-2K) when we applied C-cCCTomics to manipulate nAChRβ2 or nAChRa2. Taken together, C-cCCTomics (with Cas9.HC) achieved a relatively high gene knockout efficiency, but it was not effective enough for all genes.

### Evaluation of Cas9 with Different Chromatin-Modulating Peptides

Since the establishment of the CRISPR/Cas9 system a decade ago, many groups have attempted to improve its efficiency in gene manipulation. Most attempts have been focused on the two main components of this system, the Cas9 protein (Ding et al., 2019; Ling et al., 2020; Liu et al., 2019; Zhao et al., 2016; Zheng et al., 2020) and the single guide RNA (Chen et al., 2013; Chu et al., 2016; Dang et al., 2015; Doench et al., 2014; Filippova et al., 2019; Graf et al., 2019; Labuhn et al., 2018; Mu et al., 2019; Nahar et al., 2018; Scott et al., 2019; Xu et al., 2015). At the beginning of C-cCCTomics design, we adopted an optimized sgRNA scaffold and selected sgRNAs with predicted high efficiency. We tried to further improve the efficacy by modifying Cas9 protein. We fused a chromatin-modulating peptide (Ding et al., 2019), HMGN1 (High mobility group nucleosome binding domain 1), at the N-terminus of Cas9 and HMGB1 (High mobility group protein B1) at its C-terminus with GGSGP linker, termed Cas9.M9 (Fig. 3A, Methods). We also obtained a modified Cas9.M6 with HMGN1 at the N-terminus and an undefined peptide (UDP) at the C-terminus (Fig. 3A). We replaced the STARD linker between Cas9 and NLS in Cas9.HC with GGSGP the linker (Zhao et al., 2016), termed Cas9.M0 (Fig 3A). None of these modifications have been validated previously in flies.

**Fig. 3.**
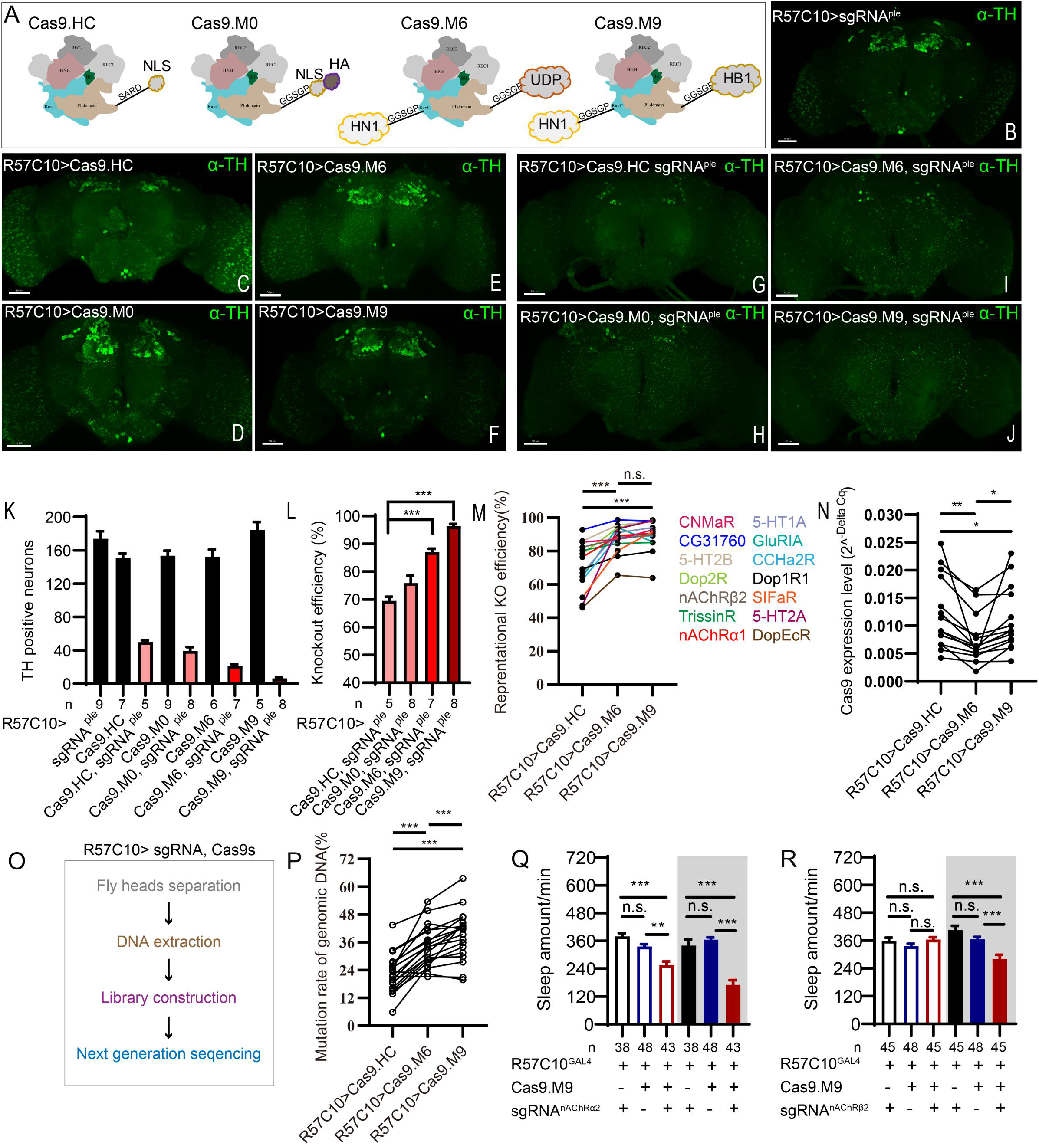
Efficiency Evaluation of Variations of Chromatin-Modulating Peptides Modified Cas9. (A) Schematics of chromatin-modulating peptides modified Cas9. (B-J) Efficiency evaluation of Cas9 variants. Fluorescence imaging of R57C10-Gal4>UAS-sgRNA^ple^ (B), R57C10-Gal4>UAS-Cas9 (C-F), and R57C10-Gal4>UAS-Cas9, UAS-sgRNA^ple^ (G-J) flies are shown. Brains were stained with anti-TH (green). Scale bar is 50μm. (K) Anterior TH positive neurons numbers of (K-U). (L) Statistical analysis of ple knockout efficiency related to (K). Modified Cas9.M6 and Cas9.M9 showed an improved efficiency comparing to Cas9.HC. Student’s t test was used. (M) Statistical analysis of representational KO efficiency of Cas9 variants as related to Figure S5. Gene symbols on the right indicate tested genes. (N) Statistical analysis of Cas9 expression level. (O-P) Workflow of efficiency validation by next-generation sequencing (O) and Statistical analysis of single-site mutation ratios induced by Cas9 variants (P). Paired t test was used in (M), (N) and (P). (Q-R) Statistical analysis of sleep amount for nAChRα2 (Q) or nAChRβ2 (R) pan-neuronal knockout flies. Knockout of nAChRα2 and nAChRβ2 by modified Cas9.M9 significantly decreased flies’ sleep amount.

To determine whether the modified Cas9 variants were more efficient, we first pan-neuronally expressed each Cas9 variant and sgRNA^ple^, and assessed their efficiency by immunofluorescence imaging. By counting anti-TH positive neurons in the brain (anterior view) after targeting by Cas9/sgRNA^ple^, we found that unmodified Cas9.HC/sgRNA^ple^ only achieved 69.58±3.04% (n=5) knockout efficiency (Fig. 3G, 3K, 3L), while Cas9.M6/sgRNA^ple^ and Cas9.M9/sgRNA^ple^ significantly improved efficiency to 87.53±3.06% (n=7) and 97.19±2.15% (n=8), respectively (Fig. 3I-3L). Fourteen additional CCT genes were subjected to pan-neuronal knockout, and the mRNA levels of the target genes were evaluated using real-time quantitative PCR with at least one primer overlapping the sgRNA targeting site (Fig. S5). Cas9.M6 and Cas9.M9 demonstrated significantly higher gene disruption efficiency compared to the unmodified Cas9.HC, achieving average efficiencies of 87.51%±2.24% and 89.59%±2.39% for Cas9.M6 and Cas9.M9, respectively, in contrast to 70.72%±3.82% for Cas9.HC. (Fig. 3M, Fig. S5). To rule out the possibility of the observed variations in gene disruption efficiency being attributed to differential Cas9 expression levels, we quantified the Cas9 expression levels and noted that both Cas9.M6 and Cas9.M9 exhibited lower mRNA levels than Cas9.HC under the experiment condition (Fig. 3N). Subsequently, genomic DNA of Drosophila head was extracted, and libraries encompassing target sites were constructed for high-throughput sequencing to verify disparities in genetic editing efficiency among these three Cas9 variants (Fig. 3O). In almost all nineteen sites tested, the mutation ratio consistently showed a trend towards Cas9.M6 and Cas9.M9 having a higher gene disruption efficiency than Cas9.HC (Fig. 3P, Fig. S6). The single-site mutation rates varied from 5.81% to 43.47% for Cas9.HC, 22.40% to 53.54% for Cas9.M6, and 19.90% to 63.57% for Cas9.M9 (Fig. 3P, Fig. S6). It should be noted that genomic DNA extracted from fly heads contained glial cells, which did not express Cas9/sgRNA, leading to a larger denominator and consequently reducing the observed mutation rates. Unmodified Cas9 displayed mutation rates comparable to those previously reported by Schlichting et al., 2022. The findings indicated that both Cas9.M6 and Cas9.M9 displayed elevated efficiency compared to Cas9.HC, with Cas9.M9 exhibiting the highest mutagenesis proficiency. These results suggest that the implementation of modified C-cCCTomics using Cas9.M6 and Cas9.M9 achieved an elevated level of efficiency. While unmodified C-cCCTomics was not efficient enough to knockout nAChRβ2 and nAChRα2 to phenocopy their mutants, we employed Cas9.M9 in conditional knockout of these genes to verify its efficiency. Pan-neuronal knocking out of nAChRβ2 or nAChRa2 by Cas9.M9/sgRNA showed significant sleep decrease which was similar to their mutants (Fig. 3Q-3R) (Dai et al., 2021).

Taken together, our results support that we have created a high efficiency toolkit for CCT gene manipulation in the nervous system, as well as more efficient Cas9 variants, Cas9.M6 and Cas9.M9, which can also be applied to genes other than those in the CCT.

### 43 CCT Genes Detected in Clock Neurons by Genetic Intersection

We analyzed the expression profile of CCT genes in circadian neurons with CCTomics driver lines in all clock neurons expressing Clk856 (Gummadova et al., 2009). With the Flp-out or split-lexA intersection strategy (Fig. 4A, 4B), we found 43 out of 148 analyzed CCT genes expressed in circadian neurons (Fig. 4C, Fig S7-S8, Table S4, Table S5). In all eight subsets of clock neurons, 23 CCT genes were expressed in DN1s, 20 in DN2s, 22 in DN3s, 28 in LNds, 14 in l-LNvs, 12 in s-LNvs, 5 in 5^th^ s-LNv and 3 in LPNs, with a total of 127 gene-subsets.

**Fig 4.**
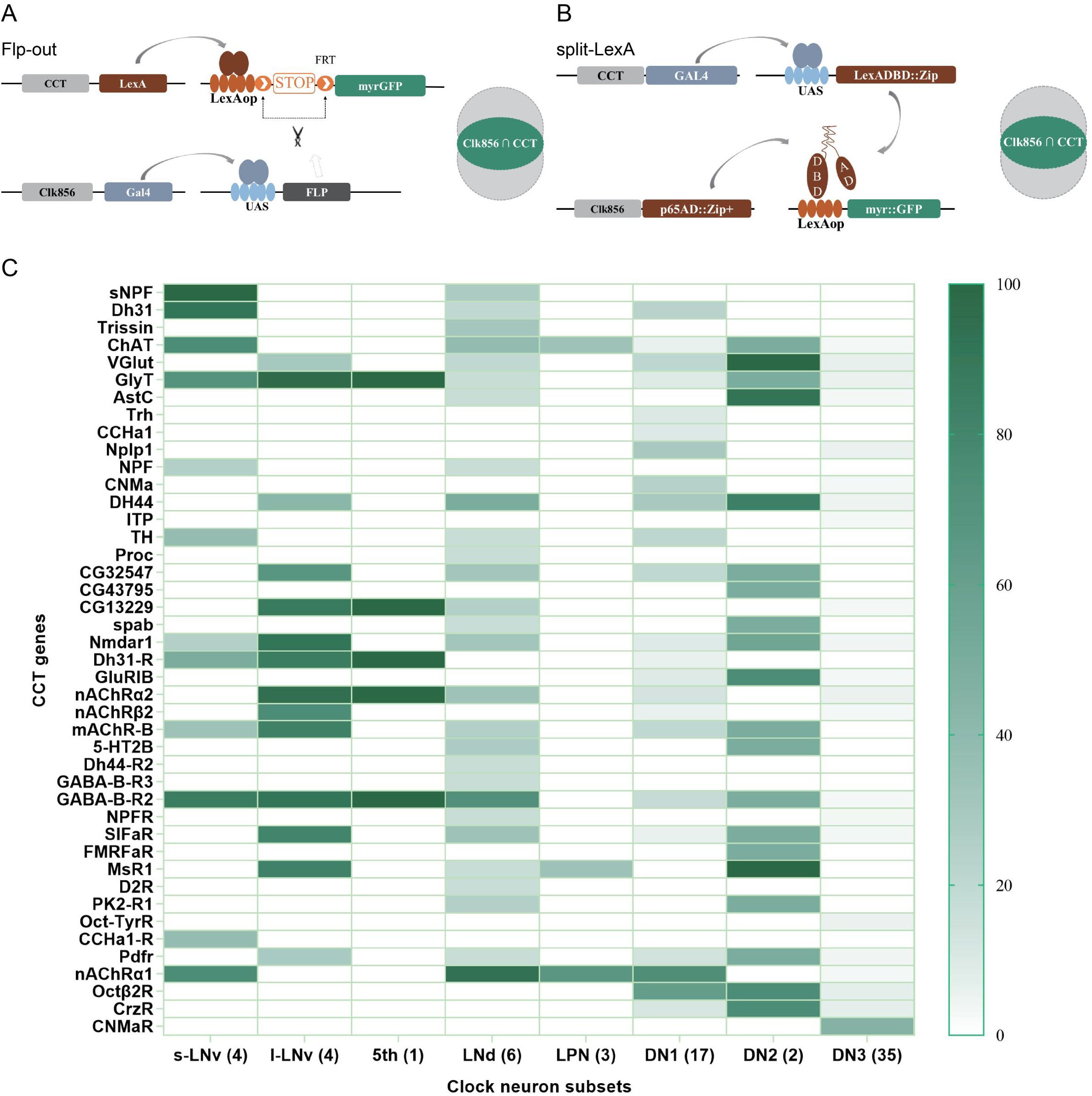
Genetic Dissection of Clk856 Labelled Clock Neurons. (A-B) Schematic of intersection strategies used in Clk856 labelled clock neurons dissection, Flp-out strategy (A) and split-LexA strategy (B). The exact strategy used for each gene is annotated in Table S5. (C) Expression profiles of CCT genes in clock neurons. Gradient color denotes proportion of neurons that were positive for the CCT gene within each subset. The exact cell number for each subset is annotated in Table S4.

To assess the accuracy of expression profiles using CCT drivers, we compared our dissection results with previous reports. Initially, we confirmed the expression of CCHa1 in two DN1s (Fujiwara et al., 2018), sNFP in four s-LNvs and two LNds (Johard et al., 2009), and Trissin in two LNds (Ma et al., 2021), aligning with previous findings. Additionally, we identified the expression of nAChRα1, nAChRα2, nAChRβ2, GABA-B-R2, CCHa1-R, and Dh31-R in all or subsets of LNvs, consistent with suggestions from studies using ligands or agonists in LNvs (Duhart et al., 2020; Fujiwara et al., 2018; Lelito and Shafer, 2012; Shafer et al., 2008) (Table S4).

Regarding previously reported Nplp1 in two DN1as (Shafer et al., 2006), we found approximately five DN1s positive for Nplp-KI-LexA, indicating a broader expression than previously reported. A similar pattern emerged in our analysis of Dh31-KI-LexA, where four DN1s, four s-LNvs, and two LNds were identified, contrasting with the two DN1s found in immunocytochemical analysis (Goda et al., 2016). Colocalization analysis of Dh31-KI-LexA and anti-PDF revealed labeling of all PDF-positive s-LNvs but not l-LNvs (Fig. S9A), suggesting that the differences may arise from the broader labeling of 3’ end knock-in LexA drivers or the amplitude effect of the binary expression system. The low protein levels might go undetected in immunocytochemical analysis. This aligns with transcriptome analysis findings showing Nplp1 positive in DN1as, a cluster of CNMa-positive DN1ps, and a cluster of DN3s(Ma et al., 2021), which is more consistent with our dissection.

Despite the well-known expression of PDF in LNvs and PDFR in s-LNvs (Shafer et al., 2008), we did not observe stable positive signals for both in Flp-out intersection experiments, although both Pdf-KI-LexA and Pdfr-KI-LexA label LNvs as expected (Fig. S9B-S9C). We also noted fewer positive neurons in certain clock neuron subsets compared to previous reports, such as NPF in three LNds and some LNvs (Erion et al., 2016; He et al., 2013; Hermann et al., 2012; Johard et al., 2009; Lee et al., 2006) and ChAT in four LNds and the 5th s-LNv (Duhart et al., 2020; Johard et al., 2009) (Table S4). We attribute this limitation to the inefficiency of LexAop-FRT-myr::GFP driven by LexA, acknowledging that our intersection results may miss some positive signals.

### Conditional Knockout of CCT Genes in Clock Neurons

To investigate the function of CCT genes in circadian neurons with our conditional knockout system, we knocked out all 67 (genes identified above and reported previously) CCT genes in Clk856-labeled clock neurons by C-cCCTomics.

In the pilot screen, we monitored fly activity by video recording (Dai et al., 2019) and analyzed rhythmic behavior under LD and DD conditions. We analyzed morning anticipation index (MAI) and evening anticipation index (EAI) under the LD condition (Harrisingh et al., 2007; Im and Taghert, 2010; Seluzicki et al., 2014) (Fig. 5A), power, period and arrhythmic rate (AR) under the DD condition. Fly activities tended to rise rapidly after ZT22.5 at dawn and ZT10 at dusk. Thus, we added two more parameters to describe the anticipatory activity patterns of LD condition. Morning anticipation pattern index (MAPI) was defined as the difference between *Pi*_a_[arctan(ZT20.5∼ZT22.5 activity increasing slope)] and *Pi*_p_[arctan(ZT22.5 ∼ZT24 activity increasing slope)], M(*Pi*_a_-*Pi*_p_). Evening anticipation pattern index (EAPI) was defined similar to MAPI (Fig. 5A, see Methods). *Pi*_a_ and *Pi*_p_ were positive, while MAPI and EAPI were negative, for wild type (wt) flies as their activities gradually increases at dawn or dusk at increasing rates.

**Fig. 5.**
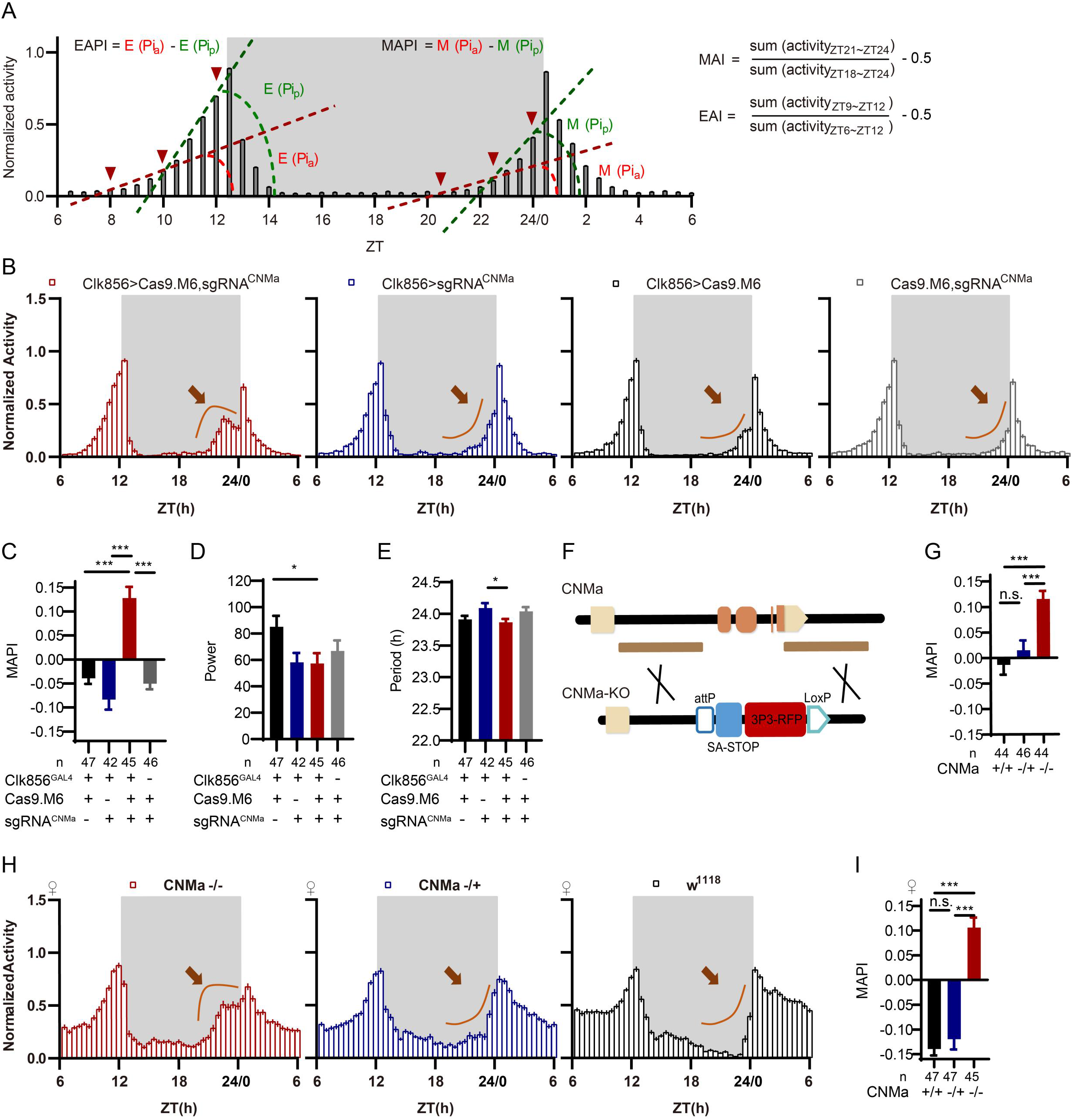
CNMa Regulation of Morning Anticipation in Clock Neuron. (A)Schematic of MAI, EAI, MAPI and EAPI definition. (B) Activity plots of male flies with CNMa knockout in clock neurons (red) and controls (blue, black and grey), plotted in 30 min bins. An advancement of morning activity peak was presented in CNMa clock-neuron-specific mutants (brown arrowhead). (C-E) Statistical analyses of MAPI, power, and period of flies in (B). MAPI was significantly increased in clock neurons-specific CNMa deficient flies (C) while power (D) and period (E) were not changed. (F) Schematic of CNMa^KO^ generation. The entire encoding region of CNMa was replaced by an attP-SAstop-3P3-RFP-loxP cassette using CRISPR-Cas9 strategy. (G) Statistical analysis of MAPI of male CNMa^KO^ flies (red) and controls (blue and black). MAPI significantly increased in male CNMa^KO^ flies. (H) Activity plots of female CNMa^KO^ flies (red) and controls (blue and black). (I) Statistical analysis of MAPI of female CNMa^KO^ flies (red) and controls (blue and black). MAPI was significantly increased in female CNMa^KO^ flies.

Knocking out Pdf or Pdfr in clock neurons phenocopied their mutants with lower MAI, advanced evening activity, low power, high arrhythmic rate and shorter period (Hyun et al., 2005; Lear et al., 2005; Renn et al., 1999) (Table S6, Fig S10A-S10D). The MAI-decreasing phenotype of Dh31 knockout was also reproduced in this pilot screen (Goda et al., 2019) (Table S6). All the above results verified the effectiveness of C-cCCTomics. Unexpectedly, additional experimental replications with full controls using Cas9.M9 revealed that leaky expression of Cas9.M9 and sgRNA might have caused disruption of Dh31, Dh44, Pdf and Pdfr (Fig. S10A-S10D) (see DISCUSSION), which was not suitable for neuronal specific mutagenesis of some genes. Therefore, in the following work we primarily focused on Cas9.M6 instead.

Analysis of the newly defined parameters MAPI and EAPI showed that control flies (Clk856-GLA4>UAS-Cas9.M9) had negative EAPI but slightly positive MAPI. The positive MAPI of control flies in this screen might be caused by Cas9.M9 toxicity. Only the Pdf and Pdfr clock-neuron knockout flies showed positive EAPIs, indicating an advanced evening activity (Table S6, Fig S10A, S10C). nAChRα1, MsR1, mAChR-B, and CNMa cKO flies had highest MAPI values (Table S6). We further confirmed their phenotypes using Cas9.M6 which revealed that CNMa plays a role in regulating morning anticipatory activity (Fig. S11A).

### Regulation of Morning Anticipation by CNMa-positive DN1p Neurons

Conditionally knocking out CNMa in clock neurons advanced morning activity (Fig. 5B, Fig. S11B) and increased MAPI (Fig. 5C, Fig S11A, S11C), leaving the power and period intact in male flies (Fig. 5D, 5E). The same advanced morning activity phenotype were also observed in female flies (Fig. S11D-S11G). To further validate this phenotype, we generated a CNMa knockout (CNMa^KO^) line by replacing its whole coding region with an attP-splicing adaptor element (Deng et al., 2019) (Fig. 5F). Both male and female CNMa^KO^ flies exhibited the same phenotypes as seen in the CNMa cKO (Fig. 5G-5I).

Previous studies have found CNMa expression in DN1 neurons (Abruzzi et al., 2017; Jin et al., 2021; Ma et al., 2021). Our intersection showed four DN1p and one DN3 CNMa positive neurons in Clk856 labelled neurons (Fig. S8 - 16, Fig. 4C, Table S4). Analysis with an endogenous CNMa-KI-GAL4 knockin driver showed that six pairs of CNMa neurons located in the DN1p region and three pairs located in the subesophageal ganglion (SOG) had the brightest GFP signals (Fig. 6A). The anatomical features of CNMa neurons were further confirmed using stingerRed and more neurons were found in regions, the anterior ventrolateral protocerebrum (AVLP), and the antennal mechanosensory and motor center (AMMC) (Fig. S12A). Dendrites of CNMa neurons were concentrated in DN1p and SOG, with their axons distributed around DN1p region, lateral horn (LH), and prow region (PRW) (Fig. 6B, 6C). Using the Trans-tango strategy (Talay et al., 2017), we also found that downstream of CNMa neurons were about fifteen pairs of neurons in the SOG, five pairs of LNd neurons, one pair of DN3 neurons and six intercerebralis (PI) neurons (Fig. 6D, arrowhead).

**Fig. 6.**
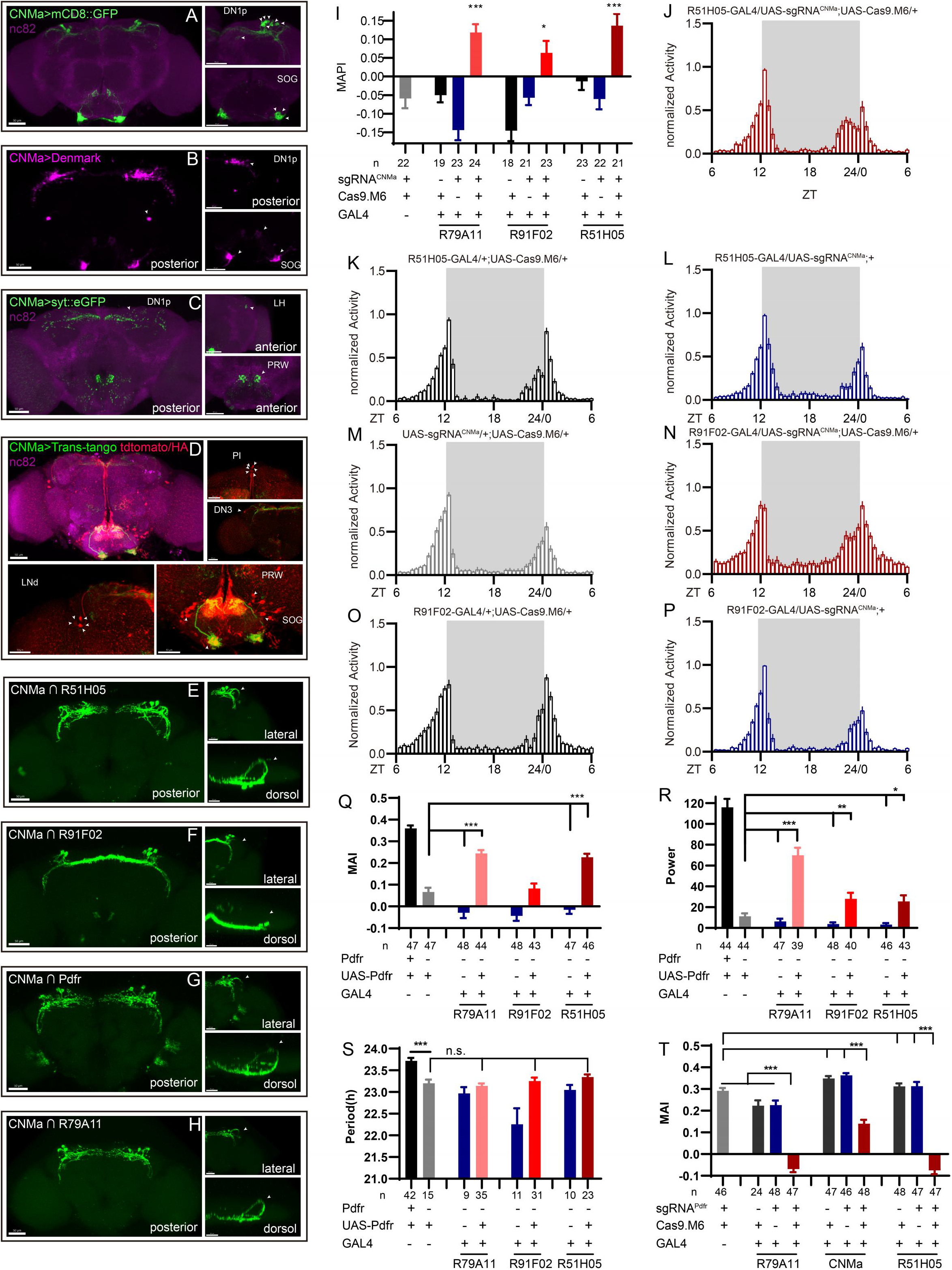
Expression, Projection and Trans-projection Feature of CNMa Neurons and Its Functional Subset. (A-C) Expression and projection patterns of CNMa-KI-Gal4 in the brain. Membrane, dendrites, and axon projections are labelled by mCD8::GFP (A), Denmark (B), and syt::eGFP (C) respectively. (D) Downstream neurons labelled through trans-tango driven by CNMa-KI-GAL4. Arrowheads indicate candidate downstream neurons: six neurons in PI, one pair in DN3, five pairs in LNd and about 15 pairs in SOG. (E-H) Intersection of DN1p CNMa neurons with DN1p labelled drivers. GMR51H05-GAL4 (E), GMR91F02-GAL4 (F), Pdfr-KI-GAL4 (G) and GMR79A11-GAL4 (H) were intersected with CNMa-p65AD, UAS-LexADBD, LexAop-myr::GFP. Two type I (E, G, H) neurons projected to anterior region and four type II (F) neurons had fewer projections to anterior region. Scale bar, 50μm. (I) MAPIs were significantly increased in all three DN1p drivers mediated CNMa knockout. (J-K) Activity plots of CNMa knockout in R51H05-GAL4 (J) and R91F02-GAL4 (N) neurons. R51H05-GAL4 mediated CNMa knockout flies showed an advanced morning activity peak (J), while R91F02-Gal4 mediated CNMa knockout flies did not (N). (Q-S) Statistical analyses of MAI, power and Period. Pdfr reintroduction in R79A11 and R51H05 neurons could partially rescue the MAI-decreased phenotype of Pdfr knockout flies. (T) Statistical analyses of MAI of Pdfr knocking out in R79A11-GAL4, CNMa-GAL4 and R51H05-GAL4 labelled neurons.

Because we had found that knocking out CNMa in Clk856-GAL4 labeled neurons produced advanced morning activity, and that CNMa intersected with Clk856-Gal4 labeled neurons in 4 pairs of DN1ps and one pair of DN3 neurons (Fig. S8-16), we focused on these neurons and performed more intersections. Taking advantage of a series of clock neuron subset-labeled drivers (Sekiguchi et al., 2020), we intersected CNMa-p65AD with 4 DN1 labelling drivers: GMR51H05-GAL4, GMR91F02-GAL4, Pdfr-KI-GAL4 and GMR79A11-GAL4 (Fig. 6E-6H). We found two arborization patterns: Type I with two neurons whose branches projecting to the anterior region, as observed in CNMa∩GMR51H05, CNMa∩Pdfr, and CNMa∩GMR79A11 (Fig 6E, 6G, 6H), and type II with four neurons branching on the posterior side with few projections to the anterior region, as observed in CNMa∩GMR91F02 (Fig. 6F). These two types of DN1ps’ subsets have been previously reported and profound discussed (Lamaze et al., 2018; Reinhard et al., 2022).

CNMa knockout in Type I or Type II neurons (GMR51H05-GAL4, GMR91F02-GAL4, and GMR79A11-GAL4) all reproduced the MAPI-increased phenotype of clk856 specific CNMa knockout (Fig. 6I). However, Type II neurons-specific CNMa knockout (CNMa ∩ GMR91F02) showed weaker advanced morning activity without advanced morning peak (Fig. 6N), while Type I neurons-specific CNMa knockout did (Fig. 6J), indicating a possibility that these two type I CNMa neurons are the main functional subset regulating the morning anticipation activity of fruit fly.

Pdf or Pdfr mutants exhibit weak or no morning anticipation, which is related to the phenotype of CNMa knockout flies. We also identified two Pdfr and CNMa double-positive DN1ps, which have a type I projection pattern (Fig. 6G). Reintroduction of Pdfr in Pdfr knockout background revealed that GMR51H05 and GMR79A11 Gal4 drivers, which covered the main functional CNMa-positive subset, could partially rescue the morning anticipation and power phenotype of Pdfr knockout flies to a considerably larger extent than the GMR91F02 driver (Fig. 6Q-6S, Fig. S13A, Table S7). Moreover, knocking out Pdfr by GMR51H05, GMR79A11 and CNMa GAL4, which cover type I CNMa neurons, decreased morning anticipation of flies (Fig. 6T, Fig. S13B). However, the decrease in morning anticipation observed in the Pdfr knockout by CNMa-GAL4 was not as pronounced as with the other two drivers. Because the presumptive main subset of functional CNMa is also PDFR-positive, there is a possibility that CNMa secretion is regulated by PDF/PDFR signal.

### Role of Neuronal CNMaR in Morning Anticipation

There is only one CNMa receptor reported in the fly genome (Jung et al., 2014). We generated a CNMaR^KO-p65AD^ line by CRISPR/Cas9 (Fig. 7A) and this knockout showed advanced morning activity (Fig. 7B, 7D) and increased MAPI (Fig. 7C, 7E) in both sexes. CNMaR^KI-Gal4^/UAS-mCD8::GFP and CNMaR^KI-Gal4^/UAS-stinger::Red showed expression of CNMaR across the whole brain (Fig. 7F, Fig S12B), especially in DN1p, DN3, the PI and the SOG. The dendrite arborization and synaptic projections of CNMaR neurons also covered broad regions (Fig. 7G, 7H), at the PI, the SOG, the posterior ventrolateral protocerebrum (PVLP) and the central complex (CC). Further conditional knockout of CNMaR in neurons by C-cCCTomics phenocopied CNMaR^KO-p65AD^ phenotype (Fig. 7I-7L). These results indicate that CNMaR is similar to CNMa in regulating morning anticipation.

**Fig. 7.**
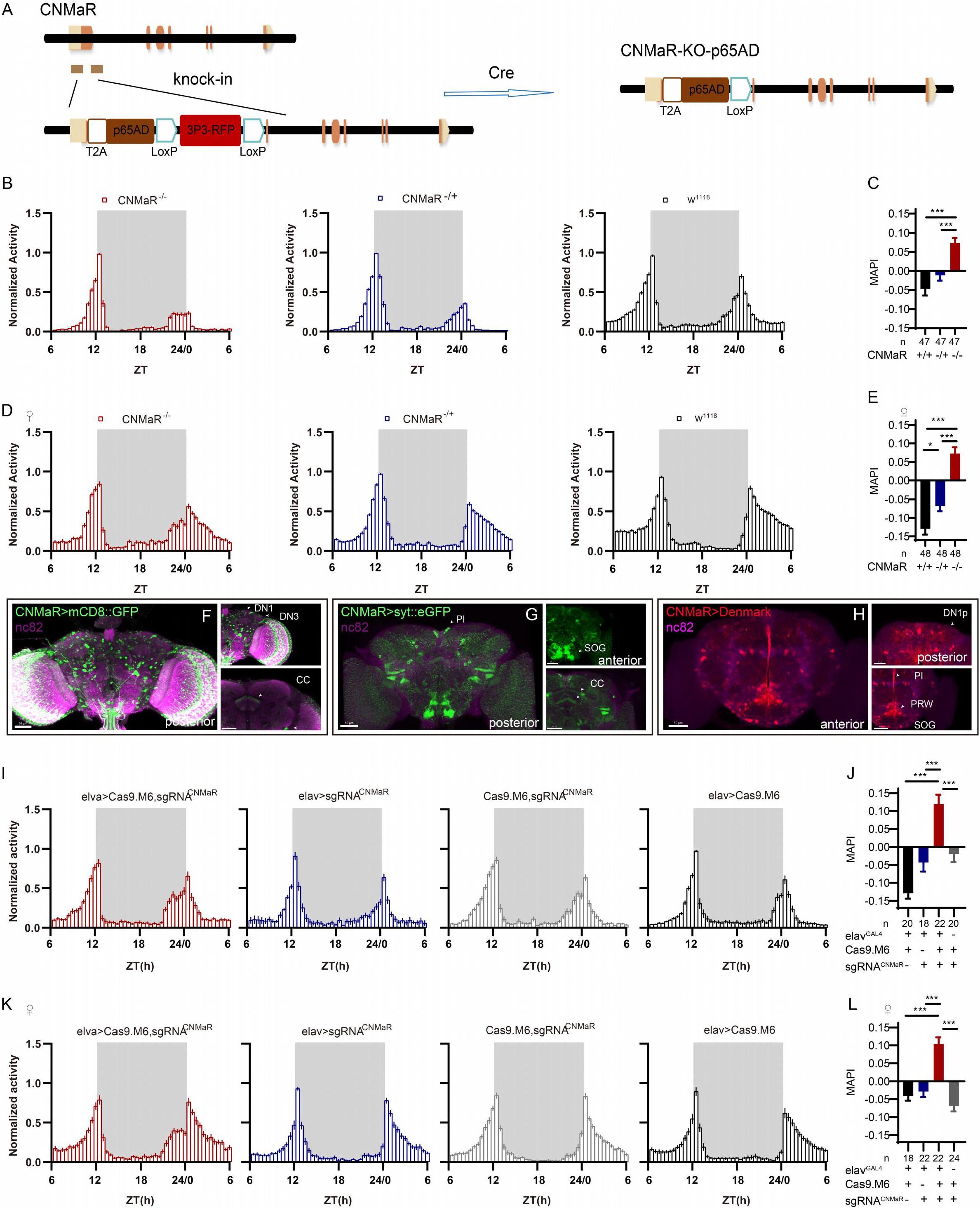
CNMaR Regulation of Morning Anticipation. (A) Schematic of CNMaR^KO-p65AD^ generation. Most of the first exon in CNMaR was replaced by a T2A-p65AD-loxP-3P3-RFP-loxP cassette using CRISPR-Cas9 strategy and the T2A-p65AD was inserted in the reading frame of the remaining CNMaR codon. 3P3-RFP was removed latterly by Cre mediated recombination. (B-E) Activity plot (B, D) and statistical analysis (C, E) of male (B-C) or female (D-E) CNMaR^KO-p65AD^ flies (red) and genotypical controls (blue and black). MAPI was significantly increased in both male and female CNMaR^KO-p65AD^ flies. In this and other figures, “♀” denotes female flies. (F-H) Expression and projection patterns of CNMaR-KI-Gal4 in the brain. Scale bars, 50μm. (I-L) Activity plots (I, K) and statistical analyses (J, L) of CNMaR pan-neuronal knockout flies. Neuronal knockout of CNMaR increased MAPI (J, L).

## DISCUSSION

### Conditional CCTomics Strategies and Toolkit

We have generated conditional gene manipulation systems based on Flp-out/GFPi or CRISPR/Cas9. cCCT based gene deletion after heat-shock or mifepristone (RU486) eliminated most GFP signals, and pan-neuronal constitutive expression of shRNA^GFP^ or flippase disrupted seven out of eight tested genes completely while targeting of SIFa^EGFP.FRT^ achieved 96±3% efficiency. Although the recombination of genetic elements is relatively cumbersome when using cCCTomics, it is worthwhile to apply this method to specific genes given its high level of efficiency. While two UAS-sgRNA libraries have been established, one primarily targeting kinases (Port et al., 2020) and the other targeting GPCRs (Schlichting et al., 2022), both libraries only cover a portion of CCT genes, and are thus insufficient for manipulating all CCT genes. The development of C-cCCTomics, however, makes CCT gene manipulation as simple as RNA interference. Furthermore, the use of modified Cas9.M6 or Cas9.M9 highly enhances the efficiency of gene disruption in the nervous system, allowing for efficient manipulation of all CCT genes in a cell-specific manner.

The toxicity of CRISPR/Cas9 depends on the Cas9 protein (Port et al., 2014). When expressed pan-neuronally in nSyb-GAL4 (R57C10-GAL4, attP40), Cas9.M9 slightly reduced viability while the expression of other Cas9 variants had no significant effect on viability (Fig. S14). Although Cas9.M9 showed leaky-expression-efficiency, this was not a problem with Cas9.M6, which successfully disrupted Dh31, Dh44, Pdf, and Pdfr (Fig. S9). A more restricted expression of Cas9.M9 with lower toxicity is necessary for better somatic gene manipulation in the future.

### CCT of Clock Neurons

Intersecting Clk856-Gal4 or Clk856-p6AD with CCTomics, we identified 45 CCT genes in Clk856 labelled clock neurons. Clock neurons appear highly heterogeneous both in our intersection dissection and in a previous transcriptomic analysis (Abruzzi et al., 2017; Ma et al., 2021). Comparing these two CCT gene expression profiles in clock neurons, 41 out of 127 gene-subsets are identical. The accuracy of our genetic intersection is limited by two possibilities: 1) KI-LexA may not fully represent the expression pattern of the corresponding gene, and 2) the efficiency of STOP cassette removal in the Flp-out strategy is limited or the efficiency of LexA>LexAop::myrGFP. Moreover, the leakage of LexAop-GFP may result in unreliable labelling in split-lexA strategy. Both genetic drivers and transcriptomic analysis contribute to our knowledge of the expression profile of neurons. The physiological significance of each gene in particular neurons should be further investigated by genetic manipulation.

### Regulation of Rhythmic Behavior by CCT genes

Multiple attractive genes have been identified in our functional screen of CCT genes in clock neurons: for example, knocking out of VGlut weakens morning anticipation (Table S8). In further screening of brain regions, we have narrowed down the morning anticipation regulation role of VGlut in R18H11-GAL4 labeled neurons (Table S9). VGlut in these neurons has also been reported to regulate sleep in Drosophila (Guo et al., 2016). Its downstream neurons may be the PI neurons or LNvs (Barber et al., 2021; Guo et al., 2016).

Moreover, the deficiency of the neuropeptide CNMa results in advanced morning activity. We have validated that two Pdfr and CNMa double-positive DN1p neurons may play a major role in regulating this process through intersectional manipulation of CNMa. Knockout and reintroduction of Pdfr in these neurons have verified that Pdfr partially functions in DN1p CNMa neurons, and PDF increases cAMP level in Pdfr positive neurons (Shafer et al., 2008), suggesting a possibility of the regulation of CNMa signaling by PDF signaling. Furthermore, given that the morning anticipation vanishing phenotype of Pdf or Pdfr mutant indicates a promoting role of PDF-PDFR signal, while the enhanced morning anticipation phenotype of CNMa mutant suggests an inhibiting role of CNMa signal, we consider the two signals to be antagonistic. However, knocking out CNMaR in Clk856 labelled clock neurons showed no significant phenotype (Table S6), whereas the mutant and pan-neuronal knockout flies had similar phenotypes to CNMa knockout flies, suggesting its role in the circadian output neurons. Previous studies have indicated that CNMa integrate thermosensory inputs to promote wakefulness, and CNMaR is thought to function in Dh44 positive PI neurons (Jin et al., 2021), a subset of circadian output neurons. To gain a deeper understanding of the downstream effects of DN1p CNMa positive neurons, further analysis focusing on specific brain regions is necessary.

We have also reproduced phenotypes of Pdf, Pdfr, and Dh31 mutant flies with C-cCCTomics as previous studies. Surprisingly, only five genes are functional among all sixty-seven CCT genes in this prior screen. This may be caused by limitations of the simple behavioral paradigm, single gene manipulation, and single GAL4 driver. For example, switching of light condition from L: D = 12h: 12h to L: D = 6h:18h, AstC/AstC-R2 would suppress flies’ evening activity intensity to adapt to the environmental change (Díaz et al., 2019), and only double knockout of AChRs and mGluRs in PI neurons can possibly result in alteration in behavioral rhythms (Barber et al., 2021). Further diversified functional analysis of CCT genes in clock neurons is required for clock circuit dissection.

## Methods

### Fly Lines and Rearing Conditions

Flies were reared on standard corn meal at 25 °C, 60% humidity, 12 h light: 12 h dark (LD) cycle. For flies used in behavior assays, they were backcrossed into our isogenized Canton S background for 5 to 7 generations. For heat induced assays, flies were reared at 20°C. All CCT attP KO lines and CCT KI driver lines were previous generated at our lab (Deng et al., 2019). Clk856-GAL4 and GMR57C10-GAL4 driver lines were gifts from Donggen Luo Lab (Peking University). 13XLexAop2 (FRT.stop) myr::GFP was gift from Rubin Lab.

### C-cCCTomics sgRNAs design

All sgRNAs target at or before functional coding regions (e.g. GPCR transmembrane domain, synthetase substrate binding domain) of each CCT genes. For each gene, about 20 sgRNAs with specific score ≥ 12 were firstly designed at CRISPRgold website (Chu et al., 2016; Graf et al., 2019), then their specificity and efficacy were further valued in Optimal CRISPR target finder(Gratz et al., 2014), E-CRISPR(Heigwer et al., 2014) and CCTop (Stemmer et al., 2015) system. The first three highest efficacy sgRNAs with no predicted off-target effect were selected. All selected sgRNAs are listed in Table S3.

### Molecular Biology

All cCCTomics knockin (KI) lines and C-cCCTomics transgenic flies were generated through phiC31 mediated attB/attP recombination, and the miniwhite gene was used as selection marker.

For cCCTomics KI lines, backbone pBSK-attB-FRT-*Hpa*I-T2A-EGFP-FRT was modified from pBSK-attB-loxP-myc-T2A-Gal4-GMR-miniwhite (Deng et al., 2019). Myc-T2A-GAL4 cassette was removed by PCR amplification while first FRT cassette was introduced. Second FRT cassette was inserted by T4 ligation between *SpeI* and *BamHI*. T2A-EGFP was cloned from pEC14 and was inserted into the backbone between two FRT cassettes. All gene spans, except for stop codon, deleted in CCT attP KO lines were cloned into pBSK-attB-FRT-*Hpa*I-T2A-EGFP-FRT at *Hpa*I site.

For C-cCCTomics UAS-sgRNA lines, backbone pMsgNull was modified from pACU2 (Han et al., 2011). Synthetic partial fly tRNA^Gly^ sequence was inserted between *EcoR*I and *Kpn*I. An irrelevant 1749 bp cassette amplified from pAAV-Efla-DIO-mScarlet (addgeen#130999) was inserted between *Eag*I and *Kpn*I. All sgRNA spacers were synthesized at primers, and “E+F” sgRNA scaffold and rice tRNA^Gly^ was amplified from a synthetic backbone PM04. Finally, gRNA-tRNA^Gly^ cassettes were cloned into pMsgNull between *Eag*I and *Kpn*I by Gibson Assembly.

All UAS-Cas9 variants generated in this research were cloned into vector pACU2 and all Cas9 sequence were amplified from hCas9 (addgene#41815). Human codon optimized Cas9 was cloned into pACU2 to generate UAS-Cas9.HC. UAS-Cas9.M0 was modified from UAS-Cas9.HC by introduce a 3x HA tag after NSL and replace the SARD linker with the 3×GGSGP linker (Zhao et al., 2016). UAS-Cas9.M6 and UAS-Cas9.M9 were designed as HMGN1-Cas9-UPD and HMGN1-Cas9-HB1 respectively. All these chromatin-modulating peptides were linked with Cas9 by 3x GGSGP linker.

CNMaR^KO-p65AD^ is generated by replace the coding region of the first exon with T2A-p65AD by CRISPR/Cas9, the T2A-p65AD was linked in frame after first ten amino acids. Spacers of gRNAs used to break the targeted CNMaR region were 5’-GCAGATTTCAGTTCATCTTT-3’, 5’-GGCTTGGCAATGAC TATATA-3’.

### Gene expression quantitation and high-throughput sequencing

Female flies were gathered six to eight days post-eclosion for gene expression quantification and high-throughput sequencing. Fly heads were isolated by chilling them on liquid nitrogen and subsequent shaking. mRNA extraction was performed using Trizol according to a previously established protocol (Green and Sambrook, 2020). Genomic DNA was removed, and cDNA was synthesized using a commercial kit (TIANGEN#DP419). For real-time quantitative PCR, at least one PCR primer was designed to overlap with the sgRNA target site.

Genomic DNA from fly heads was extracted using a standard alkali lysis protocol(Huang et al., 2009). Genomic regions approximately 130 to 230 bp in length, centered around the sgRNA target site, were amplified by PCR employing Q5 polymerase (NEB#M0494). Subsequently, libraries were prepared using the BTseq kit (Beijing Tsingke Biotech Co., Ltd.). These libraries were pooled and subjected to sequencing on the MiSeq platform (Illumina). Analysis of the libraries was conducted using Crispresso2(Clement et al., 2019).

### Generation of KI and Transgenic lines

Generation of cCCTomics KI, CNMa^KI-p65AD^ and CNMaR^KO-p65AD^ lines are the same as generation of CCTomics KI driver lines as previously described (Deng et al., 2019). To generate C-cCCTomics UAS-sgRNA or UAS-Cas9 variants lines, attB vectors were injected and integrated into the attP40, attP2 or VK00005 through phiC31 mediated gene integration.

All flies generated in this research were selected by mini-white and confirmed by PCR.

### Behavioral assays

Unmated male or female flies of 4 to 5 days were used in circadian rhythm assays. Before measurement, flies were entrained under 12 h light: 12 h dark cycle at 25°C for at least 3 days and then transferred to dark-dark condition for 7 days.

Virgin flies of 4 to 5 days were used in sleep assays. Flies were entrained to a 12 h light: 12 h dark cycle at 25°C for 2 days to eliminate the effect of CO2 anesthesia before sleep record. Sleep was defined as 5 min or longer immobility (Hendricks et al., 2000; Shaw et al., 2000) and analyzed by in-house scripts as previously described ((Dai et al., 2021; Dai et al., 2019; Deng et al., 2019).

Locomotion was obtained as previously described (Dai et al., 2021). Locomotion activity was measured and analyzed by Actogram J plugin (Dai et al., 2019). Morning anticipation index and evening anticipation index were defined as the ratio of last 3 hours activity before light-on or light-off accounts to last 6 hours activity before light-on or light-off, further subtracted by 0.5(Index=sum(3hrs)/sum(6hrs)-0.5) (Harrisingh et al., 2007; Im and Taghert, 2010; Seluzicki et al., 2014) and analyzed by an in-house python script.

### Heat Shock and Drug Treatment

For hsFLP mediated conditional knockout, flies of 4 to 6 days were heat shock at 37°C during ZT10 to ZT12 for 4 days. And they were reared at 20°C for another 4-days and then dissected.

For mifepristone (RU486) induced conditional knockout, flies of 4 to 6 days were treated with 500 μM RU486 mixed in corn food and then dissected 4 days later.

### Immunohistochemistry and Confocal Imaging

For all imaging without staining, adult flies were anesthetized on ice and dissected in cold phosphate-buffer saline (PBS). Brains or ventral nerve codes (VNC) were fixed in 2% paraformaldehyde (weight/volume) for 30 min, washed with washing buffer (PBS with 1% Triton X-100, v/v, 3% NaCl, g/mL) for 7 min three times and mounted in Focusclear (Cell Explorer Labs, FC-101).

For imaging with staining, brains and VNCs were fixed for 30 min, washed for 15 min three times. Then they were blocked in PBSTS, incubated with primary antibodies, washed with washing buffer, incubated with second antibodies and mounted as described previously(Dai et al., 2021; Dai et al., 2019).

All brains or VNCs were imaged on Zeiss LSM710 or Zeiss LSM880 confocal microscope and processed by Imaris.

The following primary antibodies were used: mouse anti-PDF (1:200, DSHB), rabbit anti-TH (1:1000, Novus Biologicals), rabbit anti-LK (1:1000, RaoLab, this paper). Rabbit anti-DSK (1:1000) was a gift from Dr. C. Zhou Lab (Institute of Zoology, Chinese Academy of Science) (Wu et al., 2020). The following secondary antibodies were used: AlexaFluor goat anti-mouse 488 (1:1000, Invitrogen), AlexaFlour goat anti-rabbit 488/633 (1:1000, Invitrogen).

For Fig 2, number of TH positive neurons were counted with Imaris Spots plugin.

### Quantification and Statistics

MAI, MAPI, EAI, EAPI, power and period were calculated by python or R scripts. ZT0 was set as the time point when light was on and ZT12 was set as the time point for light-off. Activity bins started at ZT0 and each was calculated as a sum of the total activity within 30min. Flies were regarded as death and removed if their activity value within last 2 bins was 0. A representative 24hr activity pattern was the average between corresponding activity bins from 2 consecutive days. To minimize effects from singular values, each flies’ activity was normalized using the following formula:

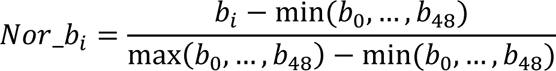

bi means the activity value for a certain bin. min(b0,…,b48) means the minimal bin value within 24hr and max(b0,…,b48) means the maximal bin value within 24hr. Nor_bi means the final normalized bin value for a certain bin from a given fly. Normalized activity was used for following analysis.

Morning activity arise (M_arise) was defined as the radian between the activity curve (ZT21-ZT22) and the time coordinate. Morning activity plateau (M_plateau) was defined as the radian between the activity curve (ZT22.5-ZT24) and the time coordinate. Evening activity arise (E_arise) was defined as the radian between the activty curve (ZT8-ZT11.5) and the time coordinate. Evening activity plateau (E_plateau) was defined as the radian between the activity curve (ZT10.5-ZT12) and the time coordinate. Morning anticipation pattern index (MAPI) was calculated by subtracting M_arise from M_plateau and Evening anticipation pattern index (EAPI) was calculated by subtracting E_arise from E_plateau.

The original activity data from 7 consecutive days in dark-dark condition was used for power and period calculation as described (Geissmann et al., 2019). Each flies’ periodogram was calculated based on Chi-Square algorithm (Sokolove and Bushell, 1978) and flies with a null power value were regarded as arrhythmic.

All statistical analyses were carried out with Prism 8 (Graphpad software). The Kruskal-Wallis ANOVA test followed by Dunn’s posttest was used to compare multiple columns.

## ACKNOWLEDGEMENTS.

We are grateful to Xiaofei Liu, Xueying Wang and Yile Jiao for sgRNA design and cloning, to Linyi Zhang, Wenli Xu, Yuqing Gong and Quiquan Chen for assistance with behavior assays, to Xiao Dong for cloning, to Xinwei Gao for assistance with imaging, to Pingping Yan, Lan Wang, Yonghui Zhang and Haixia Zeng for fly rearing, to Enxing Zhou and Wei Yang for Drosophila activity tracing scripts, to Yujie Li for cartoon illustration, to Donggen Luo, Chuan Zhou, Gerald M. Rubin, Vienna Drosophila RNAi Center and Bloomington Drosophila Stock Center for flies, to Yuh-Nung Jan for pACU2 construct.

## Author Contributions

Y.R. supervised the project. R.M. and J.Y. performed majority of experiments and data analysis. B.D., J.Y., and R.M. found advanced morning activity phenotype and carried out experiments of CNMa/CNMaR. R.M., X.D., Y.D., and S.D. carried out experiments of cCCTomics fly generation. R.M. and Y.R. wrote the manuscript.

## Supplementary Information

**Fig. S1.**
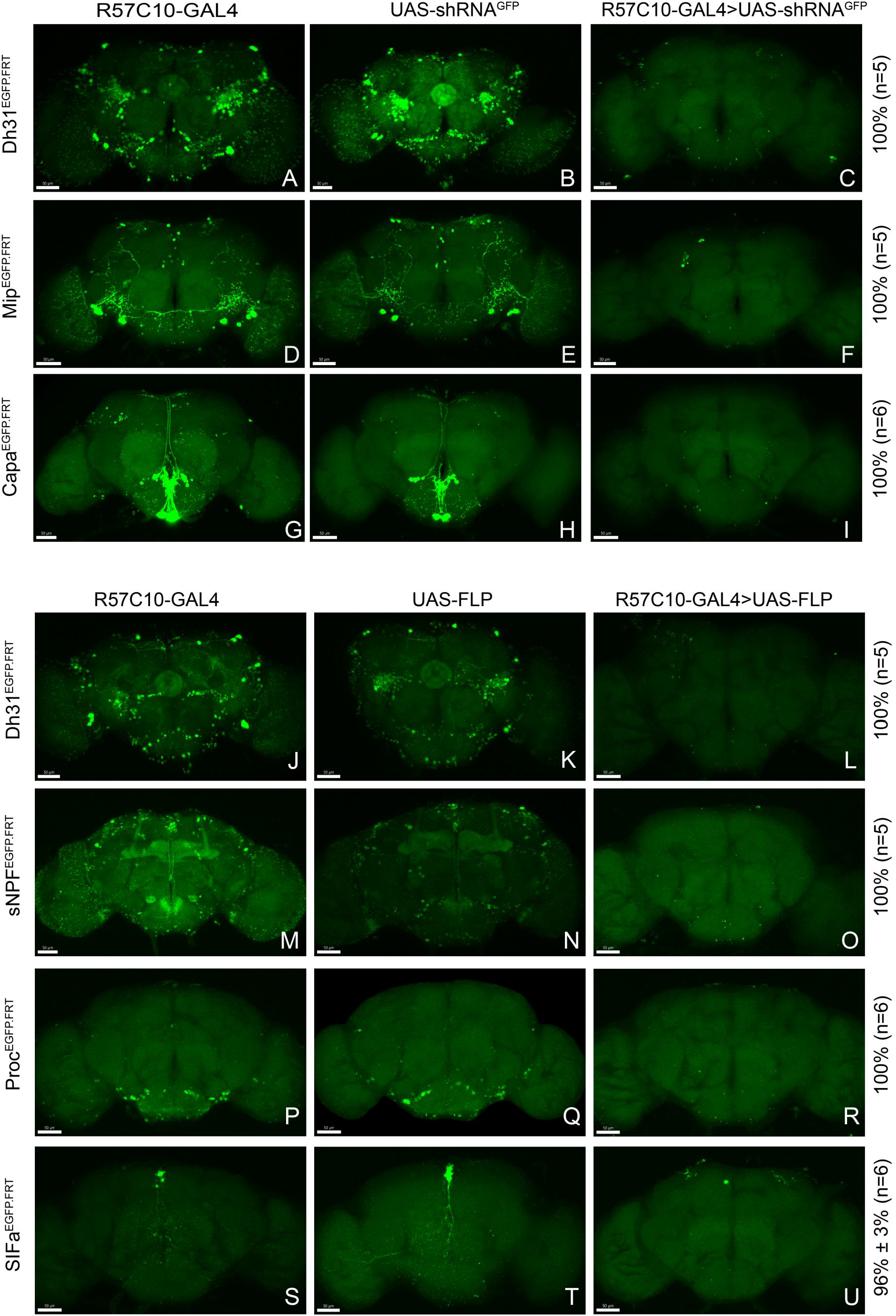
Efficient Conditional Disruption of CCT Genes by cCCTomics. (A-I) Pan-neuronal knockdown of Dh31 (A-C), Mip (D-F) and Capa (G-I) by GFPi. All experimental fly brains (C, F, I) showed no GFP signal after knockdown by GFPi. As described in Fig 1, all percentages in the right represent gene disruption efficiency and n represent number of experimental flies. (J-U) Pan-neuronal knock out of Dh31, sNPF, Proc and SIFa by Flp-out strategy. No obvious GFP signal was found in the experimental fly brains (L, O, R), excepting one GFP positive neurons in SIFa flp-out group (U).

**Fig. S2.**
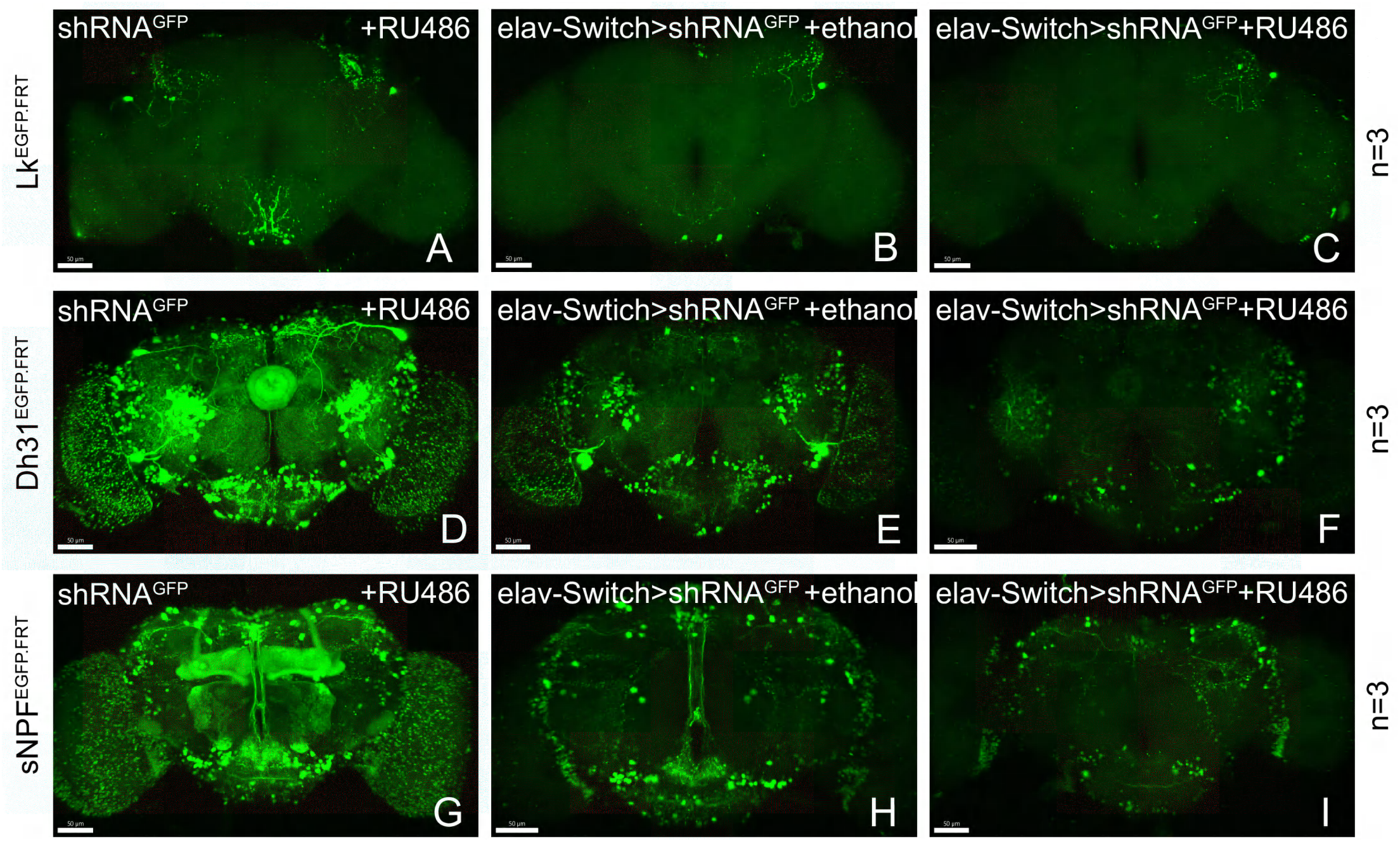
Gene Disruption of Target Genes by Induced shRNA. (A-J) Knockdown of Lk, Dh31 and sNPF by elav-Switch. Most GFP signal is lost in the experimental group (C, F, I).

**Fig. S3.**
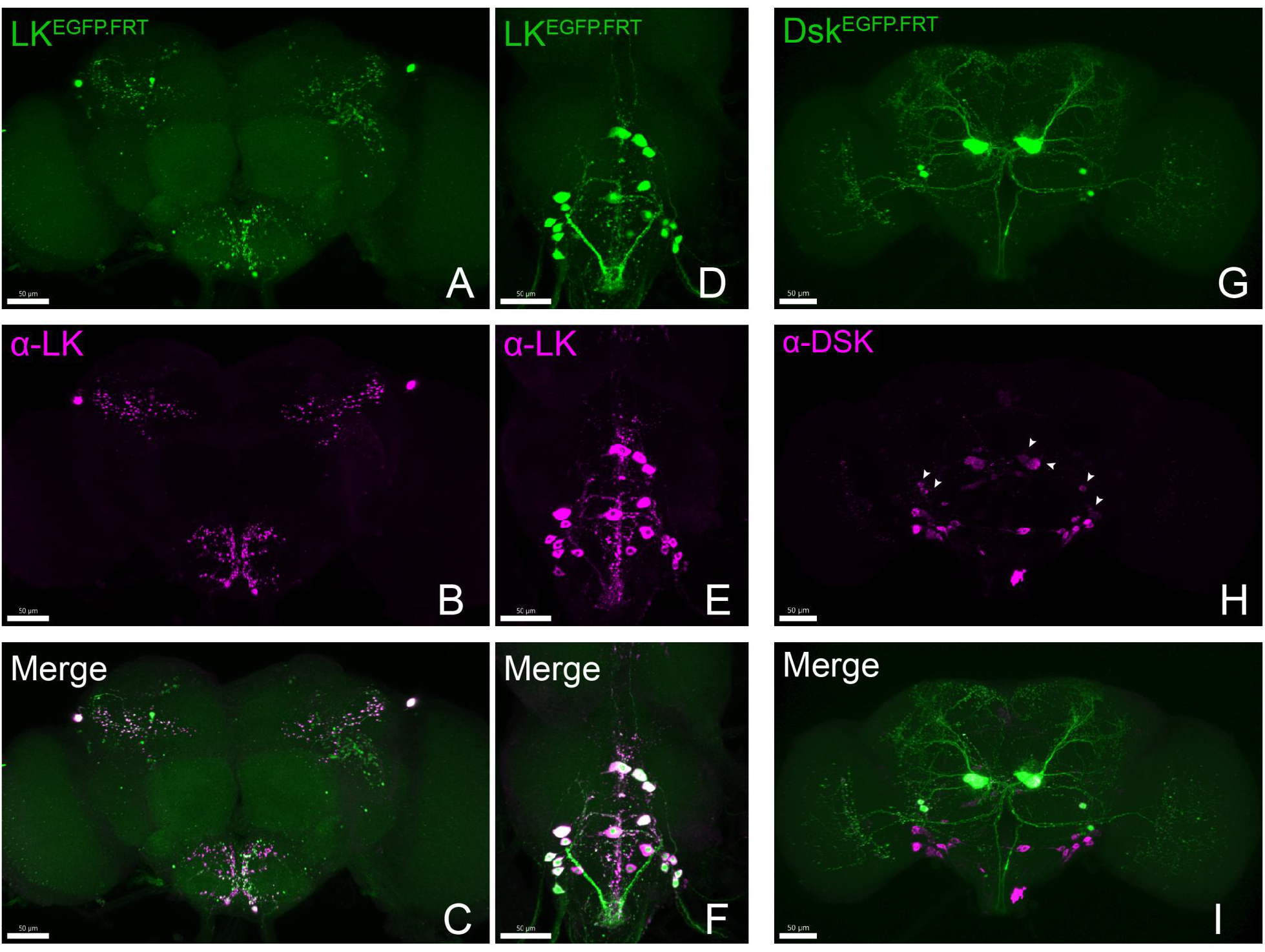
Accurate Labeling of Target Genes by cCCT Lines. (A-J) Co-localization of fused EGFP labelled CCT genes and corresponding antibodies. All the EGFP labelled Lk (A, D) and Dsk (G) neurons (Green) were co-localized with anti-LK (B, E; merge C, F) and anti-DSK (H, merge I) signal (purple). Arrowheads represent specific anti-DSK signal (Wu et al., 2020).

**Fig. S4.**
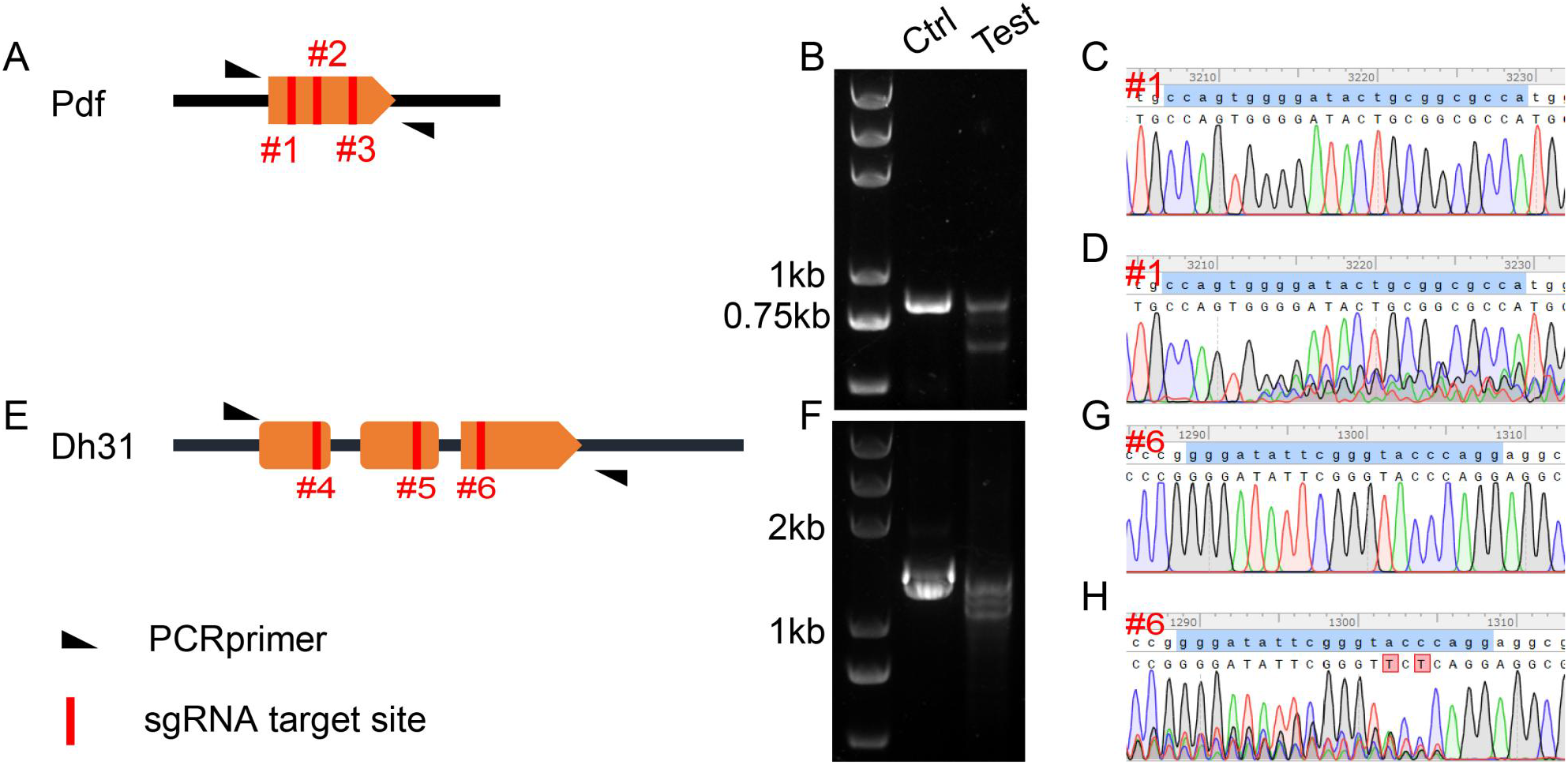
Validation of Primary C-cCCTomics. (A) Gene span of Pdf. #1, #2 and #3 with red lines denote sgRNA targets regions in Pdf coding sequence. (B) DNA gel electrophoresis of PCR products after *pdf* KO by C-cCCTomics. “Ctrl” denotes PCR products from genomic DNA mixture of Act5C-GAL4/+; UAS-Cas9.P2/+ flies and UAS-sRNA^Pdf^ flies. “Test” denotes PCR products from Act5C-GAL4/ UAS-sRNA^Pdf^; UAS-Cas9.P2 flies. (C-D) Sanger sequencing results of Ctrl (C) and Test (D) PCR products at #1 sgRNA target site. (E-H) Similar to (A-D) with Dh31 as target.

**Fig. S5.**
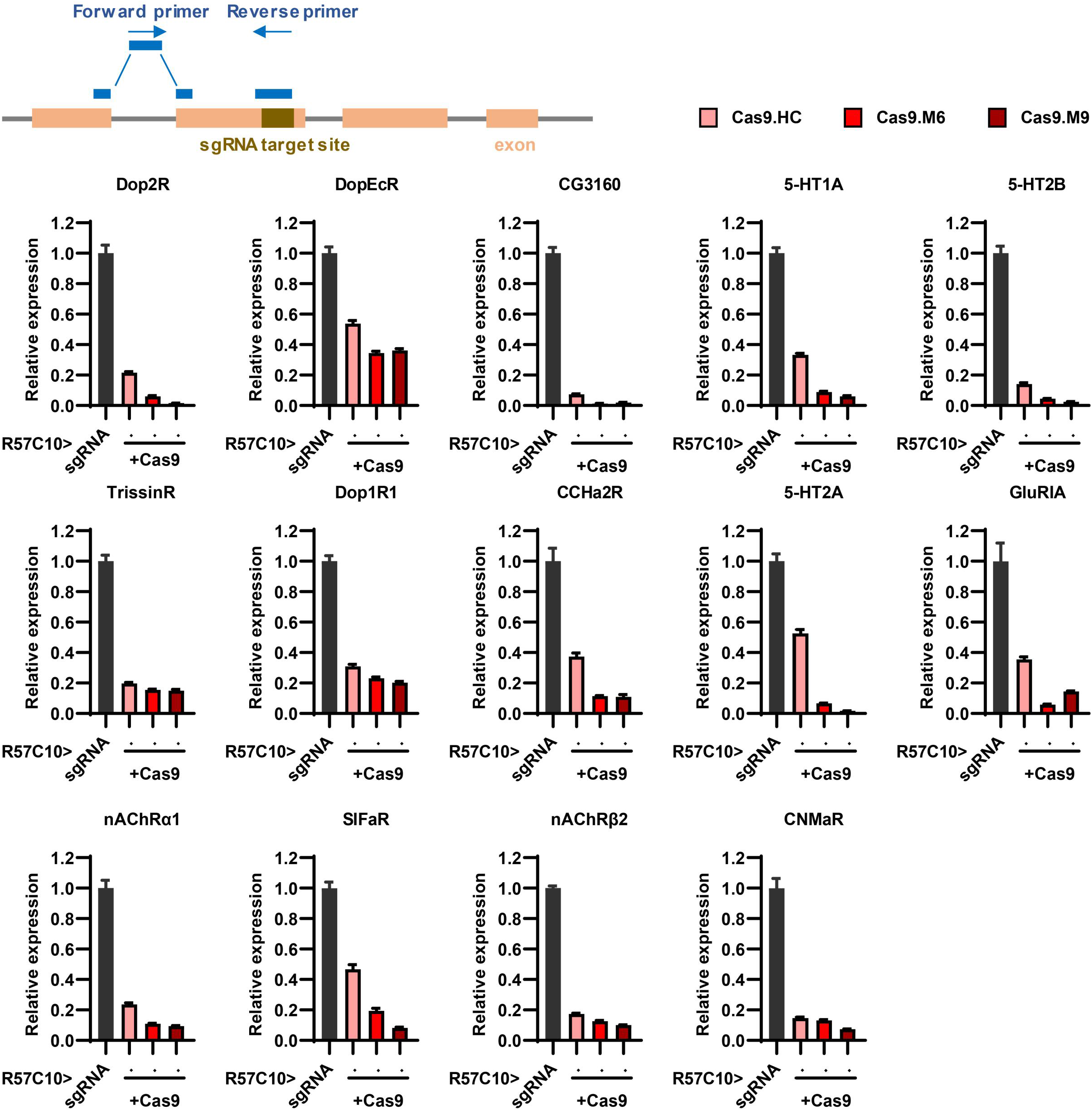
Efficiency validation by real-time quantitative PCR. This figure corresponds to Fig 3M. The schematic at the top illustrates the principle of primer design. The gene symbol above each panel indicates the target of sgRNAs. The expression of the target gene in the experimental groups (where R57C10-GAL4 drives the expression of both Cas9 variants and sgRNA) was normalized to the control group (where R57C10-GAL4 drives sgRNA expression only).

**Fig. S6.**
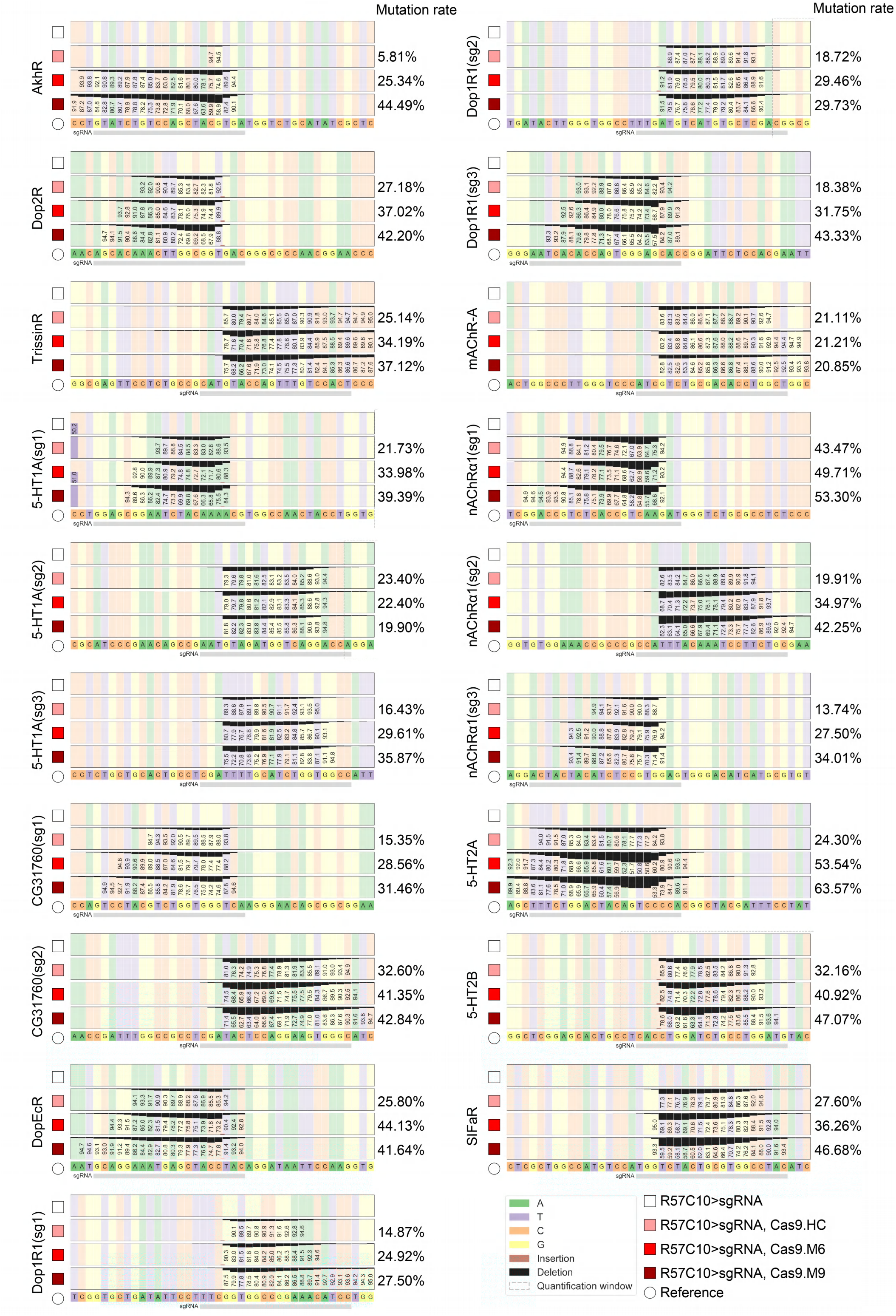
Efficiency validation by high-throughput sequencing. This figure corresponds to Fig 3P. Gene symbol on the left side of each panel denotes the target gene, while the percentages on the right denotes mutation rate as calculated by CRISPResso2. A minimum of 10,000 reads were analyzed for each genotype.

**Fig. S7.**
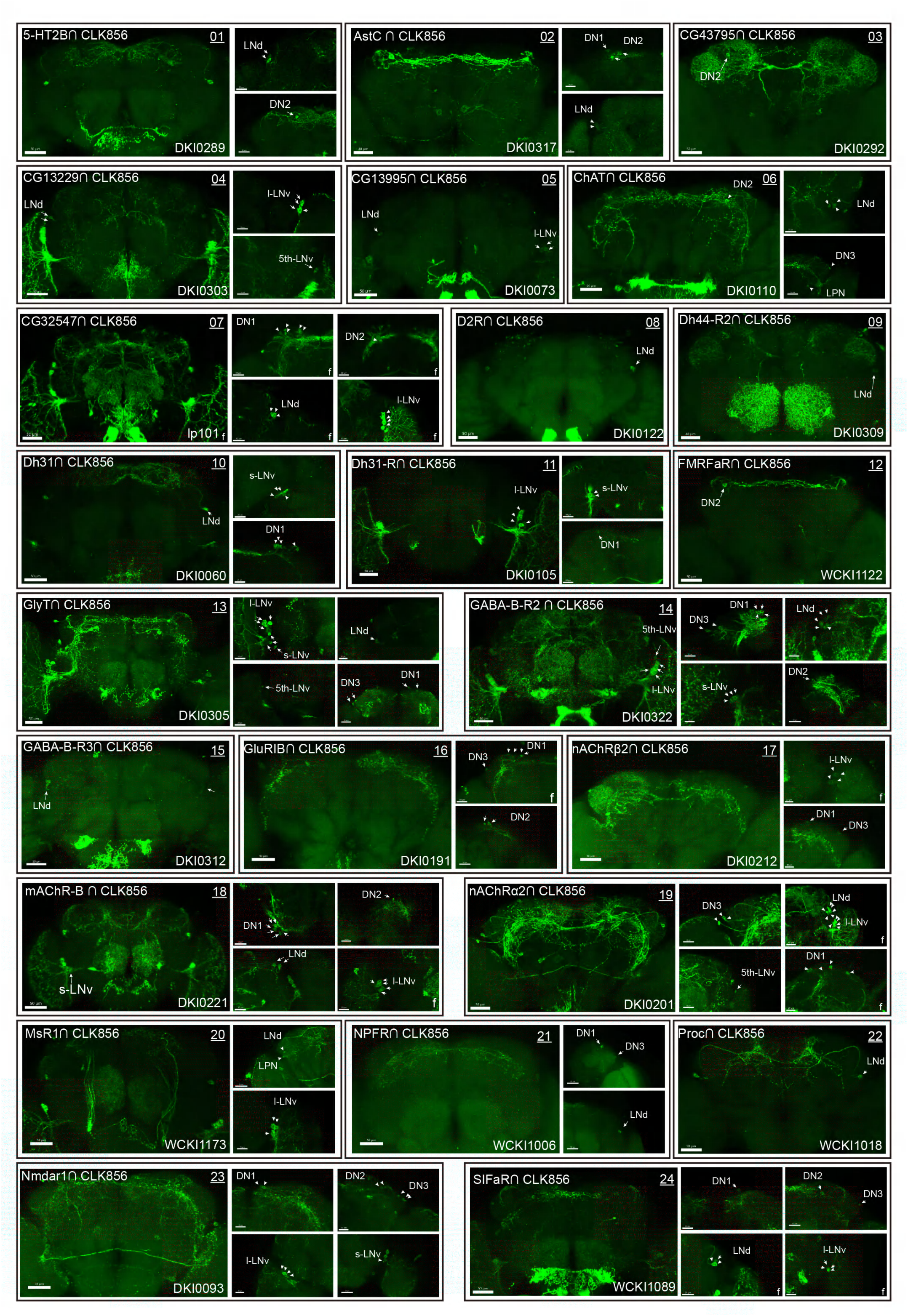
Co-expression of Clk856 with CCT Genes. (01-24) Intersectional expression patterns of CCT drivers with Clk856. The exact strategy used for each gene is annotated in Table S5. Maximum neuron numbers were presented in each image. Note at the bottom right corner is category ID of CCT drivers in Rao Lab fly stock library. “f” denotes female fly brain, otherwise male brain is shown.

**Fig. S8.**
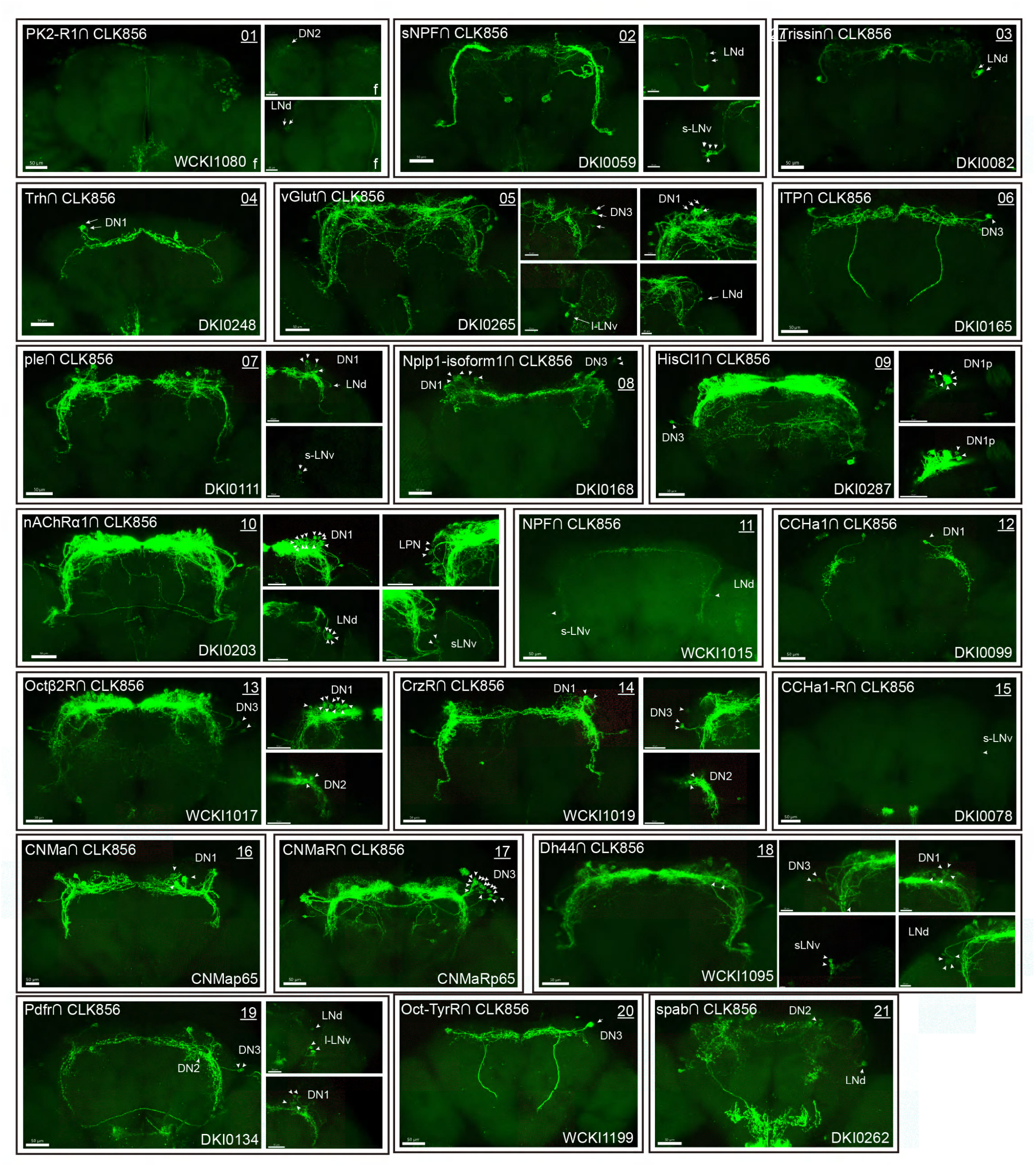
Co-expression of Clk856 with CCT Genes. (01-21) Intersectional expression patterns of CCT drivers with Clk856. The exact strategy used for each gene is annotated in Table S5. Maximum neuron numbers were presented in each image. Note at the bottom right corner is category ID of CCT drivers in Rao Lab fly stock library. “f” denotes female fly brain, otherwise male brain is shown.

**Fig. S9.**
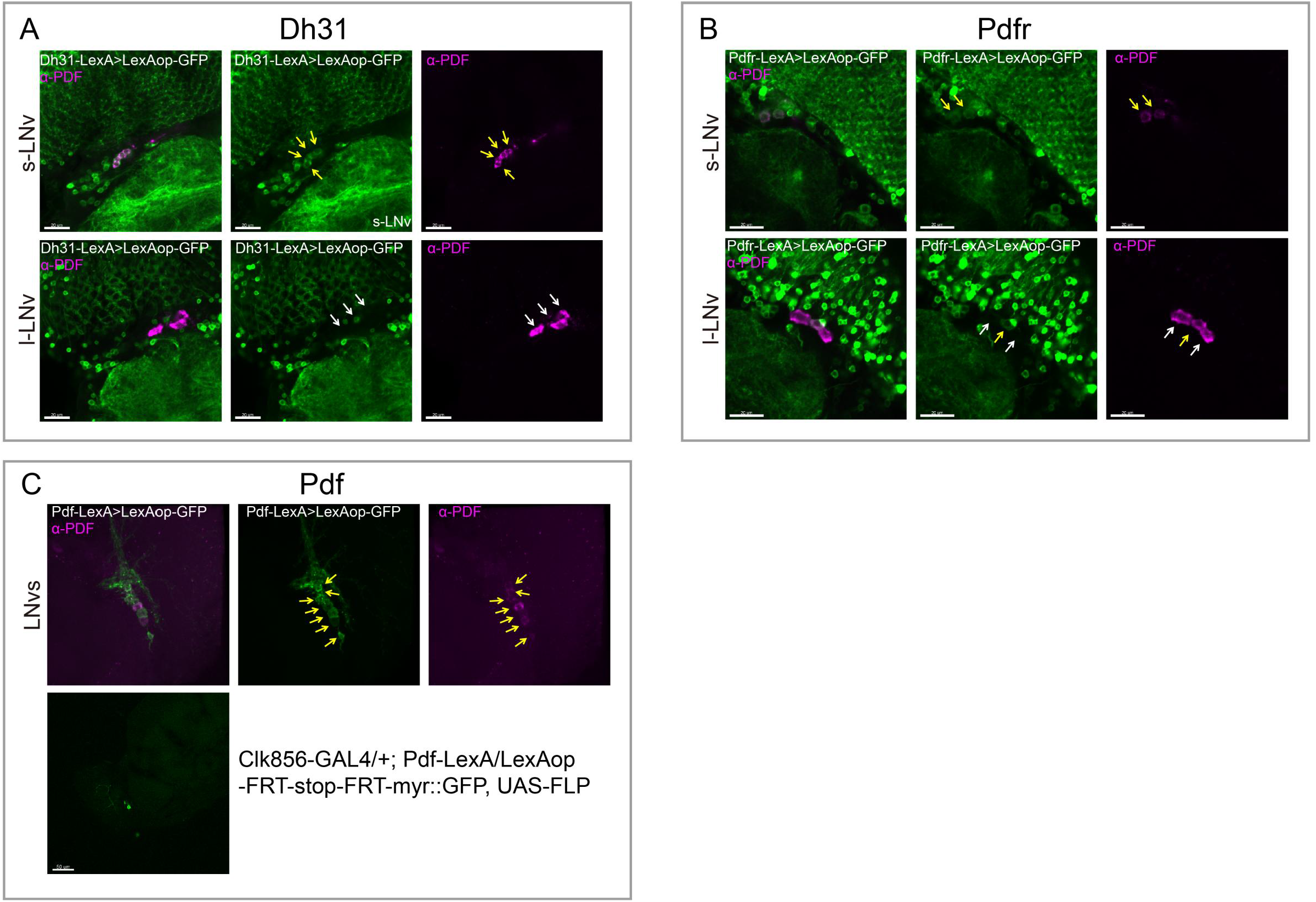
Colocalization analyse of Dh31, Pdfr, and Pdf -KI-LexA with LNvs. LNvs are labels by anti-PDF. Yellow arrow heads indicate KI-LexA (Green, A, B) colocalized with anti-PDF (violet, A, B, C) and white arrow heads indicate KI-LexA did not colocalized with anti-PDF. (C) Pdf-KI-LexA labels all PDF positive LNvs but only scattered signal observed when intersect with CLK856-GAL4.

**Fig. S10.**
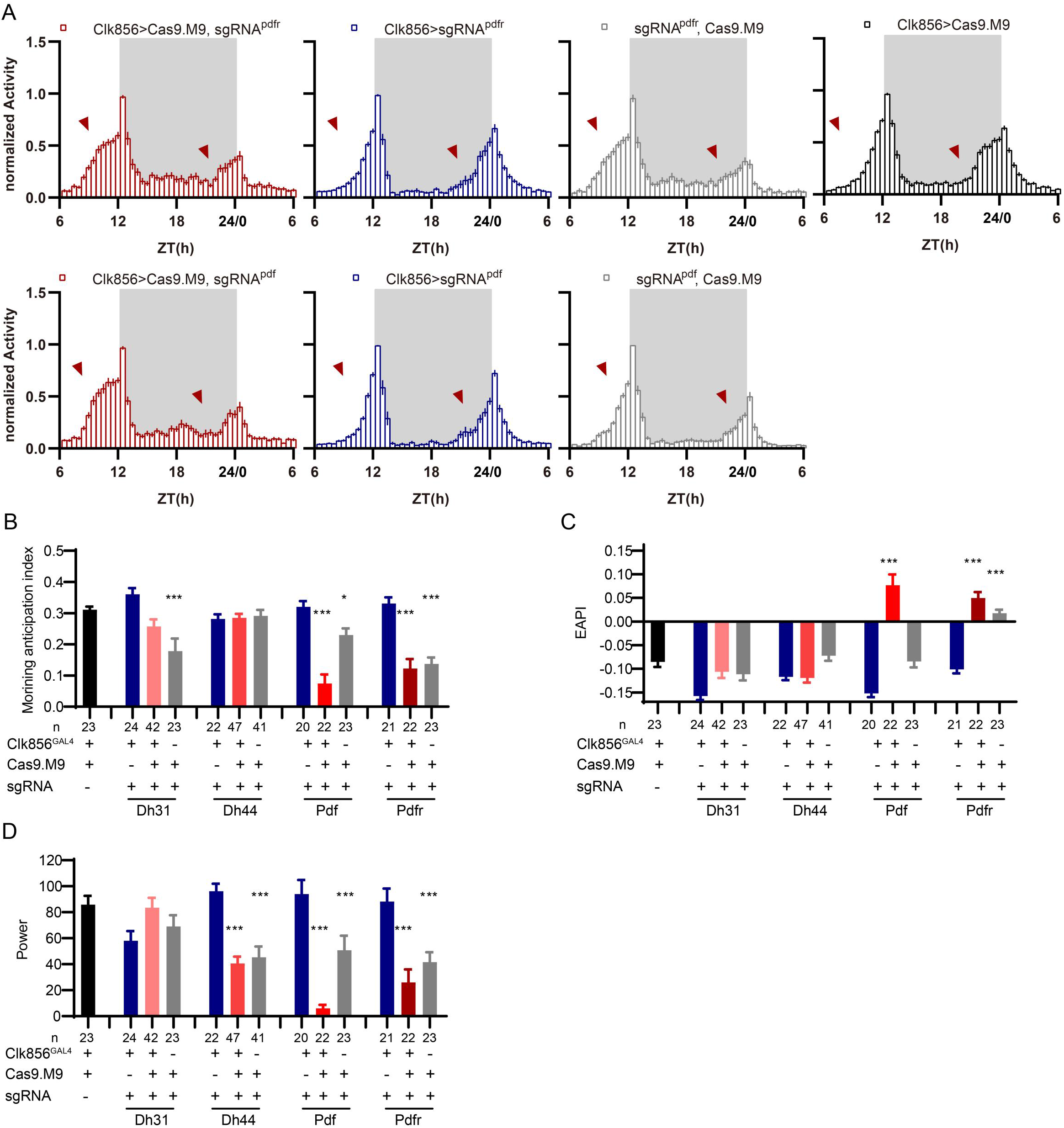
Disruption of CCT Genes due to Leaky Expression of Cas9.M9. (A) Activity plots of Pdfr or Pdf clock-neuron knockout flies (red) and control groups (blue, black, and gray). (B-D) Statistical analyses of morning anticipation index (B), EAPI (C) and power (D) after CCT genes were knockdown by Cas9.M9/sgRNAs. Leakage expression of Cas9.M9 and sgRNAs might disrupt target genes at certain levels (grey bar). Cas9.M9 denotes UAS-Cas9.M9. sgRNA denotes UAS-sgRNA.

**Fig. S11.**
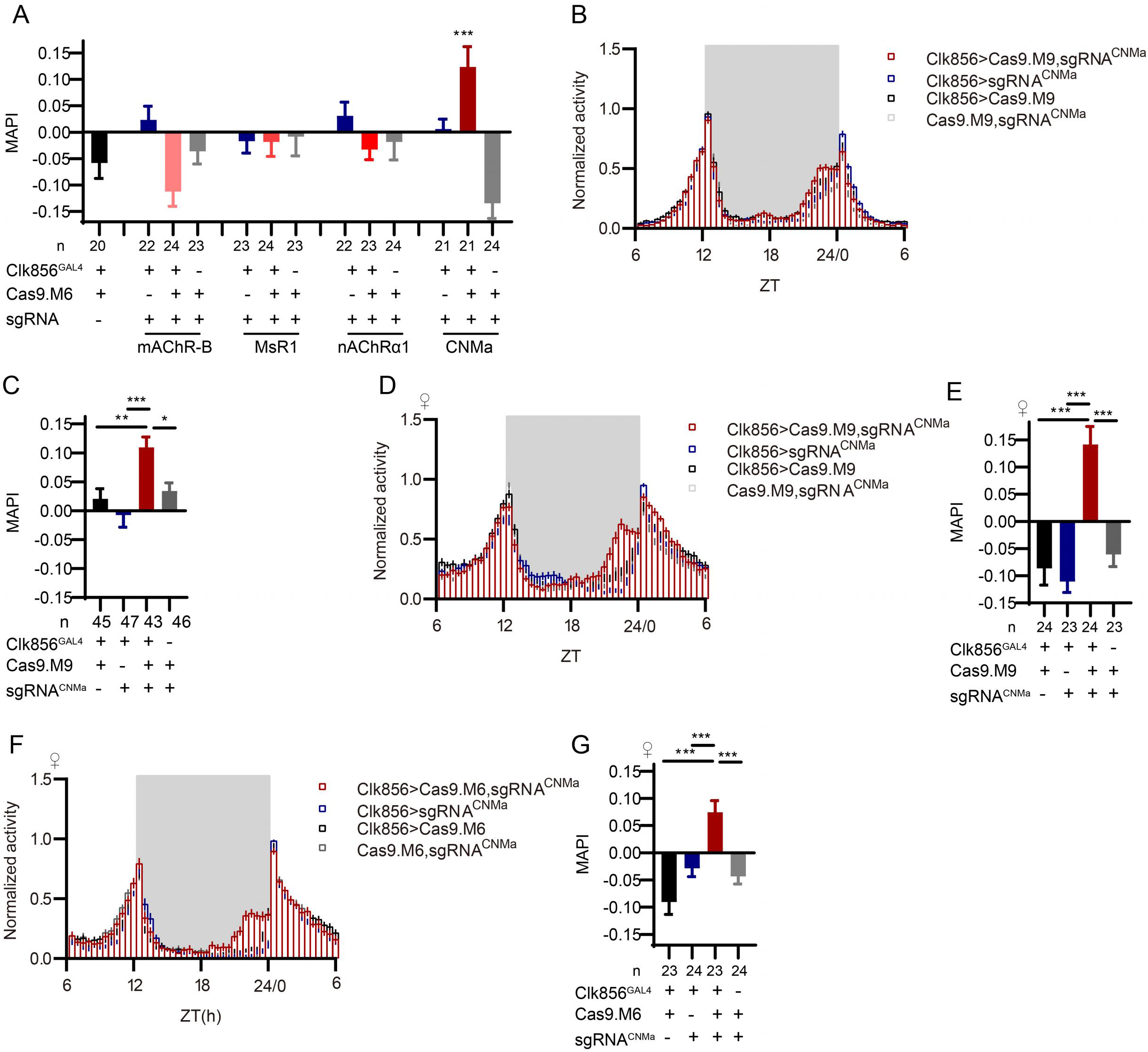
Morning Activity Advanced by Loss of CNMa. (A) MAPI statistical analyses after mAChR-B, MsR1, nAChRα1 and CNMa knockout in clock neurons. Only the CNMa knockout in clock neurons increased MAPI significantly. (B-E) Activity plots (B, D) and statistical analyses (C, E) of male (B, C) and female (D, E) flies with CNMa knockout in clock neurons. Both male and female flies showed advanced morning activity patterns B, C) and increased MAPIs (D, E). UAS-sgRNA^CNMa^/UAS-Cas9.M9 flies showed a slightly increased MAPI which indicate possible leakage expression. (F-G) Activity plots (F) and statistical analyses (G) of female flies with CNMa deficient in clock neurons with Cas9.M6 used.

**Fig. S12.**
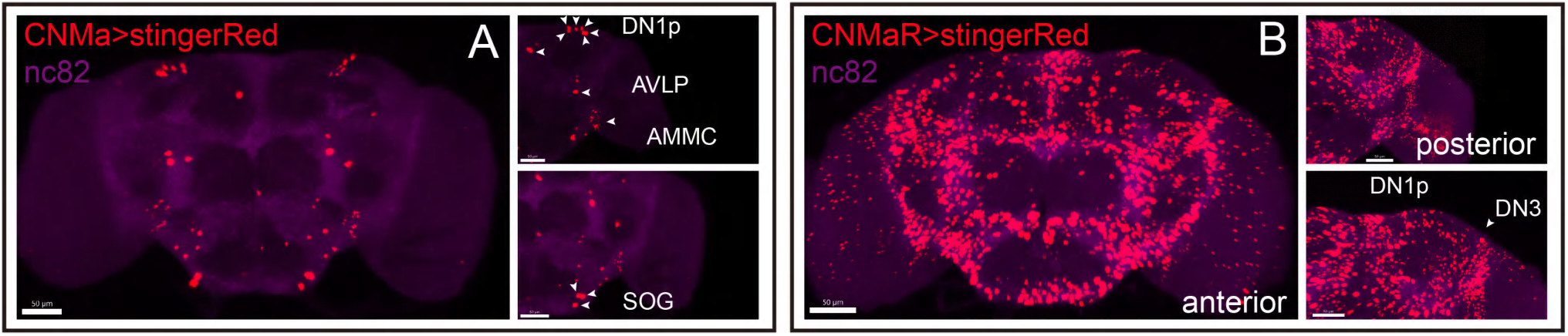
Expression of CNMa and CNMaR. (A-B) Stinger::Red labeled neurons in brain driven by CNMa-KI-GAL4 (A) and CNMaR-KI-GAL4 (B)

**Fig. S13.**
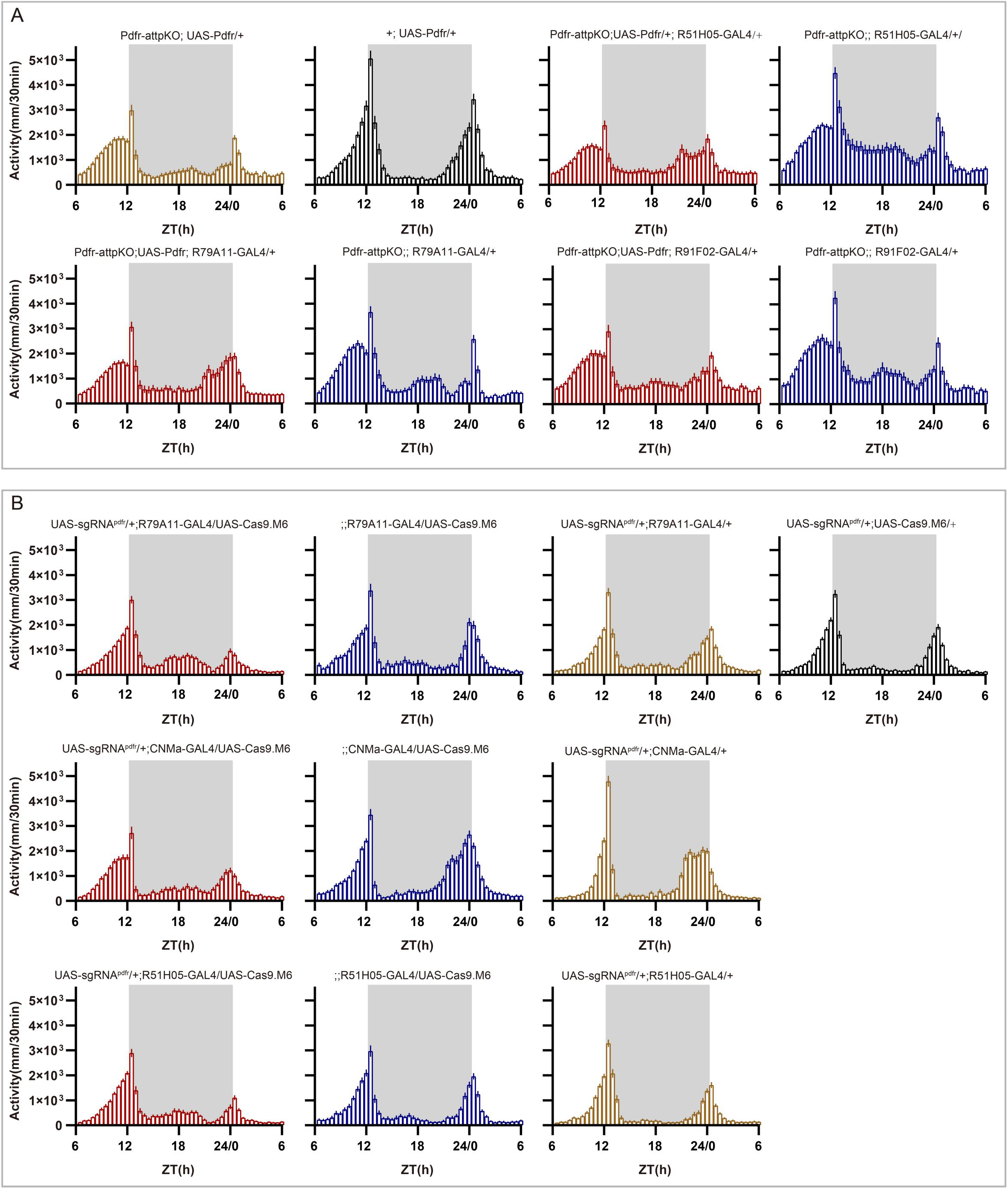
Activity plots related to. Fig 6 (A-B) Activity plots of reintroducing Pdfr (A, related to Fig 6Q) or mutating Pdfr (B, related to Fig 6T) in DN1p subset neurons. 30 min per bin is plotted.

**Fig. S14.**
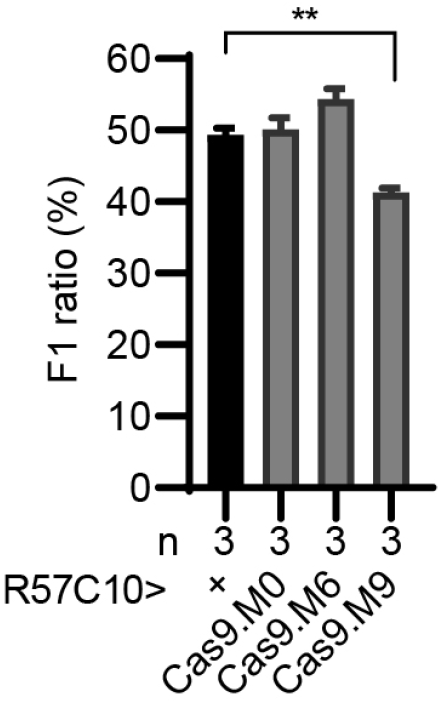
Impact on Viability of Cas9 Variants Expression by GMR57C10-GAL4. F1 ratio of UAS-Cas9 variants (male) cross GMR57C10-GAL4/cyo (virgin female). Expression of Cas9.M9 decreased progeny viability slightly. ** *p<0.01*. Two-student test was used.

**Table S1.**
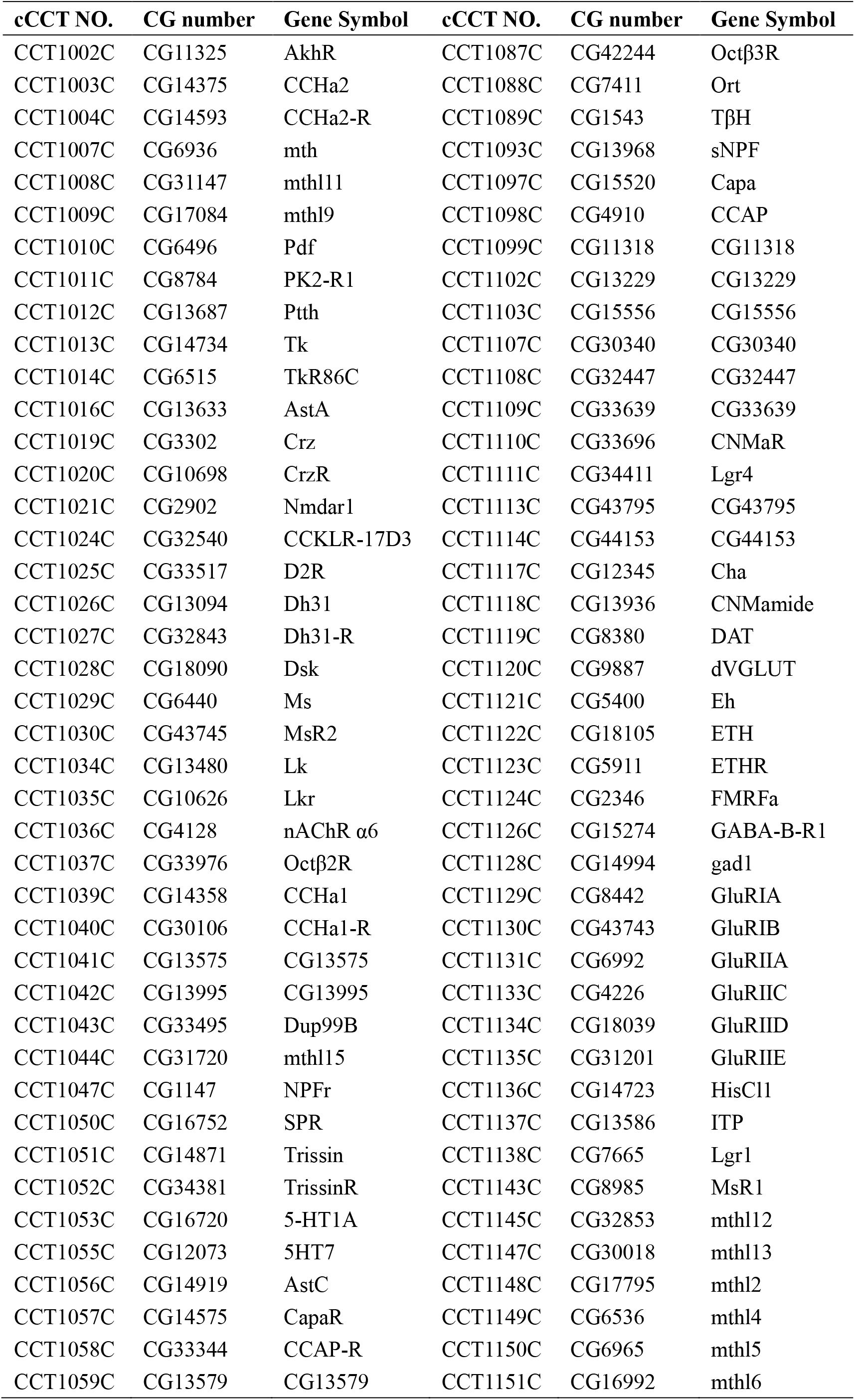
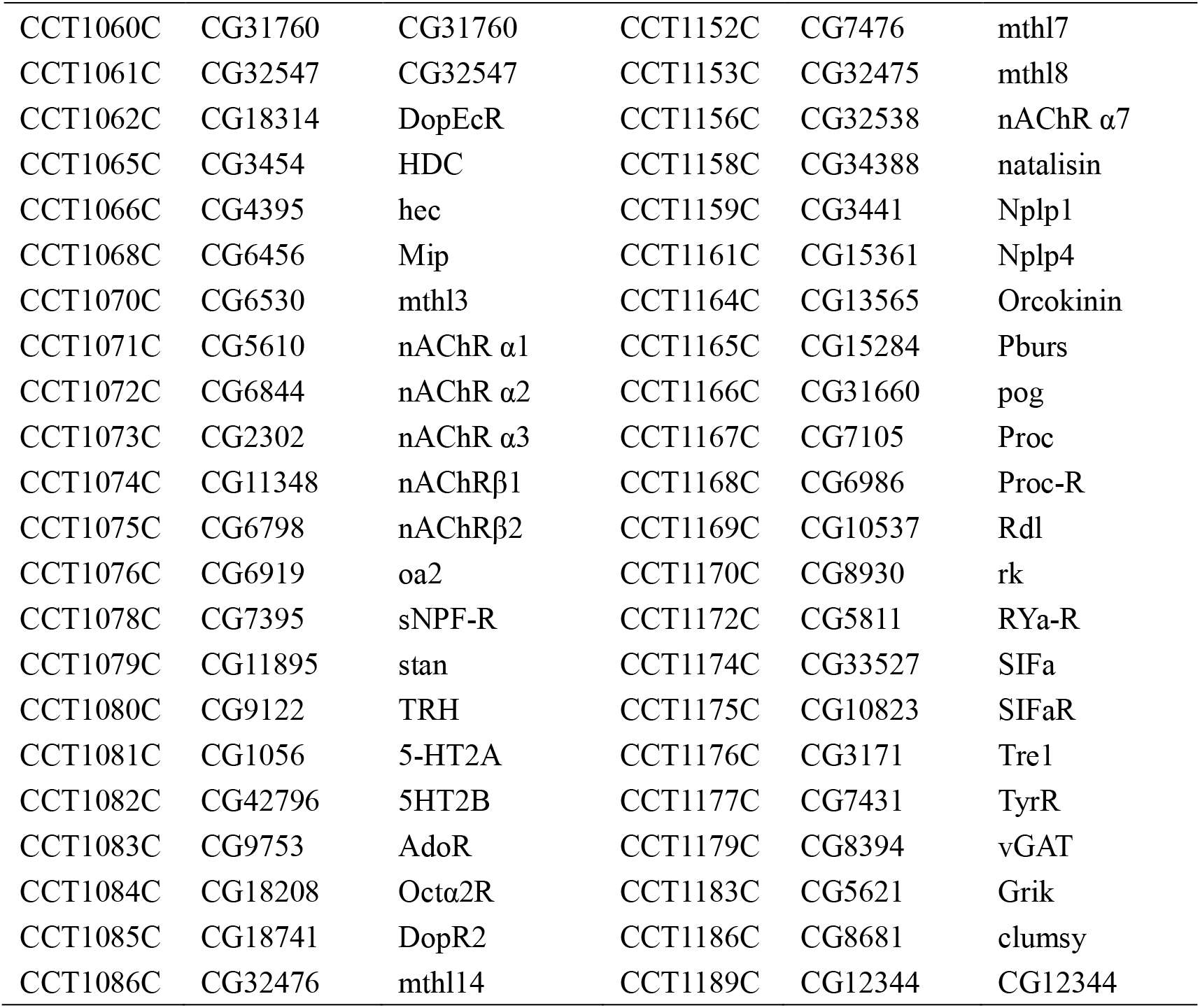
cCCT Knockin fly list.

**Table S2.**
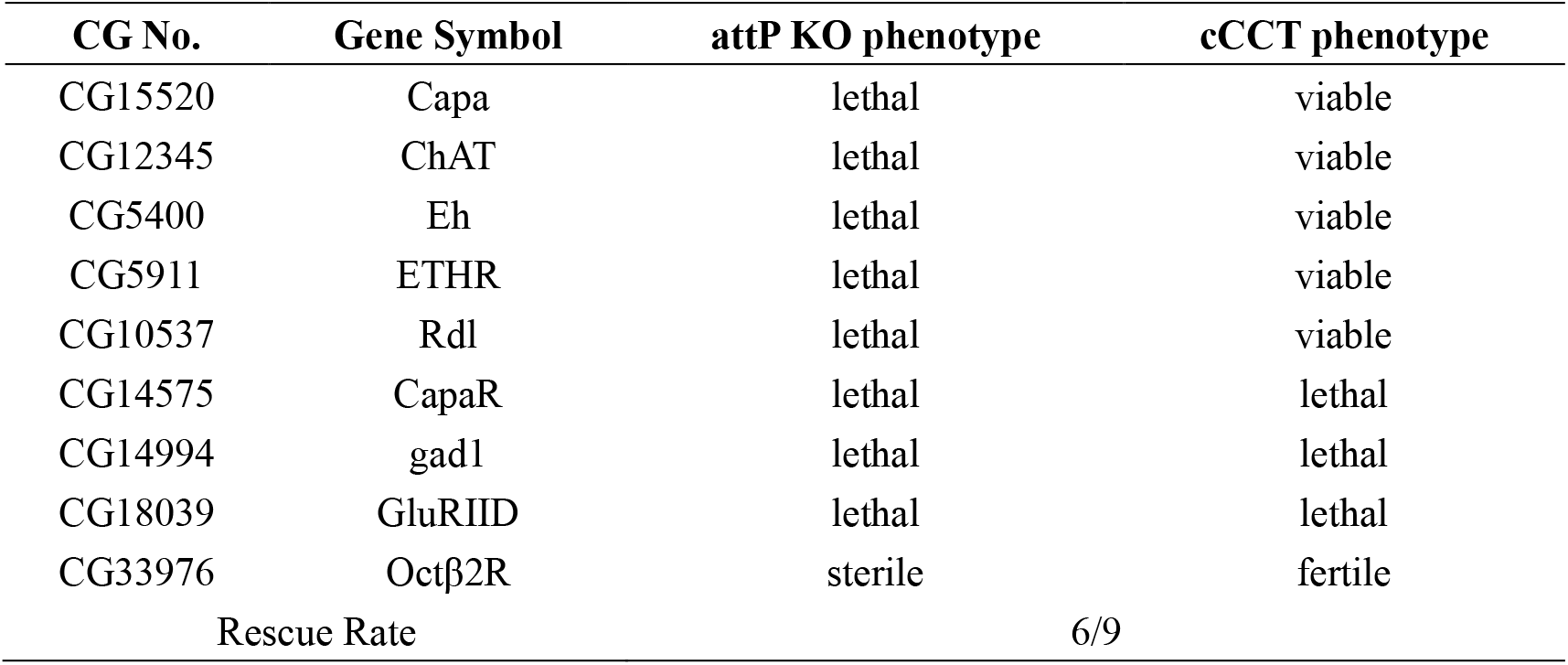
cCCT knockin strategy can’t rescue all knockout phenotype.

| **CG No.** | **Gene Symbol** | **attP KO phenotype** | **cCCT phenotype** |
| --- | --- | --- | --- |
| CG15520 | Capa | lethal | viable |
| CG12345 | ChAT | lethal | viable |
| CG5400 | Eh | lethal | viable |
| CG5911 | ETHR | lethal | viable |
| CG10537 | Rdl | lethal | viable |
| CG14575 | CapaR | lethal | lethal |
| CG14994 | gad1 | lethal | lethal |
| CG18039 | GluRIID | lethal | lethal |
| CG33976 | Octβ2R | sterile | fertile |
| Rescue Rate |  | 6/9 |  |

**Table S3.**
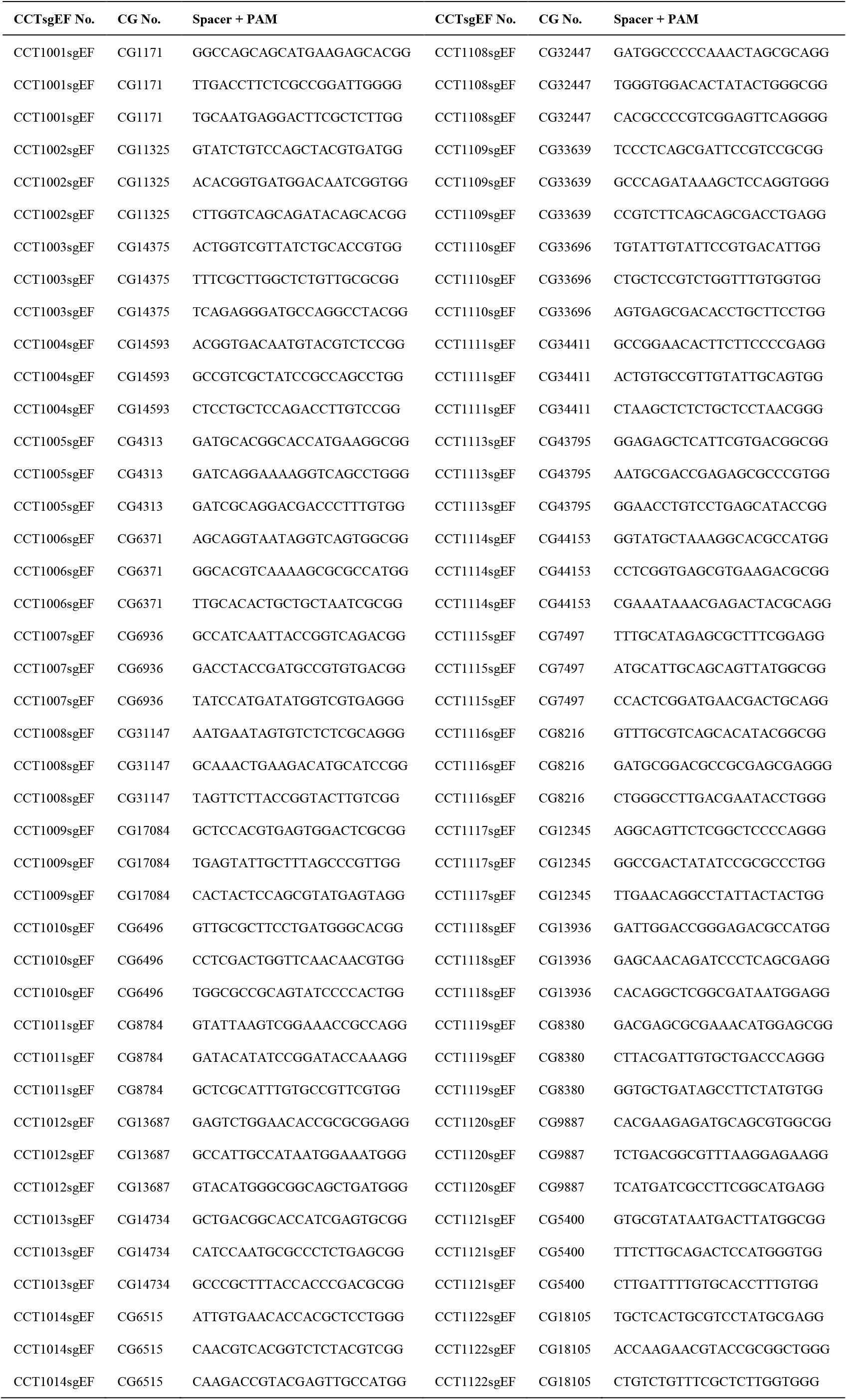
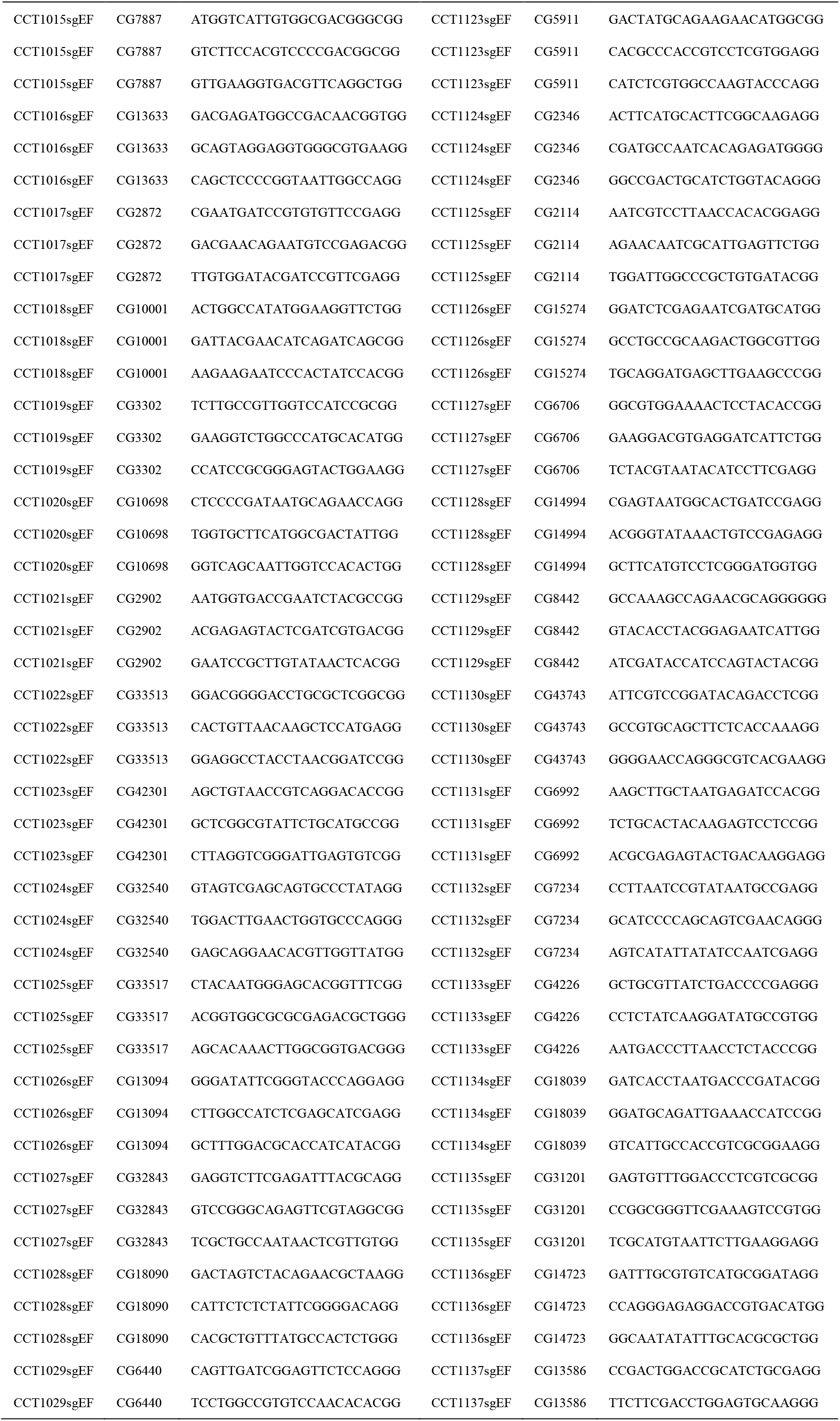
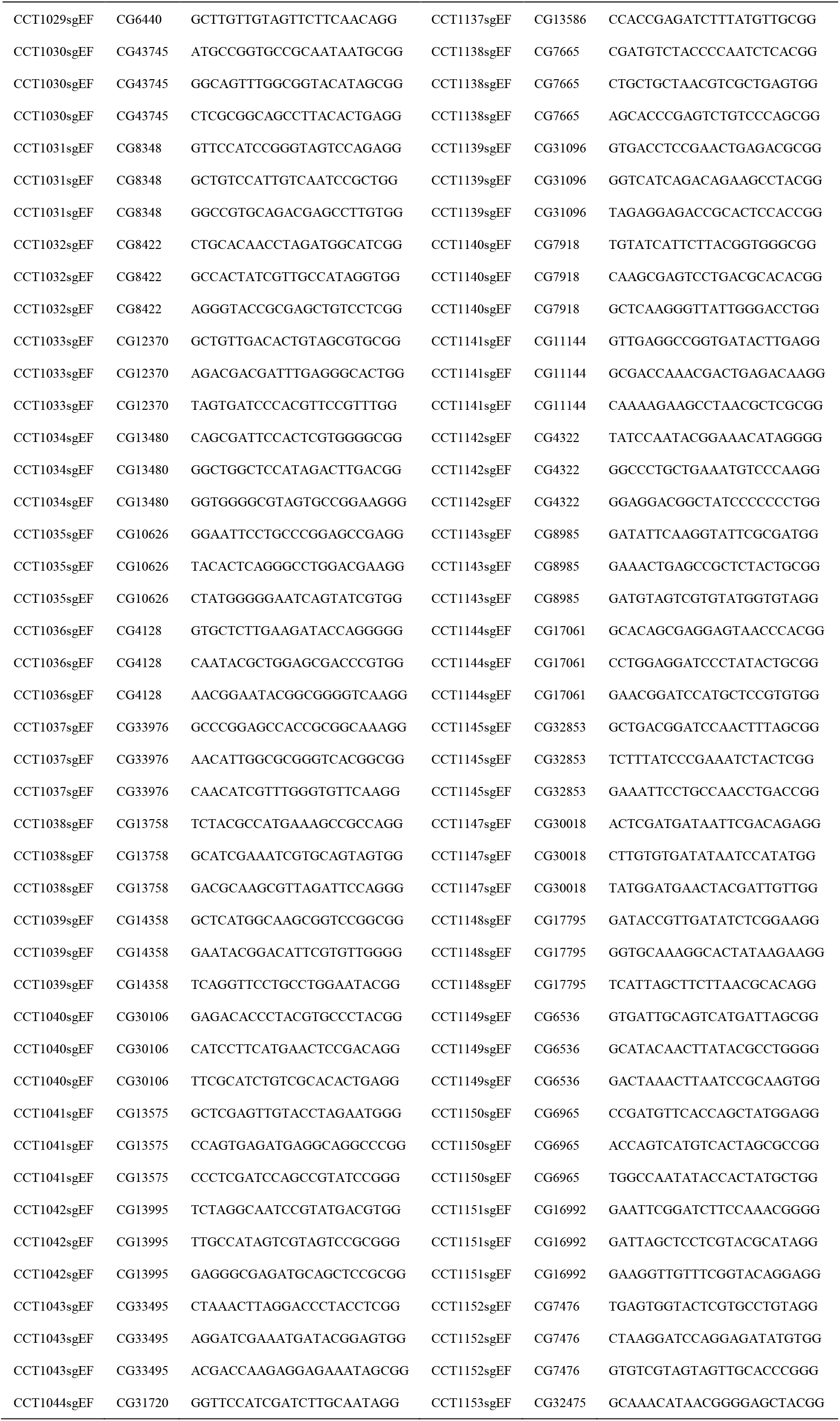
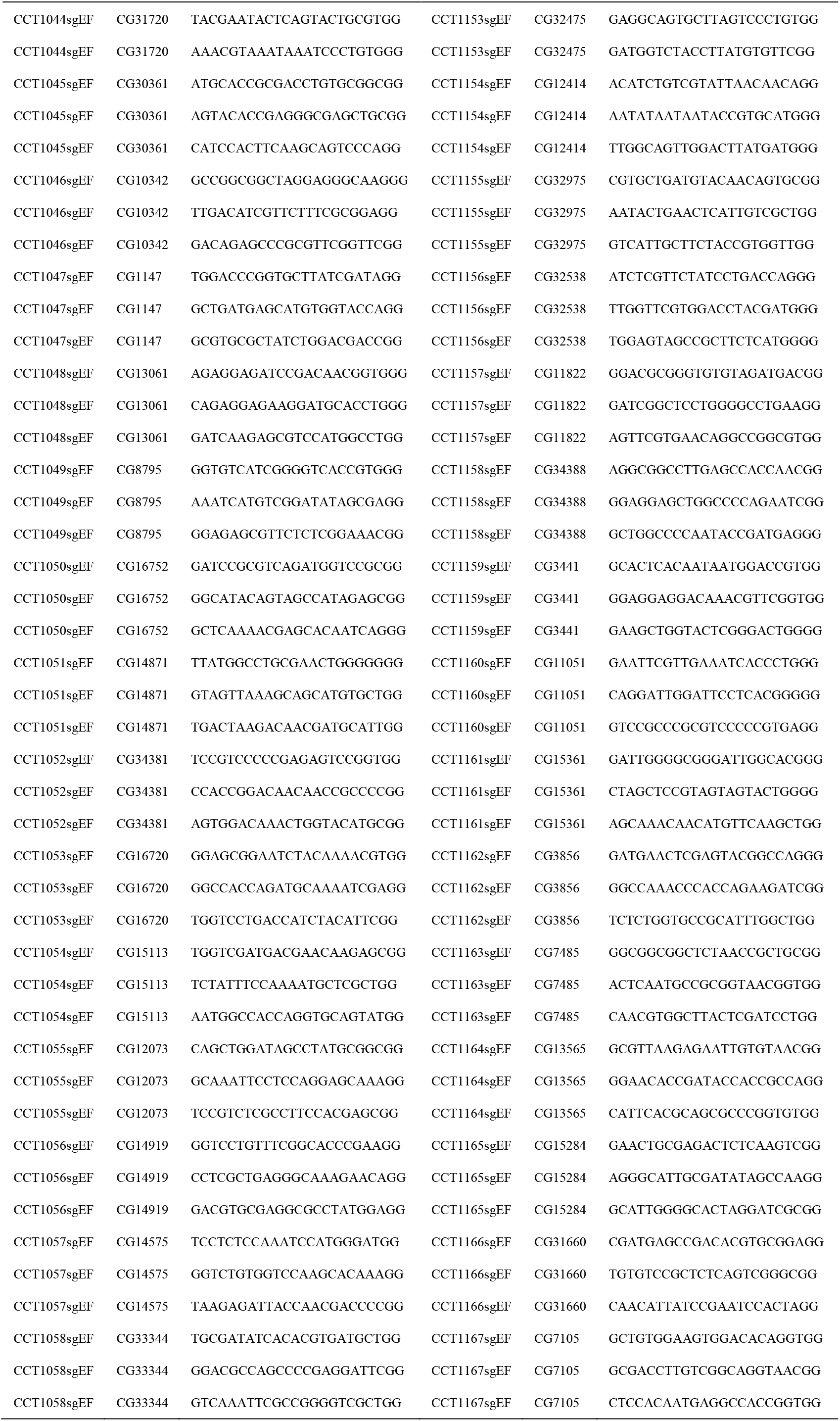
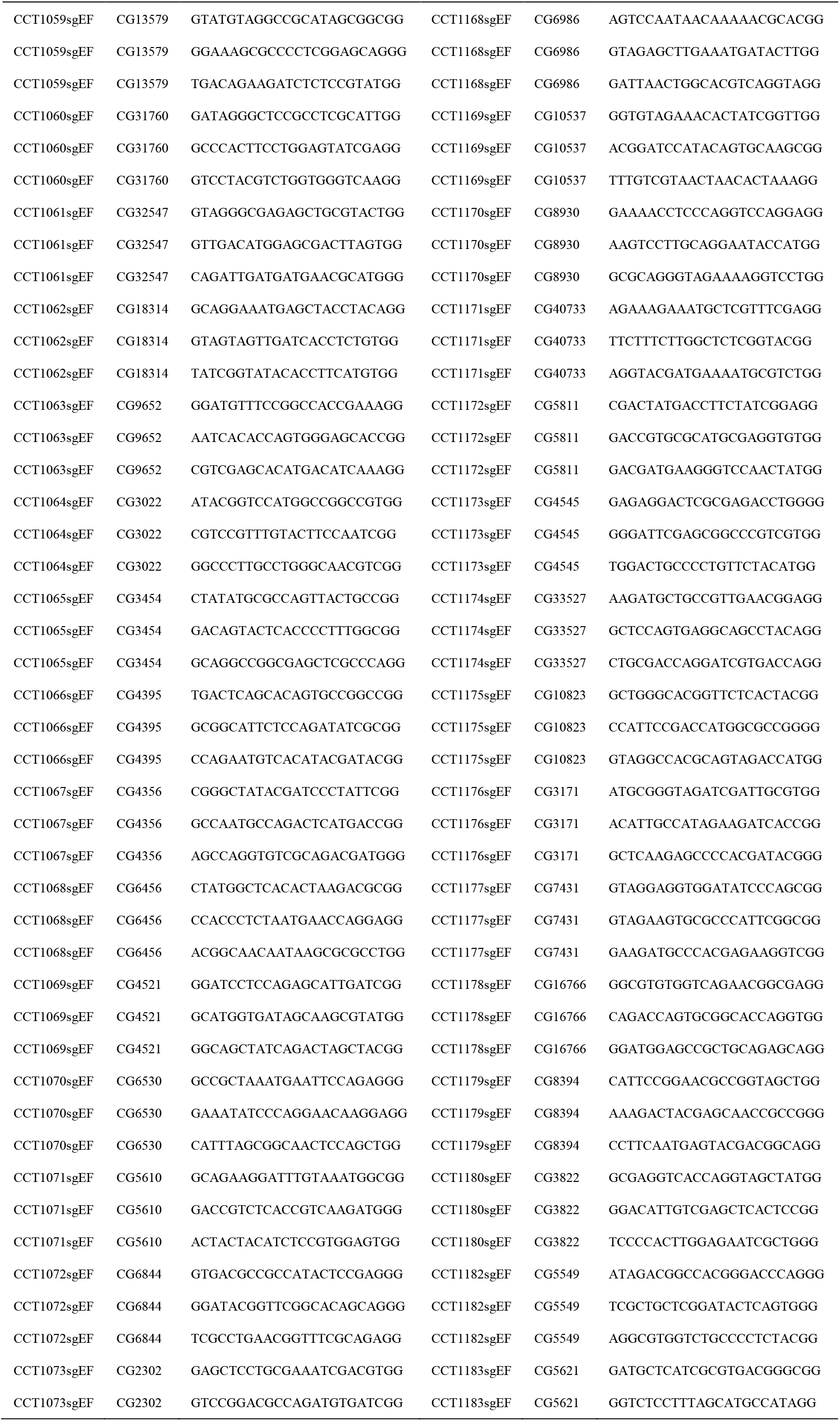
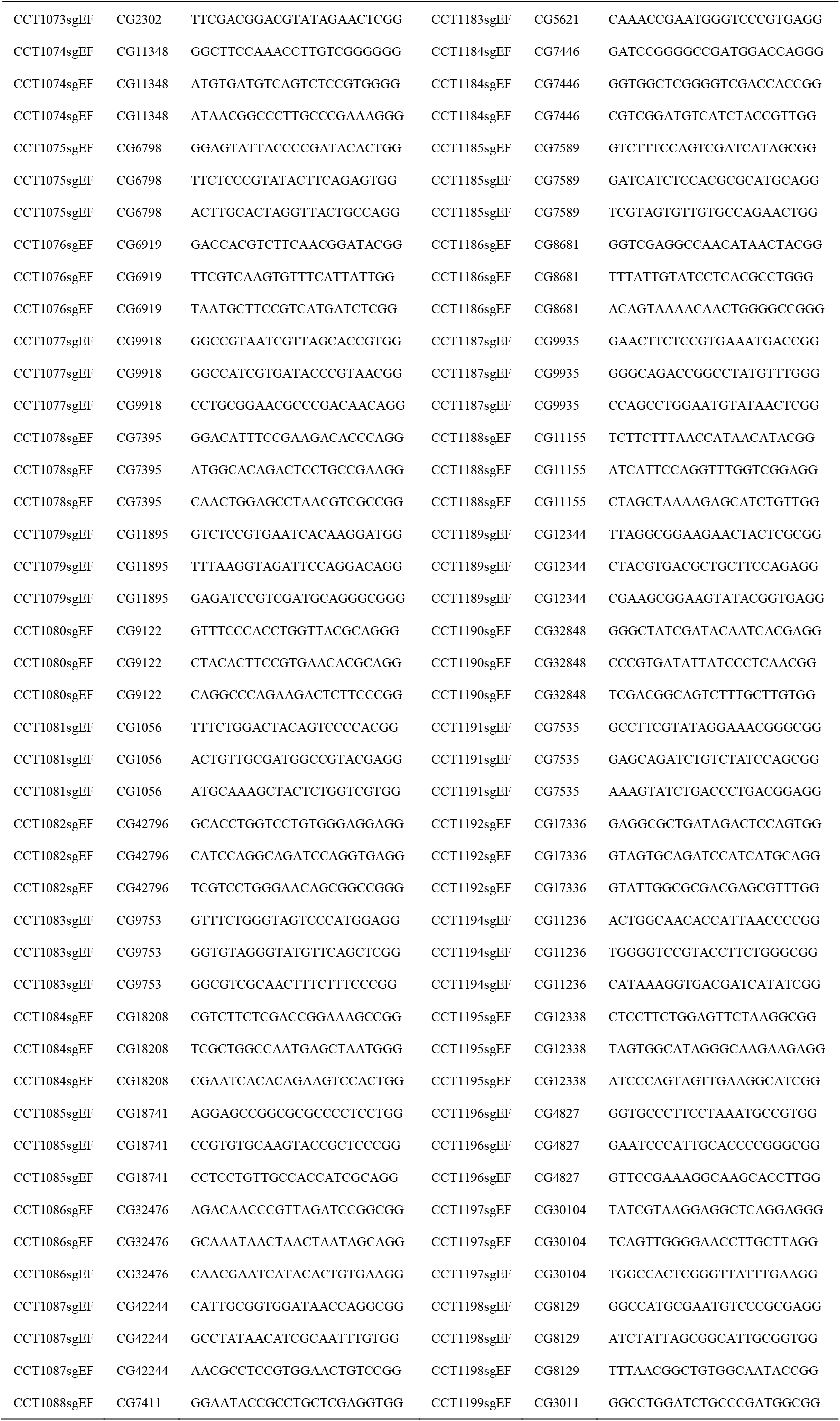
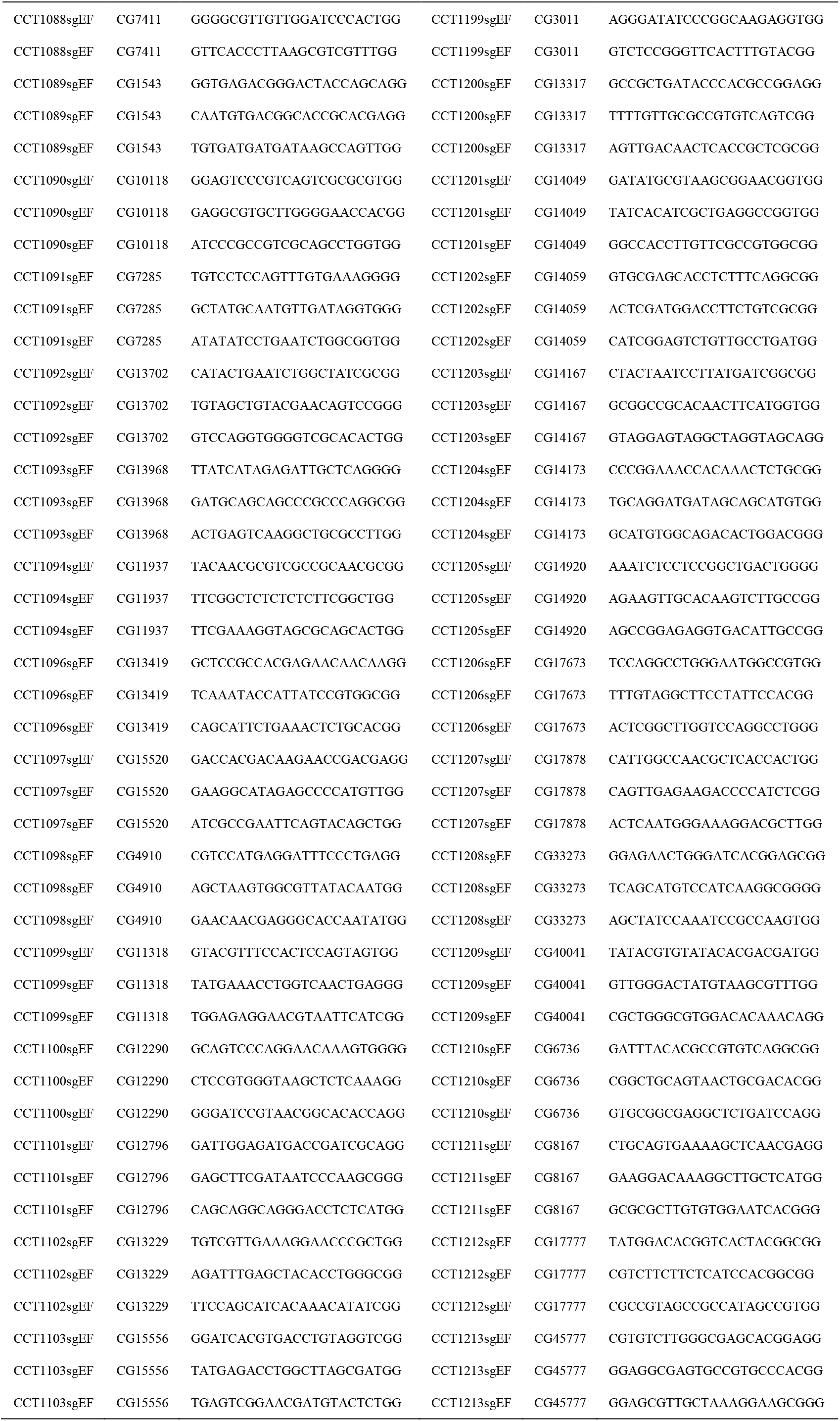
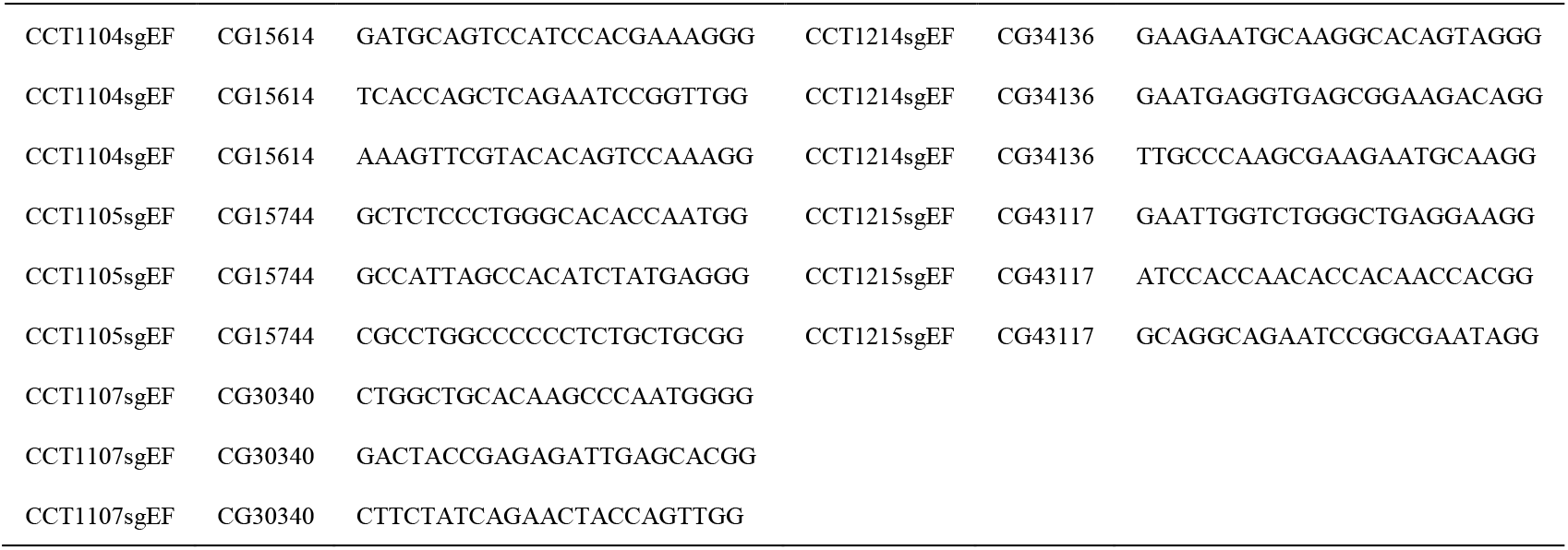
C-cCCTomics sgRNAs list.

**Table S4.** CCT gene expression profile of clock neuron.

|  | s-LNV |  | l-LNV |  | 5th |  | LNd |  | LPN |  | DN1 |  | DN2 |  | DN3 |  |
| --- | --- | --- | --- | --- | --- | --- | --- | --- | --- | --- | --- | --- | --- | --- | --- | --- |
| Gene symbol | GFP+ | N<sup>#</sup> | GFP+ | N<sup>#</sup> | GFP+ | N<sup>#</sup> | GFP+ | N<sup>#</sup> | GFP+ | N<sup>#</sup> | GFP+ | N<sup>#</sup> | GFP+ | N<sup>#</sup> | GFP+ | N<sup>#</sup> |
| AstC | / | / | / | / | / | / | 1 ± 0 | 6 | / | / | / | / | 1.83 ± 0.41 | 6 | 1 ± 0 | 1 |
| CCHa1 | / | / | / | / | / | / | / | / | / | / | 1.6 ± 0.55 | 5 | / | / | / | / |
| ChAT | 3 ± 1.41 | 2 | / | / | / | / | 2.3 ± 1.06 | 10 | 1 ± 1 | 1 | 1 ± 0 | 1 | 1 ± 0 | 10 | 1 ± 0 | 7 |
| CNMa | / | / | / | / | / | / | / | / | / | / | 4 ± 1.41 | 2 | / | / | 1 ± 0 | 2 |
| Dh31 | 3.64 ± 0.5 | 11 | / | / | / | / | 1.2 ± 0.42 | 10 | / | / | 3.8 ± 0.63 | 10 | / | / | / | / |
| Dh44 | / | / | 1.67 ± 0.58 | 3 | / | / | 3 ± 1 | 3 | / | / | 5 ± 0 | 3 | 1.67 ± 0.58 | 3 | 1.67 ± 0.58 | 3 |
| GlyT | 2.8 ± 1.64 | 5 | 3.88 ± 0.35 | 8 | 1 ± 0 | 3 | 1 ± 0 | 3 | / | / | 1.57 ± 0.79 | 7 | 1 ± 0 | 6 | 2 ± 0.93 | 8 |
| ITP | / | / | / | / | / | / | / | / | / | / | / | / | / | / | 1 ± 0 | 2 |
| NPF | 1 ± 0 | 2 | / | / | / | / | 1 ± 0 | 3 | / | / | / | / | / | / | / | / |
| Nplp1 | / | / | / | / | / | / | / | / | / | / | 5 ± 0 | 2 | / | / | 2 ± 0 | 2 |
| Proc | / | / | / | / | / | / | 1 ± 0 | 12 | / | / | / | / | / | / | / | / |
| sNPF | 4 ± 0 | 7 | / | / | / | / | 1.63 ± 0.52 | 8 | / | / | / | / | / | / | / | / |
| TH | 1.5 ± 0.71 | 2 | / | / | / | / | 1 ± 0 | 2 | / | / | 3.5 ± 0.71 | 2 | / | / | / | / |
| Trh | / | / | / | / | / | / | / | / | / | / | 1.75 ± 0.5 | 4 | / | / | / | / |
| Trissin | / | / | / | / | / | / | 1.82 ± 0.4 | 11 | / | / | / | / | / | / | / | / |
| VGlut | / | / | 1.2 ± 0.45 | 5 | / | / | 1.17 ± 0.41 | 6 | / | / | 3.5 ± 0.55 | 6 | 2 ± 0 | 1 | 2.2 ± 0.45 | 5 |
| CG13229 | / | / | 3.5 ± 0.55 | 6 | 1 ± 0 | 1 | 1.5 ± 0.55 | 6 | / | / | / | / | / | / | 1 ± 0 | 3 |
| CG32547 | / | / | 2.71 ± 0.95 | 7 | / | / | 1.86 ± 0.9 | 7 | / | / | 3.29 ± 1.8 | 7 | 1 ± 0 | 5 | / | / |
| CG43795 | / | / | / | / | / | / | / | / | / | / | / | / | 1 ± 0 | 6 | / | / |
| 5-HT2B | / | / | / | / | / | / | 1.62 ± 0.65 | 13 | / | / | / | / | 1 ± 0 | 1 | / | / |
| CCHa1-R | 1.5 ± 0.71 | 2 | / | / | / | / | / | / | / | / | / | / | / | / | / | / |
| CNMaR | / | / | / | / | / | / | / | / | / | / | / | / | / | / | 15.67 ± 0.58 | 3 |
| CrzR | / | / | / | / | / | / | / | / | / | / | 2 ± 0 | 2 | 1.5 ± 0.71 | 2 | 2.5 ± 0.71 | 2 |
| D2R | / | / | / | / | / | / | 1 ± 0 | 6 | / | / | / | / | / | / | / | / |
| Dh31-R | 2 ± 0 | 2 | 3.43 ± 0.53 | 7 | 1 ± 0 | 2 | / | / | / | / | 1 ± 0 | 3 | / | / | / | / |
| Dh44-R2 | / | / | / | / | / | / | 1 ± 0 | 6 | / | / | / | / | / | / | / | / |
| FMRFaR | / | / | / | / | / | / | / | / | / | / | / | / | 1 ± 0 | 6 | / | / |
| GABA-B-R2 | 3.43 ± 0.53 | 7 | 3.63 ± 0.52 | 8 | 1 ± 0 | 4 | 4.29 ± 1.7 | 7 | / | / | 3 ± 0 | 6 | 1 ± 0 | 1 | 1 ± 0 | 1 |
| GABA-B-R3 | / | / | / | / | / | / | 1 ± 0 | 4 | / | / | / | / | / | / | / | / |
| GluRIB | / | / | / | / | / | / | / | / | / | / | **1.56 ± 1.01** | 9 | 1.5 ± 0.58 | 4 | *1 ± 0* | 2 |
| mAChR-B | 1.33 ± 0.58 | 3 | 3.33 ± 0.87 | 9 | / | / | 1.5 ± 0.53 | 8 | / | / | 3.38 ± 2.33 | 8 | 1 ± 0 | 5 | / | / |
| MsR1 | / | / | **3.29 ± 0.49** | 7 | / | / | **1 ± 0** | 7 | *1 ± 1* | *1* | / | / | 2 ± 0 | 1 | / | / |
| nAChRα1 | 3 ± 0 | 2 | / | / | / | / | **5.6 ± 0.89** | 5 | 2 ± 1 | 3 | **12.4 ± 1.52** | 5 | / | / | 1 ± 0 | 2 |
| nAChRα2 | / | / | 3.8 ± 0.45 | 5 | *1 ± 0* | *1* | 2 ± 0.89 | 6 | / | / | 2.17 ± 1.17 | 6 | / | / | 2 ± 0.71 | 5 |
| nAChRβ2 | / | / | 3 ± 1 | 3 | / | / | / | / | / | / | 1 ± 0 | 2 | / | / | 1 ± 0 | 5 |
| Nmdar1 | *1 ± 0* | 2 | 3.64 ± 0.5 | 11 | / | / | 1.88 ± 0.35 | 8 | / | / | 1.5 ± 0.58 | 4 | 1.11 ± 0.33 | 9 | 1.4 ± 0.84 | 10 |
| NPFR | / | / | / | / | / | / | 1 ± 0 | 10 | / | / | / | / | / | / | **1.14 ± 0.38** | 7 |
| Oct-TyrR | / | / | / | / | / | / | / | / | / | / | / | / | / | / | 2 ± 0 | 2 |
| Octβ2R | / | / | / | / | / | / | / | / | / | / | **10.5 ± 0.71** | 2 | **1.5 ± 0.71** | 2 | 2.5 ± 0.71 | 2 |
| Pdfr | / | / | 1.14 ± 0.38 | 7 | / | / | *1 ± 0* | 2 | / | / | **2.29 ± 1.11** | 7 | 1 ± 0 | 6 | 1.33 ± 0.58 | 3 |
| PK2-R1 | / | / | / | / | / | / | **1.5 ± 0.71** | 2 | / | / | / | / | 1 ± 0 | 3 | / | / |
| SIFaR | / | / | **3.21 ± 0.43** | 14 | / | / | **2 ± 0.39** | 14 | / | / | **1 ± 0** | 5 | 1 ± 0 | 6 | **1.22 ± 0.44** | 9 |
| spab | / | / | / | / | / | / | 1 ± 0 | 4 | / | / | / | / | 1 ± 0 | 2 | / | / |

N# represents GFP positive brain number. The largest N# number indicates total brains analyzed for certain CCT line.
/ represents no GFP found in corresponding clock neuron subset
The *italic* values indicate that the number of positive brains is less than 1/3 of the total analyzed brains
The **bolder** values (41/127) indicate that genes were also reported in transcriptome analysis (Ma et al., 2021)

**Table S5.** List of CCT genes intersected with Clk856 drivers.

| **RaoLab Stock No.** | **CG No.** | **Gene Symbol** | **Driver** | **Intersection with Clk856** |
| --- | --- | --- | --- | --- |
| DKI0289 | CG42796 | 5-HT2B | LexA | Clk856-GAL4 |
| DKI0317 | CG14919 | AstC | LexA | Clk856-GAL4 |
| DKI0292 | CG43795 | CG43795 | LexA | Clk856-GAL4 |
| DKI0303 | CG13229 | CG13229 | LexA | Clk856-GAL4 |
| DKI0073 | CG13995 | CG13995 | LexA | Clk856-GAL4 |
| DKI0110 | CG12345 | ChAT | LexA | Clk856-GAL4 |
| DKI0254 | CG32547 | CG32547 | LexA | Clk856-GAL4 |
| DKI0121 | CG33517 | Dop2R | LexA | Clk856-GAL4 |
| DKI0309 | CG12370 | Dh44-R2 | LexA | Clk856-GAL4 |
| DKI0060 | CG13094 | Dh31 | LexA | Clk856-GAL4 |
| DKI0105 | CG32843 | Dh31-R | LexA | Clk856-GAL4 |
| WCKI1122 | CG2114 | FMRFaR | LexA | Clk856-GAL4 |
| DKI0305 | CG5549 | GlyT | LexA | Clk856-GAL4 |
| DKI0322 | CG6706 | GABA-B-R2 | LexA | Clk856-GAL4 |
| DKI0312 | CG3022 | GABA-B-R3 | LexA | Clk856-GAL4 |
| DKI0191 | CG43743 | GluRIB | LexA | Clk856-GAL4 |
| DKI0212 | CG6798 | nAChRβ2 | LexA | Clk856-GAL4 |
| DKI0221 | CG7918 | mAChR-B | LexA | Clk856-GAL4 |
| DKI0201 | CG6844 | nAChRα2 | LexA | Clk856-GAL4 |
| WCKI1173 | CG8985 | MsR1 | LexA | Clk856-GAL4 |
| WCKI1006 | CG1147 | NPFR | LexA | Clk856-GAL4 |
| WCKI1018 | CG7105 | Proc | LexA | Clk856-GAL4 |
| DKI0093 | CG2902 | Nmdar1 | LexA | Clk856-GAL4 |
| WCKI1089 | CG10823 | SIFaR | LexA | Clk856-GAL4 |
| WCKI1080 | CG8784 | PK2-R1 | LexA | Clk856-GAL4 |
| DKI0059 | CG13968 | sNPF | LexA | Clk856-GAL4 |
| DKI0082 | CG14871 | Trissin | LexA | Clk856-GAL4 |
| DKI0248 | CG9122 | Trh | LexA | Clk856-GAL4 |
| DKI0265 | CG9887 | VGlut | LexA | Clk856-GAL4 |
| DKI0165 | CG13586 | ITP | Gal4 | Clk856-p65AD |
| DKI0111 | CG10118 | TH | Gal4 | Clk856-p65AD |
| DKI0168 | CG3441 | Nplp1 | Gal4 | Clk856-p65AD |
| DKI0287 | CG14723 | HisCl1 | Gal4 | Clk856-p65AD |
| DKI0203 | CG5610 | nAChRα1 | Gal4 | Clk856-p65AD |
| WCKI1015 | CG10342 | NPF | LexA | Clk856-GAL4 |
| DKI0099 | CG14358 | CCHa1 | LexA | Clk856-GAL4 |
| WCKI1017 | CG33976 | Octβ2R | Gal4 | Clk856-p65AD |
| WCKI1019 | CG10698 | CrzR | Gal4 | Clk856-p65AD |
| DKI0078 | CG30106 | CCHa1-R | LexA | Clk856-GAL4 |
| DKI0029 | CG13936 | CNMa | p65AD | Clk856-GAL4 |
| DKI0288 | CG33696 | CNMaR | p65AD | Clk856-GAL4 |
| WCKI1095 | CG8348 | Dh44 | GAL4 | Clk856-p65AD |
| DKI0134 | CG13758 | Pdfr | LexA | Clk856-GAL4 |
| WCKI1199 | CG7485 | Oct-TyrR | Gal4 | Clk856-p65AD |
| DKI0262 | CG8216 | spab | LexA | Clk856-GAL4 |
| DKI0013 | CG10537 | Rdl | LexA | Clk856-GAL4 |
| DKI0069 | CG1056 | 5-HT2A | LexA | Clk856-GAL4 |
| WCKI1162 | CG10626 | Lkr | LexA | Clk856-GAL4 |
| DKI0251 | CG11318 | CG11318 | LexA | Clk856-GAL4 |
| WCKI1030 | CG11325 | AkhR | LexA | Clk856-GAL4 |
| WCKI1201 | CG1171 | Akh | LexA | Clk856-GAL4 |
| DKI0215 | CG11822 | nAChRβ3 | LexA | Clk856-GAL4 |
| CG11883-L-LexA | CG11883 | CG11883 | LexA | Clk856-GAL4 |
| CG11883-S-LexA | CG11883 | CG11883 | LexA | Clk856-GAL4 |
| WCKI1180 | CG11937 | amn | LexA | Clk856-GAL4 |
| DKI0307 | CG12344 | CG12344 | LexA | Clk856-GAL4 |
| Nplp3-LexA | CG13061 | Nplp3 | LexA | Clk856-GAL4 |
| L53-T5-W- | CG13480 | Lk | LexA | Clk856-GAL4 |
| DKI0302 | CG13565 | Orcokinin | LexA | Clk856-GAL4 |
| DKI0050 | CG13575 | CG13575 | LexA | Clk856-GAL4 |
| DKI0160 | CG13579 | CG13579 | LexA | Clk856-GAL4 |
| DKI0286 | CG13579 | CG13579-RB | LexA | Clk856-GAL4 |
| DKI0091 | CG13633 | AstA | LexA | Clk856-GAL4 |
| DKI0096 | CG14375 | CCHa2 | LexA | Clk856-GAL4 |
| DKI0086 | CG14575 | CapaR | LexA | Clk856-GAL4 |
| DKI0007 | CG14593 | CCHa2-R | LexA | Clk856-GAL4 |
| DKI0094 | CG14734 | Tk | LexA | Clk856-GAL4 |
| DKI0020 | CG14994 | gad1 | LexA | Clk856-GAL4 |
| DKI0089 | CG15113 | 5-HT1B | LexA | Clk856-GAL4 |
| DKI0187 | CG15274 | GABA-B-R1 | LexA | Clk856-GAL4 |
| DKI0055 | CG15284 | Pburs | LexA | Clk856-GAL4 |
| Tβh-LexA | CG1543 | Tβh | LexA | Clk856-GAL4 |
| DKI0143 | CG15520 | Capa | LexA | Clk856-GAL4 |
| WCKI1196 | CG15614 | CG15614 | LexA | Clk856-GAL4 |
| DKI0159 | CG15744 | CG15744 | LexA | Clk856-GAL4 |
| DKI0030 | CG16720 | 5-HT1A | LexA | Clk856-GAL4 |
| DKI0146 | CG16752 | SPR | LexA | Clk856-GAL4 |
| DKI0156 | CG16992 | mthl6 | LexA | Clk856-GAL4 |
| DKI0294 | CG17061 | mthl10 | LexA | Clk856-GAL4 |
| WCKI1120 | CG17084 | CG17084 | LexA | Clk856-GAL4 |
| DKI0031 | CG17795 | mthl2 | LexA | Clk856-GAL4 |
| DKI0196 | CG18039 | GluRIID | LexA | Clk856-GAL4 |
| DKI0148 | CG18090 | Dsk | LexA | Clk856-GAL4 |
| WCKI1097 | CG18208 | CG18208 | LexA | Clk856-GAL4 |
| DKI0321 | CG18314 | DopEcR | LexA | Clk856-GAL4 |
| DKI0123 | CG18741 | DopR2 | LexA | Clk856-GAL4 |
| DKI0205 | CG2302 | nAChR $\alpha 3$ | LexA | Clk856-GAL4 |
| WCKI1184 | CG2346 | FMRFa | LexA | Clk856-GAL4 |
| DKI0225 | CG2872 | AstA-R1 | LexA | Clk856-GAL4 |
| DKI0290 | CG30018 | mthl13 | LexA | Clk856-GAL4 |
| DKI0066 | CG30340 | CG30340 | LexA | Clk856-GAL4 |
| DKI0151 | CG31096 | Lgr3 | LexA | Clk856-GAL4 |
| DKI0231 | CG31147 | mthl11 | LexA | Clk856-GAL4 |
| DKI0232 | CG31760 | CG31760 | LexA | Clk856-GAL4 |
| DKI0234 | CG32447 | CG32447 | LexA | Clk856-GAL4 |
| DKI0259 | CG32475 | mthl8 | LexA | Clk856-GAL4 |
| WCKI1107 | CG32476 | CG32476 | LexA | Clk856-GAL4 |
| DKI0252 | CG32540 | CCKLR-17D3 | LexA | Clk856-GAL4 |
| DKI0206 | CG32975 | nAChR $\alpha 5$ | LexA | Clk856-GAL4 |
| DKI0101 | CG33344 | CCAP-R | LexA | Clk856-GAL4 |
| DKI0284 | CG33495 | Dup99B | LexA | Clk856-GAL4 |
| DKI0087 | CG33495 | Dup99B | LexA | Clk856-GAL4 |
| DKI0128 | CG33513 | Nmdar2 | LexA | Clk856-GAL4 |
| DKI0315 | CG33527 | SIFa | LexA | Clk856-GAL4 |
| DKI0180 | CG33639 | CG33639 | LexA | Clk856-GAL4 |
| DKI0246 | CG34388 | natalisin | LexA | Clk856-GAL4 |
| DKI0175 | CG34411 | CG34411 | LexA | Clk856-GAL4 |
| DKI0102 | CG3454 | HDC | LexA | Clk856-GAL4 |
| WCKI1101 | CG3856 | oamb | LexA | Clk856-GAL4 |
| DKI0208 | CG4128 | nAChR $\alpha 6$ | LexA | Clk856-GAL4 |
| WCKI1044 | CG42244 | Octβ3R | LexA | Clk856-GAL4 |
| DKI0193 | CG4226 | GluRIIC | LexA | Clk856-GAL4 |
| DKI0173 | CG42301 | CCKLR-17D1 | LexA | Clk856-GAL4 |
| WCKI1166 | CG4313 | CG4313 | LexA | Clk856-GAL4 |
| DKI0276 | CG4356 | mAChR-A | LexA | Clk856-GAL4 |
| DKI0016 | CG43745 | MsR2 | LexA | Clk856-GAL4 |
| WCKI1169 | CG4395 | hec | LexA | Clk856-GAL4 |
| NT5E1-lexA | CG4827 | NT5E-1 | LexA | Clk856-GAL4 |
| WCKI1176 | CG5400 | Eh | LexA | Clk856-GAL4 |
| WCKI1203 | CG5811 | RYa-R | LexA | Clk856-GAL4 |
| WCKI1205 | CG5911 | ETHR | LexA | Clk856-GAL4 |
| WCKI1055 | CG6371 | CG6371 | LexA | Clk856-GAL4 |
| WCKI1009 | CG6440 | Ms | LexA | Clk856-GAL4 |
| WCKI1011 | CG6456 | Mip | LexA | Clk856-GAL4 |
| WCKI1192 | CG6530 | mthl3 | LexA | Clk856-GAL4 |
| WCKI1194 | CG6536 | mthl4 | LexA | Clk856-GAL4 |
| WCKI1049 | CG6919 | oa2 | LexA | Clk856-GAL4 |
| WCKI1118 | CG6965 | CG6965 | LexA | Clk856-GAL4 |
| DKI0192 | CG6992 | GluRIIA | LexA | Clk856-GAL4 |
| WCKI1093 | CG7285 | AstC-R1 | LexA | Clk856-GAL4 |
| DKI0071 | CG7411 | Ort | LexA | Clk856-GAL4 |
| DKI0311 | CG7431 | TyrR | LexA | Clk856-GAL4 |
| DKI0299 | CG7446 | Grd | LexA | Clk856-GAL4 |
| DKI0320 | CG7497 | CG7497 | LexA | Clk856-GAL4 |
| WCKI1094 | CG7665 | Lgr1 | LexA | Clk856-GAL4 |
| DKI0240 | CG8380 | DAT | LexA | Clk856-GAL4 |
| DKI0047 | CG8394 | vGAT | LexA | Clk856-GAL4 |
| WCKI1181 | CG8422 | Dh44-R1 | LexA | Clk856-GAL4 |
| DKI0269 | CG8442 | GluRIA | LexA | Clk856-GAL4 |
| WCKI1174 | CG8795 | PK2-R2 | LexA | Clk856-GAL4 |
| WCKI1060 | CG10001 | CG10001 | Gal4 | Clk856-p65AD |
| DKI0130 | CG14723 | HisCl1 | Gal4 | Clk856-p65AD |
| DKI0041 | CG15361 | Nplp4 | Gal4 | Clk856-p65AD |
| WCKI1198 | CG15556 | CG15556 | Gal4 | Clk856-p65AD |
| DKI0255 | CG18105 | ETH | Gal4 | Clk856-p65AD |
| DKI0198 | CG31201 | GluRIIE | Gal4 | Clk856-p65AD |
| WCKI1164 | CG3171 | Tre1 | Gal4 | Clk856-p65AD |
| DKI0300 | CG4910 | CCAP | Gal4 | Clk856-p65AD |
| DKI0226 | CG6936 | mth | Gal4 | Clk856-p65AD |
| WCKI1178 | CG6986 | Proc-R | Gal4 | Clk856-p65AD |
| DKI0297 | CG7476 | mthl7 | Gal4 | Clk856-p65AD |
| WCKI1053 | CG9918 | PK1-R | Gal4 | Clk856-p65AD |

**Table S6.**
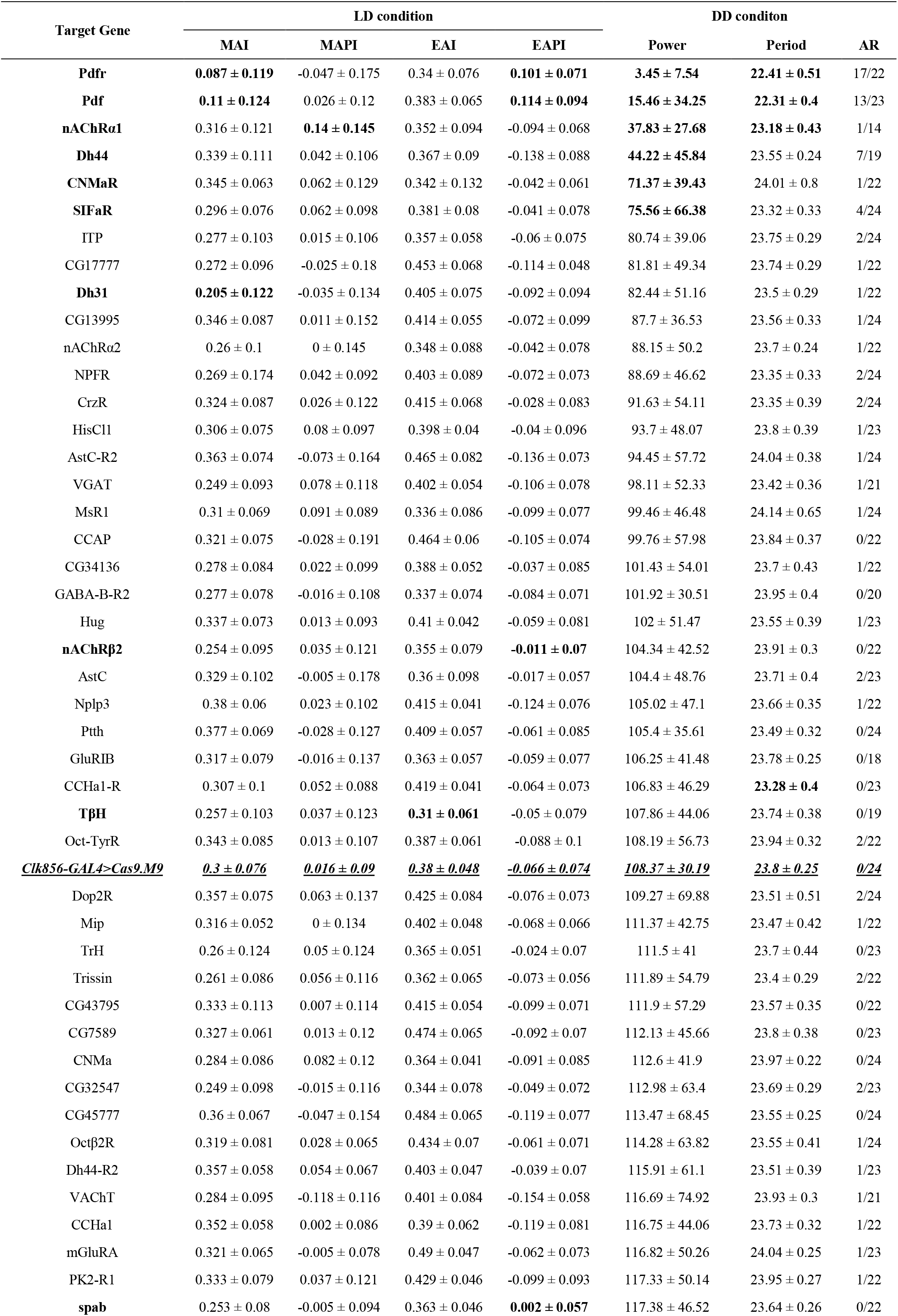
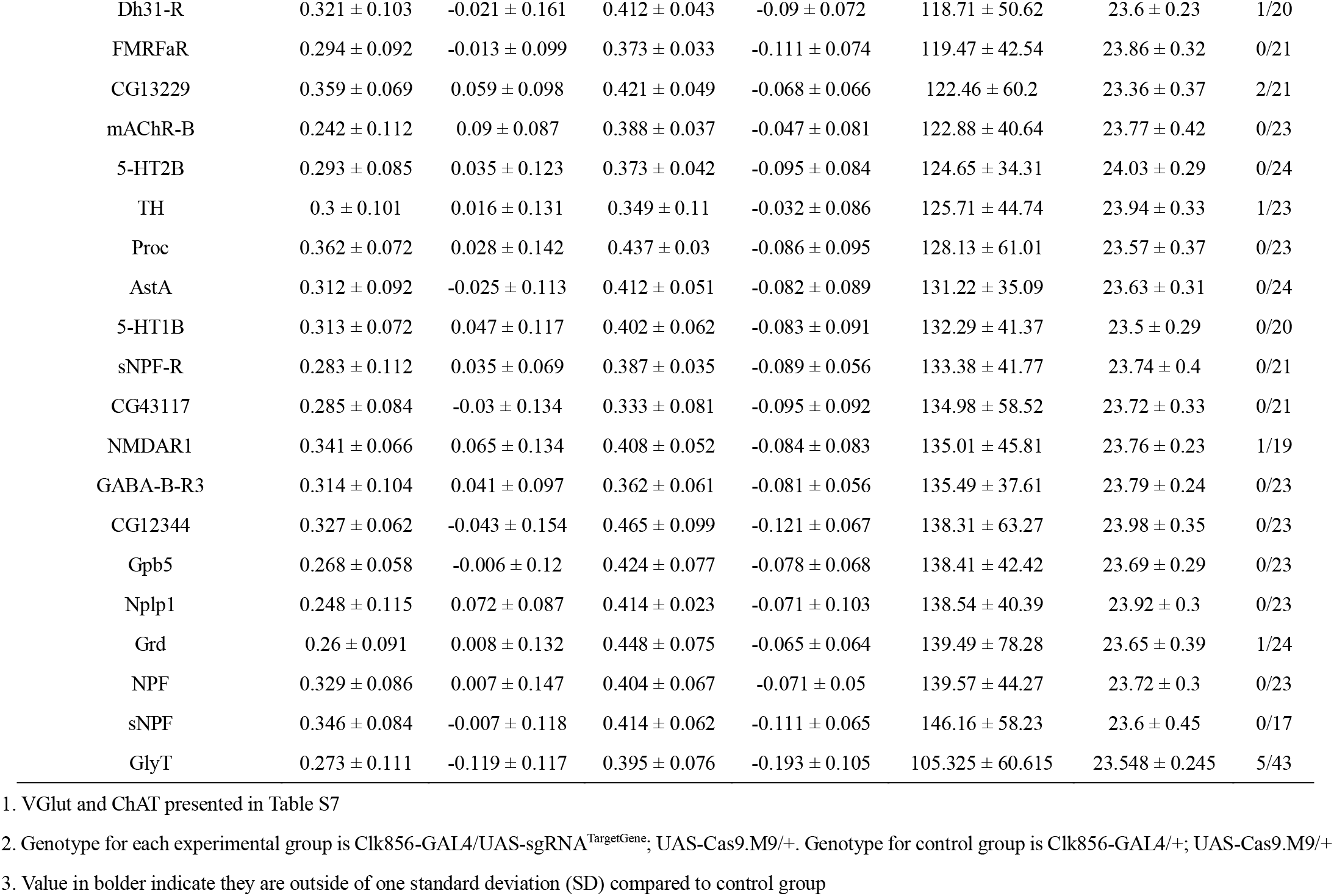
Phenotypes of CCT genes knocking out in clock neurons.

**Table S7.**
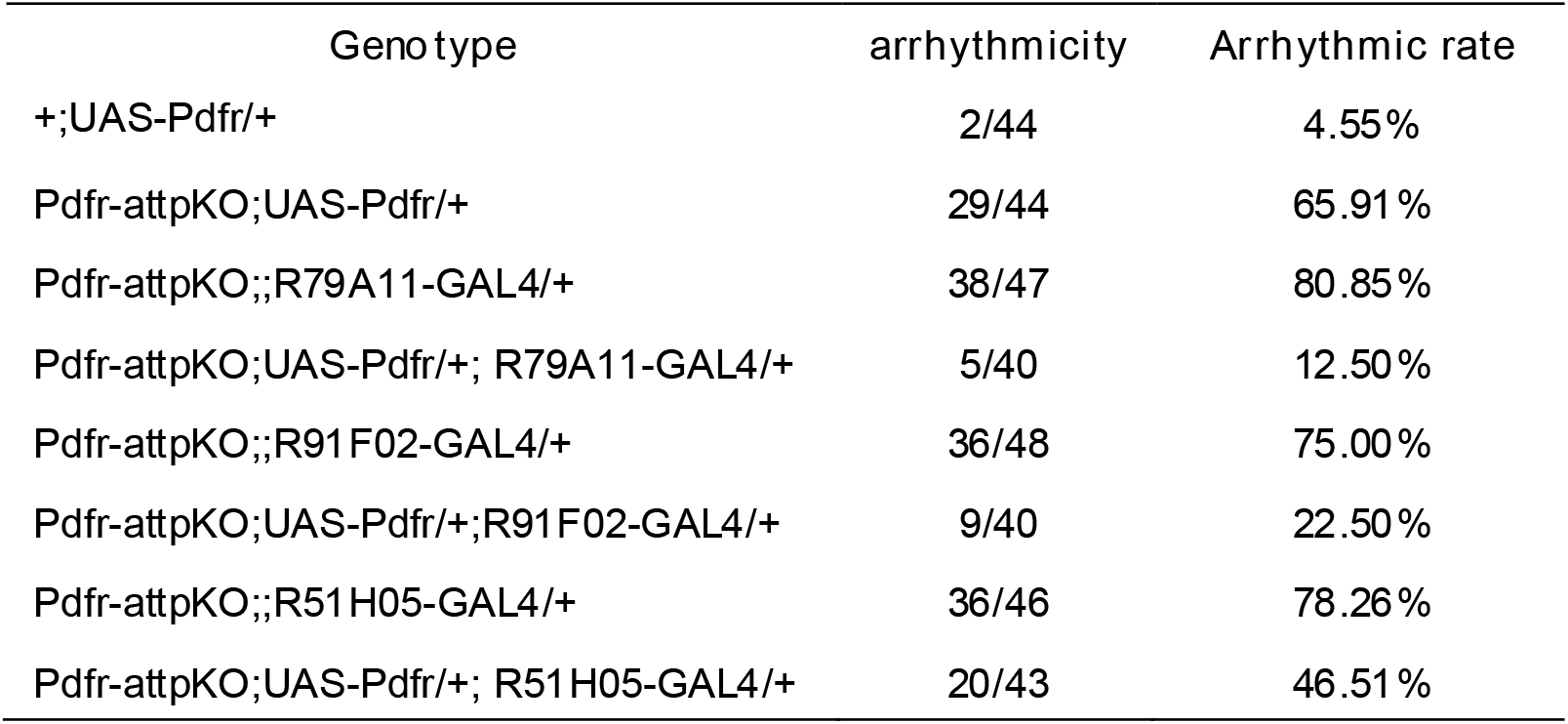
arrhythmicity related to Figure 6R.

**Table S7.** Phenotypes of candidate CCT genes knockout in clock neurons.

| Genotype | LD condition |  |  |  | DD condition |  |  |
| --- | --- | --- | --- | --- | --- | --- | --- |
|  | MAI | MAPI | EAI | EAPI | Power | Period | AR |
| sgRNA<sup>Dh44</sup>, Cas9.M6 | 0.329 ± 0.086 | -0.058 ± 0.119 | 0.38 ± 0.058 | -0.081 ± 0.065 | 66.38 ± 40.4 | 23.18 ± 0.31 | 2/22 |
| Clk856> sgRNA<sup>Dh44</sup> | 0.354 ± 0.093 | 0.03 ± 0.131 | 0.444 ± 0.045 | -0.085 ± 0.066 | 94 ± 45.35 | 23.61 ± 0.34 | 2/20 |
| Clk856> sgRNA<sup>Dh44</sup>, Cas9.M6 | 0.346 ± 0.096 | -0.02 ± 0.165 | 0.397 ± 0.059 | -0.085 ± 0.077 | 74.64 ± 27.25 | 23.75 ± 0.31 | 0/20 |
| sgRNA<sup>nAChRα1</sup>, Cas9.M6 | 0.29 ± 0.089 | -0.064 ± 0.139 | 0.363 ± 0.056 | -0.071 ± 0.064 | 58.35 ± 43.82 | 23.53 ± 0.45 | 2/13 |
| Clk856> sgRNA<sup>nAChRα1</sup> | 0.318 ± 0.096 | 0.019 ± 0.137 | 0.401 ± 0.052 | -0.08 ± 0.071 | 56.89 ± 45.03 | 23.66 ± 0.43 | 1/12 |
| Clk856> sgRNA<sup>nAChRα1</sup>, Cas9.M6 | 0.307 ± 0.154 | -0.031 ± 0.099 | 0.381 ± 0.056 | -0.074 ± 0.074 | 69.21 ± 52.1 | 23.71 ± 0.29 | 2/13 |
| sgRNA<sup>ChAT</sup>, Cas9.M6 | 0.292 ± 0.096 | 0.026 ± 0.155 | 0.291 ± 0.092 | -0.1 ± 0.055 | 91.48 ± 55.18 | 23.43 ± 0.41 | 3/19 |
| Clk856> sgRNA<sup>ChAT</sup> | 0.298 ± 0.098 | -0.016 ± 0.157 | 0.353 ± 0.071 | -0.079 ± 0.051 | 78.66 ± 41.65 | 23.54 ± 0.25 | 1/22 |
| Clk856> sgRNA<sup>ChAT</sup>, Cas9.M6 | 0.22 ± 0.137 | 0.003 ± 0.123 | 0.318 ± 0.078 | -0.124 ± 0.075 | 49.35 ± 42.11 | 23.7 ± 0.2 | 4/14 |
| sgRNA<sup>CNMa</sup>, Cas9.M6 | 0.342 ± 0.076 | -0.141 ± 0.138 | 0.35 ± 0.091 | -0.088 ± 0.079 | 92.81 ± 39.89 | 23.98 ± 0.44 | 1/23 |
| Clk856> sgRNA<sup>CNMa</sup> | 0.355 ± 0.181 | 0.019 ± 0.093 | 0.431 ± 0.05 | -0.121 ± 0.079 | 18.96 ± 22.19 | 23.8 ± 0.27 | 7/14 |
| Clk856> sgRNA<sup>CNMa</sup>, Cas9.M6 | 0.354 ± 0.11 | 0.113 ± 0.169 | 0.353 ± 0.086 | -0.072 ± 0.066 | 93.51 ± 48.39 | 24.17 ± 0.42 | 1/20 |
| sgRNA<sup>VGlut</sup>, Cas9.M6 | 0.254 ± 0.114 | -0.115 ± 0.14 | 0.32 ± 0.066 | -0.073 ± 0.043 | 56.28 ± 39.85 | 23.41 ± 0.38 | 1/20 |
| Clk856> sgRNA<sup>VGlut</sup> | 0.306 ± 0.102 | -0.026 ± 0.146 | 0.369 ± 0.074 | -0.104 ± 0.067 | 63.82 ± 48.6 | 23.6 ± 0.36 | 5/20 |
| Clk856> sgRNA<sup>VGlut</sup>, Cas9.M6 | 0.179 ± 0.1** | -0.087 ± 0.107 | 0.27 ± 0.07 | -0.158 ± 0.063 | 45.09 ± 41.35 | 23.7 ± 0.27 | 6/21 |
| sgRNA<sup>mAChR-B</sup>, Cas9.M6 | 0.37 ± 0.062 | -0.047 ± 0.109 | 0.374 ± 0.064 | -0.076 ± 0.054 | 110.16 ± 32.99 | 23.78 ± 0.47 | 0/21 |
| Clk856> sgRNA<sup>mAChR-B</sup> | 0.358 ± 0.095 | -0.003 ± 0.109 | 0.442 ± 0.038 | -0.143 ± 0.087 | 73 ± 36 | 23.48 ± 0.45 | 0/17 |
| Clk856> sgRNA<sup>mAChR-B</sup>, Cas9.M6 | 0.259 ± 0.1 | -0.109 ± 0.139 | 0.357 ± 0.066 | -0.088 ± 0.074 | 109.35 ± 45.07 | 23.71 ± 0.36 | 0/22 |
| sgRNA<sup>MsR1</sup>, Cas9.M6 | 0.339 ± 0.078 | -0.011 ± 0.175 | 0.368 ± 0.061 | -0.072 ± 0.087 | 55.68 ± 49.3 | 23.77 ± 0.55 | 5/22 |
| Clk856> sgRNA<sup>MsR1</sup> | 0.378 ± 0.098 | -0.023 ± 0.114 | 0.4 ± 0.036 | -0.143 ± 0.075 | 55.05 ± 43.19 | 23.89 ± 0.4 | 3/19 |
| Clk856> sgRNA<sup>MsR1</sup>, Cas9.M6 | 0.276 ± 0.085 | -0.04 ± 0.123 | 0.313 ± 0.063 | -0.085 ± 0.083 | 78.21 ± 50.06 | 23.85 ± 0.48 | 2/21 |
| sgRNA<sup>SIFaR</sup>, Cas9.M6 | 0.254 ± 0.104 | -0.075 ± 0.14 | 0.34 ± 0.047 | -0.072 ± 0.052 | 82.85 ± 42.29 | 23.63 ± 0.35 | 0/23 |
| Clk856> sgRNA<sup>SIFaR</sup> | 0.329 ± 0.122 | -0.058 ± 0.121 | 0.375 ± 0.059 | -0.096 ± 0.05 | 81.71 ± 41.2 | 23.82 ± 0.36 | 0/21 |
| Clk856> sgRNA<sup>SIFaR</sup>, Cas9.M6 | 0.219 ± 0.143 | -0.142 ± 0.172 | 0.299 ± 0.095 | -0.117 ± 0.088 | 94.45 ± 32.78 | 24.04 ± 0.38 | 0/19 |
| Clk856> Cas9.M6 | 0.295 ± 0.12 | -0.043 ± 0.109 | 0.325 ± 0.081 | -0.118 ± 0.048 | 90.64 ± 43.86 | 23.7 ± 0.27 | 1/16 |

Clk856> sgRNA<sup>VGlut</sup>, Cas9.M6 vs Clk856> sgRNA<sup>VGlut</sup>, \*\* $P < 0.01$
Clk856> sgRNA<sup>VGlut</sup>, Cas9.M6 vs Clk856> Cas9.M6, \*\* $P < 0.01$
Clk856> sgRNA<sup>VGlut</sup>, Cas9.M6 vs sgRNA<sup>VGlut</sup>, Cas9.M6, $P = 0.0921$

**Table S8.** Conditional knockout of VGlut in DN1s.

| Genotype | LD condition |  |  |  | DD condition |  |  |
| --- | --- | --- | --- | --- | --- | --- | --- |
|  | MAI | MAPI | EAI | EAPI | POWER | PERIOD | AR |
| sgRNA<sup>VGlut</sup>,Cas9.M6 | 0.268 ± 0.124 | -0.072 ± 0.106 | 0.328 ± 0.071 | -0.065 ± 0.035 | 79.06 ± 49.45 | 22.89 ± 0.19 | 1/23 |
| R18H11>sgRNA<sup>VGlut</sup> | 0.258 ± 0.085 | -0.093 ± 0.104 | 0.327 ± 0.05 | -0.124 ± 0.069 | 67.49 ± 42.52 | 23.38 ± 0.36 | 1/21 |
| R18H11>sgRNA<sup>VGlut</sup>,Cas9.M6 | **0.136 ± 0.148**<br>** | -0.109 ± 0.116 | 0.284 ± 0.106 | -0.084 ± 0.061 | 61.9 ± 42.35 | 23.18 ± 0.27 | 4/24 |
| R18H11>Cas9.M6 | 0.25 ± 0.093 | -0.068 ± 0.124 | 0.387 ± 0.055 | -0.048 ± 0.057 | 55.46 ± 49.14 | 23.33 ± 0.43 | 5/20 |
| R51H05>sgRNA<sup>VGlut</sup> | 0.348 ± 0.067 | -0.078 ± 0.078 | 0.398 ± 0.051 | -0.103 ± 0.047 | 122.02 ± 38.89 | 23.4 ± 0.37 | 0/20 |
| R51H05>sgRNA<sup>VGlut</sup>,Cas9.M6 | 0.26 ± 0.097 | -0.16 ± 0.116 | 0.312 ± 0.073 | -0.062 ± 0.059 | 82.56 ± 48.06 | 23.23 ± 0.33 | 2/22 |
| R51H05>Cas9.M6 | 0.251 ± 0.098 | -0.118 ± 0.143 | 0.349 ± 0.087 | -0.055 ± 0.072 | 45.76 ± 41.2 | 23.19 ± 0.39 | 3/17 |
| R79A11>sgRNA<sup>VGlut</sup> | 0.172 ± 0.131 | -0.11 ± 0.102 | 0.378 ± 0.055 | -0.087 ± 0.062 | 88.87 ± 40.43 | 23.43 ± 0.31 | 1/21 |
| R79A11>sgRNA<sup>VGlut</sup>,Cas9.M6 | 0.178 ± 0.11 | -0.131 ± 0.083 | 0.325 ± 0.08 | -0.068 ± 0.037 | 75.67 ± 45.37 | 23.3 ± 0.29 | 1/19 |
| R79A11>Cas9.M6 | 0.19 ± 0.132 | -0.128 ± 0.114 | 0.346 ± 0.052 | -0.082 ± 0.065 | 31.29 ± 32.62 | 23.16 ± 0.47 | 3/21 |
| R91F02>sgRNA<sup>VGlut</sup> | 0.233 ± 0.099 | -0.108 ± 0.1 | 0.341 ± 0.069 | -0.081 ± 0.049 | 97.13 ± 41.94 | 23.73 ± 0.39 | 0/21 |
| R91F02>sgRNA<sup>VGlut</sup>,Cas9.M6 | 0.301 ± 0.109 | -0.109 ± 0.056 | 0.381 ± 0.071 | -0.061 ± 0.042 | 112.28 ± 47.81 | 23.45 ± 0.31 | 0/24 |
| R91F02>Cas9.M6 | 0.26 ± 0.166 | -0.078 ± 0.107 | 0.366 ± 0.065 | -0.035 ± 0.04 | 72.82 ± 45.83 | 23.43 ± 0.41 | 2/22 |
| CNMa-KI-GAL4>sgRNA<sup>VGlut</sup> | 0.28 ± 0.123 | -0.058 ± 0.088 | 0.357 ± 0.043 | -0.063 ± 0.019 | 54.7 ± 43.96 | 23.03 ± 0.33 | 3/23 |
| CNMa-KI-GAL4>sgRNA<sup>VGlut</sup>,Cas9.M6 | 0.268 ± 0.094 | -0.117 ± 0.119 | 0.331 ± 0.055 | -0.048 ± 0.086 | 51.21 ± 36.91 | 22.83 ± 0.29 | 3/18 |
| CNMa-KI-GAL4>Cas9.M6 | 0.261 ± 0.066 | -0.155 ± 0.114 | 0.342 ± 0.051 | -0.088 ± 0.08 | 54.14 ± 36.21 | 22.85 ± 0.21 | 2/24 |
| Clk4.1M>sgRNA<sup>VGlut</sup> | 0.281 ± 0.116 | -0.052 ± 0.102 | 0.4 ± 0.057 | -0.174 ± 0.043 | 61.29 ± 46.5 | 23.45 ± 0.39 | 3/23 |
| Clk4.1M>sgRNA<sup>VGlut</sup>,Cas9.M6 | 0.181 ± 0.111 | -0.074 ± 0.07 | 0.313 ± 0.101 | -0.112 ± 0.047 | 77.13 ± 38.86 | 23.44 ± 0.38 | 0/21 |
| Clk4.1M>Cas9.M6 | 0.294 ± 0.102 | -0.096 ± 0.098 | 0.41 ± 0.035 | -0.087 ± 0.051 | 80.18 ± 40.76 | 23.51 ± 0.25 | 0/24 |

\*\* $P < 0.01$

